# Transcriptomic Signature and PROTAC Strategy Revealed Histone Lysine Demethylase as a Target of Anticancer Activity of Deferiprone

**DOI:** 10.1101/2025.10.13.682199

**Authors:** Alexis Johnston, Jeremiah O. Olugbami, Dipak Walunj, Arvind Bangaru, Bocheng Wu, Ryan Kern, Ruiqiao Yang, Travis J. Nelson, Brandon J. Clarke, Janani Murugan, Nathaniel A Hathaway, Yuhong Fan, Adegboyega K. Oyelere

## Abstract

Deferiprone (DFP) is an iron chelator approved for treating iron overload in thalassemia patients. Recent observations have suggested that DFP has promising anticancer activities ascribed to several mechanisms including reduction of the intracellular free labile iron and zinc ion pools and inhibition of the activities of other intracellular targets, including ribonucleotide reductase (RNR). We previously reported that DFP inhibits the demethylase activities of several Fe(II)/α- ketoglutarate dependent histone lysine demethylases (KDMs) at much lower concentrations at which it inhibits RNR activities and/or reduces the labile intracellular iron and zinc ion pools. In this study, we used RNA sequencing (RNA seq) and PROTACs strategies to validate and quantify the contribution of intracellular KDM inhibition to the antiproliferative activities of DFP. We report herein that DFP elicited gene expression signature that is largely similar to that of JIB-04, an established KDM inhibitor (KDMi), in two breast cancer (BCa) cells (MCF-7 and MDA-MD-231). Importantly, RNA seq revealed that DFP and JIB-04 downregulated the expression of hypoxia-inducible factor 1α (HIF-1α), an oncogene whose expression is commonly modulated through histone demethylation mediated by KDMs and degraded by several KDMi.

Moreover, DFP-derived PROTACs elicited enhanced cancer cell selective antiproliferative activities and intracellular on-target effects, downregulating several KDMs implicated in the etiology of BCa cells, including a strong degradation of KDMs 2A, 3A and 5B, and a moderate degradation of KDMs 4A-C, 5C, 6B. Collectively, our data supports KDM inhibition as a key mechanism of anticancer activity of DFP and identifies PROTAC is a viable strategy to obtain novel DFP analogs with improved potency and therapeutic index.

## Introduction

Deferiprone (DFP) (Figure 1a), an FDA-approved iron chelator originally used for treating iron overload in thalassemia patients,^1^ has emerged as a promising candidate for the development of novel therapies targeting breast cancer (BCa). We previously demonstrated that DFP functions as a pan-selective histone lysine demethylase (KDM) inhibitor (KDMi), inhibiting KDMs that are implicated in BCa etiology.

**Figure 1.**
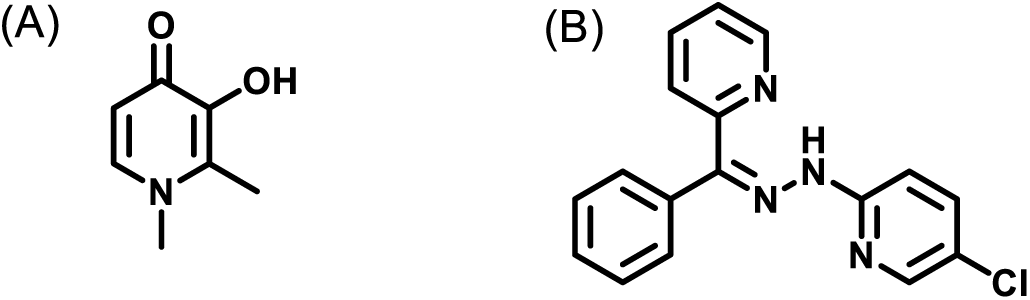
Structures of DFP (A) and JIB-04 (B).

In a cell-free assay, DFP inhibits a cohort of Fe(II)/α-ketoglutarate dependent KDMs with low micromolar IC_50_s^2^ that are several folds lower than the concentrations at which DFP reduces intracellular labile iron or zinc ion pools or inhibits the activities of other intracellular targets, such as ribonucleotide reductase (RNR), which have been attributed to be responsible for the antiproliferative effects of DFP.^3–7^ Intriguingly, the effects of DFP on KDM inhibition in a cell-based assay^8^ and on the proliferation of representative breast cancer (BCa) cells (MCF-7 and MDA-MB-231) are not as pronounced as its cell-free inhibition effects of KDMs. DFP inhibits the proliferation of MCF-7 and MDA-MB-231 activities and induces global histone protein hypermethylation at concentration higher than 100 μM. Subsequent structure-based optimization resulted in DPF-based KDMi with enhanced KDM inhibition and antiproliferative activities against MCF-7 and MDA-MB-231. A cohort of these compounds are preferentially cytotoxic to the triple-negative breast cancer (TNBC) cell line MDA-MB-231 while a representative lead compound effectively reduced tumor growth in murine models of ER+ and ER- BCas.^2^

The goal of this study is to validate and quantify the contribution of intracellular KDM inhibition to the antiproliferative activities of DFP and its analogs. Using RNA sequencing (RNA-seq), we quantitatively studied the effects of DFP on the transcriptome, first focusing on specific genes that have been described as drivers of the antiproliferative activities of JIB-04 (Figure 1b), a pan-selective KDMi.^9^ We observed that DFP perturbs the expression pattern of these genes in MCF-7 and MDA-MB-231 cells in a similar manner to JIB-04. In depth analysis of the RNA-seq data revealed that DFP perturbs similar cancer relevant pathways as JIB-04.

Subsequently, we designed PROTAC analogs of DFP and observed that these PROTACs effectively inhibit histone lysine demethylation in cell-based assay and also inhibit the proliferation of several cancer cell lines with IC_50_s more than 260-fold higher than DFP. Furthermore, a representative DFP-derived PROTAC strongly degrades KDMs 2A, 3A and 5B, and moderately degrades KDMs 4A-C, 5C, and 6B.

Collectively, we present evidence supporting KDM inhibition as a key mechanism of anticancer activity of DFP. Moreover, our study furnished novel, highly potent DFP-derived PROTACs whose anticancer activities merit further investigation.

## Results

### RNA seq analysis revealed DFP elicits KDM inhibition activity

JIB-04, like DFP, is a previously reported pan-selective KDM inhibitor that also impair tumor cell survival in hepatocellular carcinoma cells, Ewing Sarcoma, and colorectal cancer cells.^9–11^ Comparison of the effects of JIB-04 and DFP on the transcriptome could help validate the contribution of intracellular KDM inhibition to the antiproliferative activities of DFP. Given the difference in their target KDMs,^2, 9–12^ such analysis could also reveal disparities in the cellular pathways perturbed by JIB-04 and DFP. Therefore, we first analyzed the effects of JIB-04 and DFP at IC_50_ and 2x IC_50_ on the expression status of cohort of genes, proposed to mediate JIB-04 inhibition in Ewing Sarcoma,^9^ in a TNBC (MDA-MB-231) and ER+ (MCF-7) BCa cell lines treated with each compound for 24 h. The effect of each compound, relative to the DMSO-treated control, on RNA expression levels in each cell is represented as log2 fold change in Figure 2 and Supplemental Figure S1. Similar to JIB-04, the most significantly perturbed genes by DFP are tumor suppressor CDKN1A/p21 and oncogenes AURKA and CCNB1 in MDA-MB-231 cells (Figure 2a and Supplemental Table S1). CDKN1A was heavily upregulated by DFP (+2.1, 2x IC_50_) and JIB-04 (+2.4, 2x IC_50_) while AURKA and CCNB1 were downregulated by DFP (−1.3 and −1.6, 2x IC_50_) and JIB-04 (−1.6 and −1.9, 2x IC_50_). Analysis of treatment to MCF-7 cells also revealed CDKN1A upregulation by DFP and JIB-04 (+1.3, 2x IC_50_ for both treatments) and downregulation of CCNB1, AURKA, and AURKB by DFP (−2.3, −1.7, and −1.2, 2x IC_50_) and JIB-04 (−1.2, -0.7, and −1.0, 2x IC_50_) (Figure 2c and Supplemental Table S1). Additionally, tumor suppressor PTEN was upregulated by JIB-04 and DFP (+1.0, +1.3, 2x IC_50_). JIB-04 upregulated cell cycle inhibitor CDKN2D and pro-apoptotic factor BCL10 (+1.5 and +1.6, 2x IC_50_) and downregulated cell cycle promoters AKT1 and CDK5, and oncogene EPHB4 (−1.5, −1.3 and −1.0, 2x IC_50_). The fold-change gene in expression levels induced by JIB-04 treatment compared to DFP at 2x IC_50_, ranked from highest to least expression levels, revealed the similarity between these two small molecules in each respective cell line. In MDA-MB-231 cells, the only exceptions are the differing expression levels of pro-apoptotic factor BCL2L11 and oncogene PDGFRB (Figure 2b). In MCF-7, CDKN1C, FOXO4, EPHB4, and AKT1 were downregulated by JIB-04 and upregulated by DFP while APC was downregulated by DFP and upregulated by JIB-04 (Figure 2c and Supplemental Table S1).

**Figure 2.**
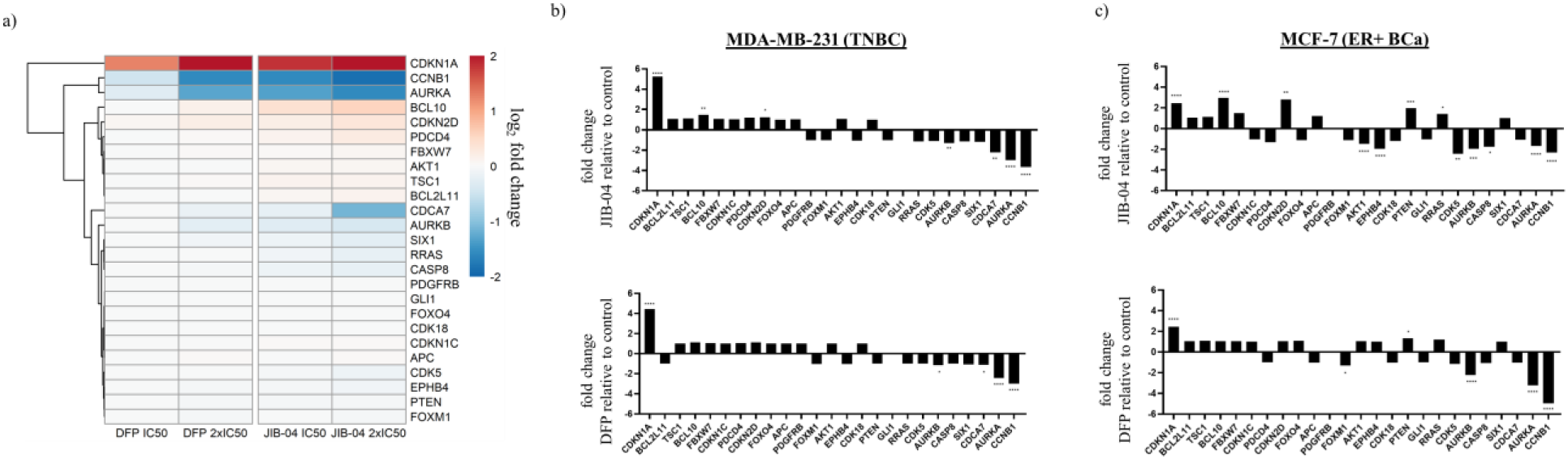
Effects of DFP on genes implicated in KDM inhibition activity of JIB-04. a) Log2 fold change heatmap of genes implicated in KDM inhibition by JIB-04^9^ displaying upregulation of CDKN1A (p21) by DFP 2x IC_50_ (log2 fold change = +2.1) and JIB-04 2x IC_50_ (log2 fold change = +2.4), downregulation of AURKA (DFP 2x IC_50_ log2 fold change = −1.3, JIB-04 2x IC_50_ log2 fold change = −1.6), and downregulation of CCNB1 (DFP 2x IC_50_ log2 fold change = - 1.6, JIB-04 1µM log2 fold change = −1.9) in MDA-MB-231 cells. Fold change bar graphs of DFP 2x IC_50_ and JIB-04 2x IC_50_ relative to DMSO in MDA-MB-231 (b) and MCF-7 (c) cells. Gene names arranged in descending order of expression levels resulting from JIB-04 treatment to MDA-MB-231 cells. * p < 0.05, ** p < 0.01, *** p < 0.001, **** p < 0.0001.

### DFP selectively enriched hallmark and gene ontology biological processes gene sets

Subsequently, we performed a hallmark gene set enrichment analysis (GSEA) of differentially expressed genes (DEGs) in MDA-MB-231 and MCF-7 cells treated with DFP and JIB-04, relative to the DMSO-treated cells control. Normalized enrichment scores (NES) were used to compare GSEA results across samples, and significantly enriched gene sets defined by a p-value < 0.05 and false discovery rate (FDR) < 0.25 were analyzed. DFP (2x IC_50_) and JIB-04 (IC_50_ and 2x IC_50_) significantly enriched hallmark gene sets (Figure 3) in both MCF-7 and MDA-MB-231 cells lines. DFP at 2x IC_50_ negatively enriched G2M checkpoint (NES = −3.8, MCF-7, and −2.1, MDA-MB-231), mitotic spindle (NES = −2.7, MCF-7, and −1.5, MDA-MB-231), estrogen response late (NES = −2.3, MCF-7) and positively enriched the hypoxia (NES = +3.1, MCF-7, and +2.3, MDA-MB-231) gene sets. The hallmark gene sets negatively enriched by JIB-04 at 2x IC_50_ include G2M checkpoint (NES = −2.8, MDA-MB-231), mitotic spindle (NES = −2.1, MDA-MB-231) and E2F targets (NES = −1.8, MCF-7, and −2.3, MDA-MB-231) while it positively enriched hypoxia (NES = +4.2, MCF-7, and +2.8, MDA-MB-231). Gene sets identified as significantly impacted by DFP and JIB-04 through Gene Ontology Biological Process (GOBP) analysis were predominantly negatively enriched in MDA-MB-231 cells. In MCF-7 cells, GOBP gene sets were similarly negatively enriched while JIB-04 positively enriched several gene sets. Significant gene sets were determined using the same parameters from the hallmark GSEA (p < 0.05, FDR < 0.25). Cell division, chromosome organization, and microtubule and spindle organization gene sets were negatively enriched by DFP (2x IC_50_) in both cell lines and by JIB- 04 (IC_50_ and 2x IC_50_) in MDA-MB-231 cells (Figure 4). Due to no significant enrichment by DFP at IC_50_ in either cell line, comparison between DFP and JIB-04 at 2x IC_50_ was prioritized.

**Figure 3.**
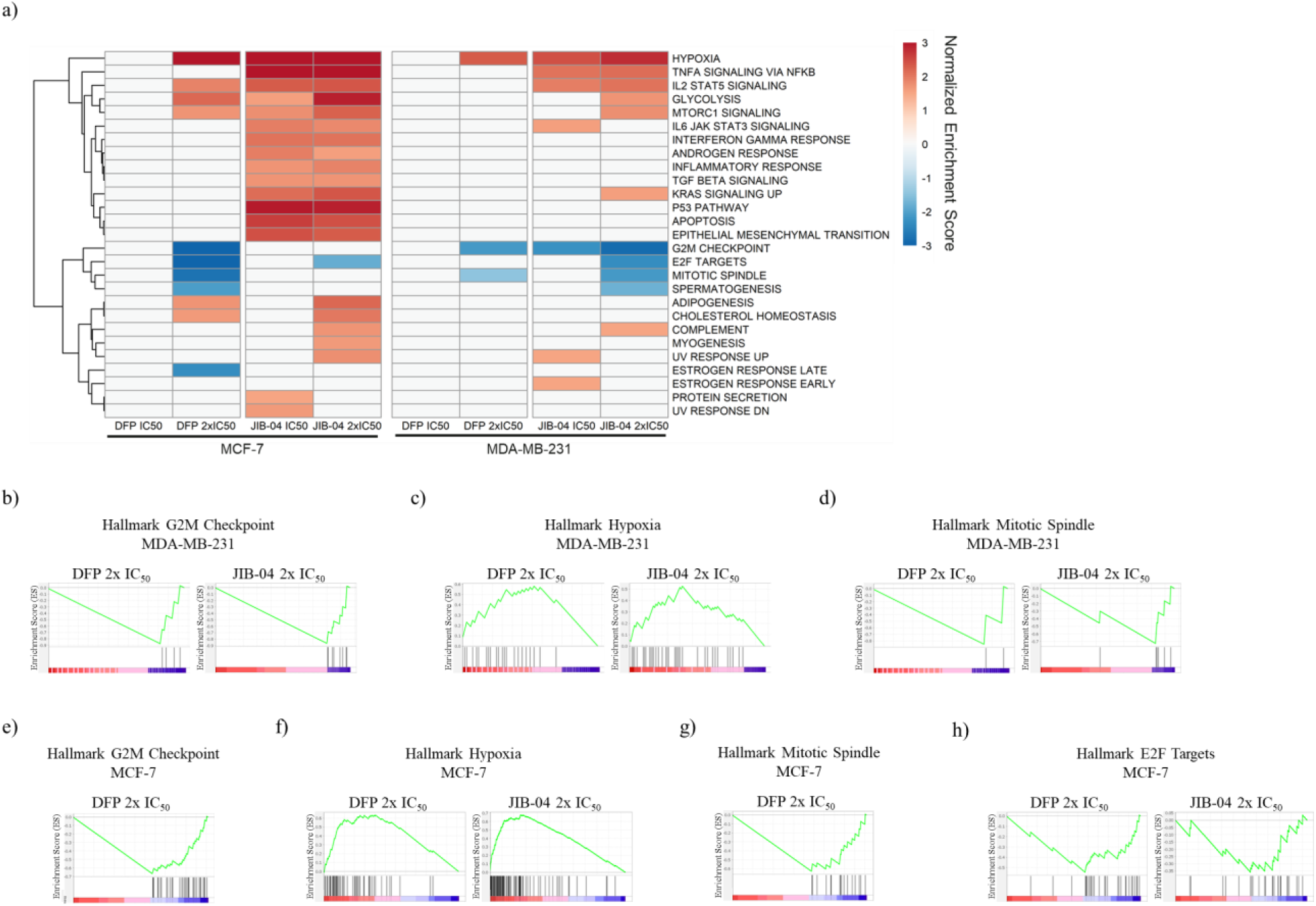
Hallmark Gene Set Enrichment Analysis (GSEA) in MCF-7 and MDA-MB-231 cells treated with DFP and JIB-04. a) Heatmap of normalized enrichment scores (NES) of significantly enriched hallmark gene sets in MCF-7 and MDA-MB-231 cells treated with DFP 2x IC_50_, JIB-04 IC_50_, and JIB-04 2x IC_50_ (p < 0.05, FDR < 0.25). Note that DFP at IC_50_ did not significantly enrich hallmark gene sets. b) G2M checkpoint enrichment plot for DFP 2x IC_50_ (NES = −2.1) and JIB-04 2x IC_50_ (NES = −2.8) in MDA-MB-231 cells. c) Mitotic spindle enrichment plot for DFP 2x IC_50_ (NES = −1.5) and JIB-04 2x IC_50_ (NES = −2.1) in MDA-MB- 231 cells. d) Hypoxia enrichment plot for DFP 2x IC_50_ (NES = +2.3) and JIB-04 2x IC_50_ (NES = +2.8) in MDA-MB-231 cells. e) G2M checkpoint enrichment plot for DFP 2x IC_50_ (NES = −3.8) in MCF-7 cells. f) Hypoxia enrichment plot for DFP 2x IC_50_ (NES = +3.1) and JIB-04 2x IC_50_ (NES = +4.2) in MCF-7 cells. g) Mitotic spindle enrichment plot for DFP 2x IC_50_ (NES = −2.7) in MCF-7 cells. h) E2F targets for DFP 2x IC_50_ (NES = −2.9) in MCF-7 cells.

**Figure 4.**
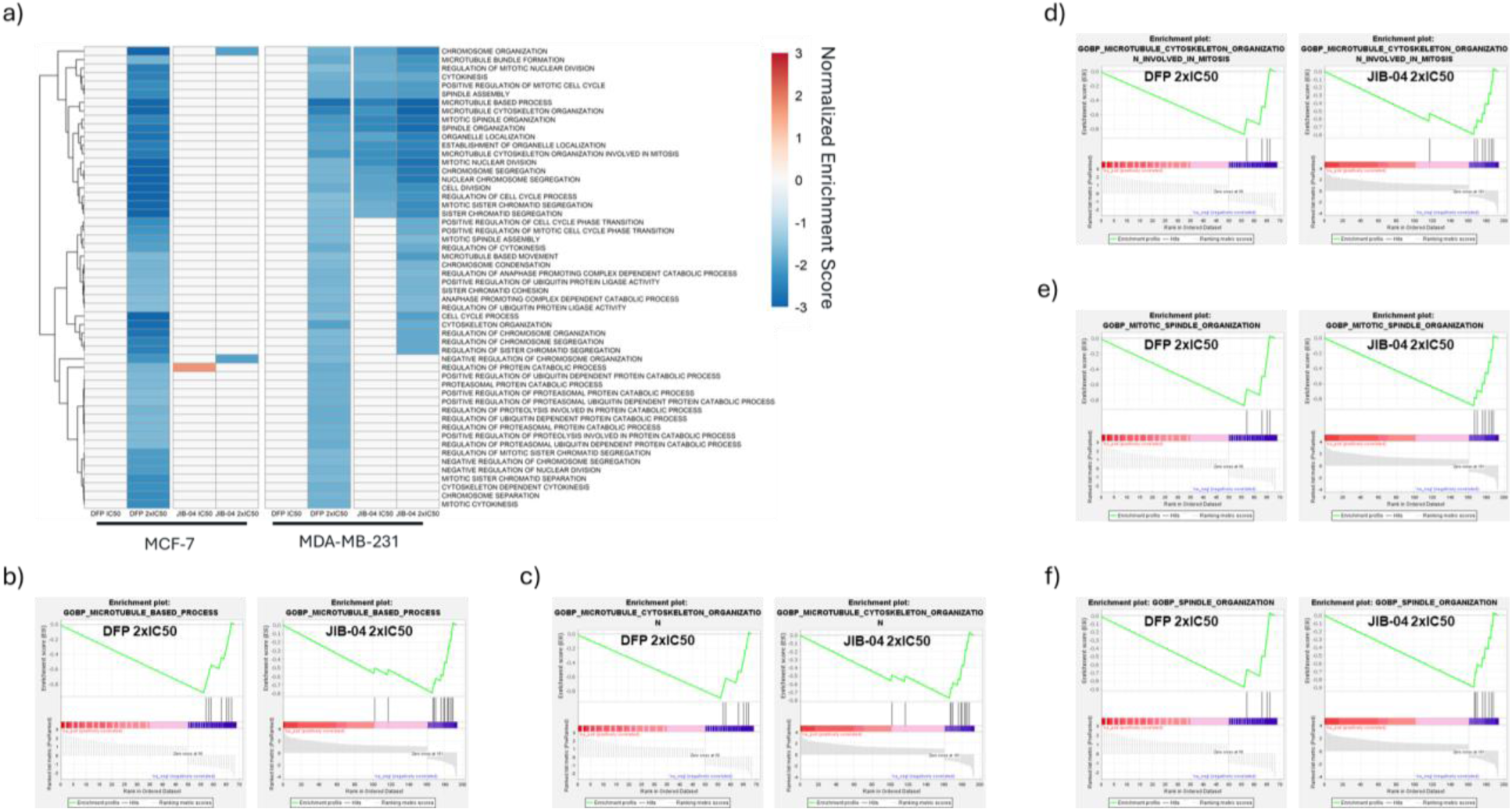
GSEA of GOBP in MCF-7 and MDA-MB-231 cells treated with DFP and JIB-04. a) Heatmap of NES of negatively enriched GOBP gene sets resulting from DFP 2x IC_50_, JIB-04 IC_50_, and JIB-04 2x IC_50_ treatment (p < 0.05, FDR < 0.25). DFP IC_50_ did not significantly enrich GOBP gene sets, DFP 2x IC_50_, JIB-04 IC_50_, and JIB-04 2x IC_50_ negatively enriched gene sets related to cell division, chromosome organization, and microtubule and spindle organization. b) Microtubule-based process enrichment plot for DFP 2x IC_50_ (NES = −2.8) and JIB-04 2x IC_50_ (NES = −3.2) in MDA-MB231 cells. c) Microtubule cytoskeleton organization enrichment plot for DFP 2x IC_50_ (NES = −2.5) and JIB-04 2x IC_50_ (NES = −3.0) in MDA-MB-231 cells. d) Microtubule cytoskeleton organization involved in mitosis enrichment plot for DFP 2x IC_50_ (NES = −2.1) and JIB-04 2x IC_50_ (NES = −2.6) in MDA-MB-231 cells. e) Mitotic spindle organization enrichment plot for DFP 2x IC_50_ (NES = −2.1) and JIB-04 2x IC_50_ (NES = −2.7) in MDA-MB-231 cells. f) Spindle organization enrichment plot for DFP 2x IC_50_ (NES = −2.0) and JIB-04 2x IC_50_ (NES = −2.8) in MDA-MB-231 cells.

More stringent parameters were implemented to identify the most significantly enriched gene sets (p < 0.01, FDR < 0.1). This increase in stringency revealed a total of 5 significantly enriched gene sets in MDA-MB-231 which were compared across both cell lines. Of the gene sets identified, the microtubule-based process gene set, −2.9 (DFP, MCF-7), −2.8 (DFP, MDA-MB- 231) and −3.2 (JIB-04, MDA-MB-231), exhibit the lowest normalized enrichment scores (NES). The other gene sets identified were all subsets of the microtubule-based process: microtubule cytoskeleton organization, −2.9 (DFP, MCF-7), −2.5 (DFP, MDA-MB-231) and −3.0 (JIB-04, MDA-MB-231), microtubule cytoskeleton organization involved in mitosis, −2.6 (DFP, MCF-7), −2.1 (DFP, MDA-MB-231) and −2.6 (JIB-04, MDA-MB-231), mitotic spindle organization, −2.4 (DFP, MCF-7), −2.1 (DFP, MDA-MB-231) and −2.7 (JIB-04, MDA-MB-231), and spindle organization, −2.7 (DFP, MCF-7), −2.0 (DFP, MDA-MB-231) and −2.8 (JIB-04, MDA-MB-231) (Figure 4 and Supplementary Figure S2). Implementing the same stringency for the MCF-7 GOBP data returned approximately 400 gene sets with no similarities in negatively enriched gene sets and minimal overlap in gene sets positively enriched by DFP and JIB-04 at 2x IC_50_ (Supplemental Figure S3)

### KDM inhibition downregulates HIF-1α

Hypoxia is one of the most significantly upregulated pathways by both the DFP and JIB-04. Despite their upregulation of hypoxia, we found that Hypoxia-inducible factor 1α (HIF-1α) was significantly downregulated by DFP at 2x IC_50_ (−1.1, MCF-7, and −1.3, MDA-MB-231) and JIB-04 at IC_50_ (−1.4, MDA-MB-231) and 2x IC_50_ (−2.0, MDA-MB-231). Further probing into HIF-1α direct interactors and downstream targets revealed minimal effects (Figure 5). However, DFP and JIB-04 caused upregulation of EGLN3 (PHD3) in MDA-MB-231 cells and EGLN1 (PHD2) in both cell lines, with DFP eliciting a more profound effect on EGLN1 in MCF-7 cells and JIB-04 more significantly upregulating EGLN3 in MDA-MB-231 cells. The expression levels of traditional targets known to regulate HIF-1α expression in TNBC cells, including the NF-κβ signaling, RAS-RAF-MEK-ERK, PI3K/Akt/mTOR signaling, and JAK-STAT3 pathways, were not affected DFP and JIB-04 (Supplemental Figure S4).^12^

**Figure 5.**
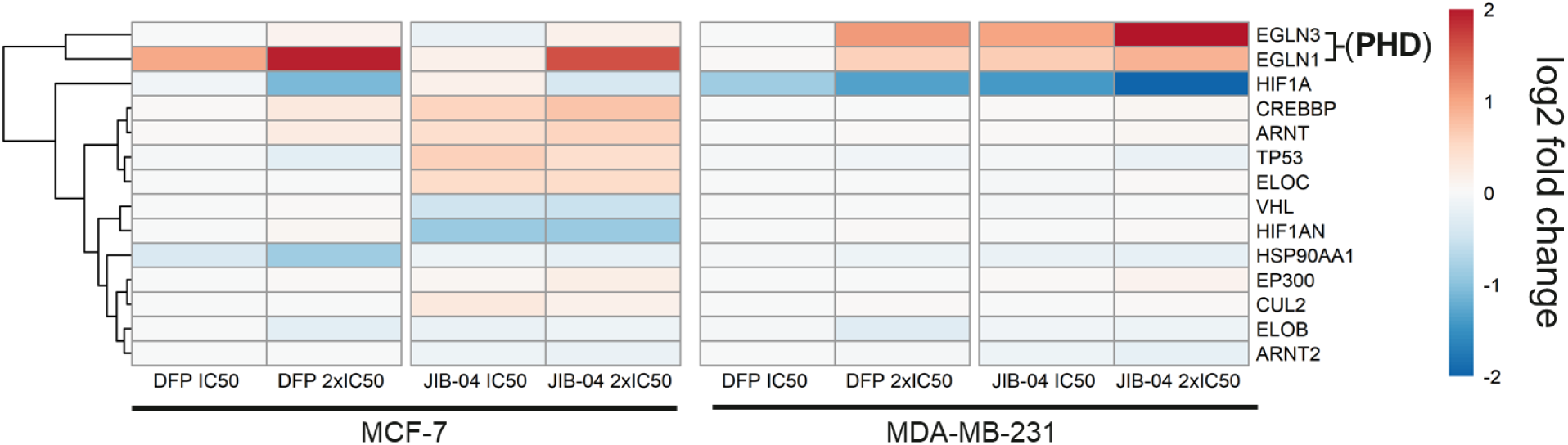
Effects of DFP and JIB-04 on hypoxia inducible factor 1α (HIF-1α) and related genes in MCF-7 and MDA-MB-231 cells. Heatmap of HIF1A direct interactors displaying downregulation of HIF1A by DFP and JIB-04 at 2x IC_50_ in MCF-7 cells and by DFP and JIB-04 at both concentrations in MDA-MB-231. Both DFP and JIB-04 upregulated prolyl hydroxylase domain (PHD) enzymes EGLN3 and EGLN1 in both MCF-7 and MDA-MB-231 cell lines.

### DFP and JIB-04 regulate gene expression in a dose-dependent manner

To identify the potential direct gene targets downstream of DFP and JIB-04 regulation, we screened for differentially regulated genes (DEGs) that had 2 fold-change (FC) and Padj<0.05 upon DFP and JIB-04 treatment as compared with DMSO control samples. In both MCF-7 and MDA-MB-231 cell lines, 2x IC_50_ treatments caused more dramatic gene expression changes than IC_50_ treatments for both DFP and JIB-04 (Figure 6). In response to IC_50_ vs. 2x IC_50_ of DFP treatment, MCF-7 and MDA-MB-231 cells had 270 vs. 1532 and 105 vs. 691 DEGs, respectively, indicating gene expression in response to DFP is dose-dependent. Likewise, JIB-04 treatment of MCF-7 and MDA-MB-231 cells led to more DEGs in 2x IC_50_ than IC_50_ treatments, showing 4644 vs. 5236 and 863 vs. 1425 DEGs in response to IC_50_ vs. 2x IC_50_ treatments, respectively. Consistently, the majority of DEGs elicited in IC_50_ treatments were among the DEGs by 2x IC_50_ treatments. DFP IC_50_ treatments of MCF-7 and MDA-MB-231 cells had 78.8% (213/270) and 80.0% (84/105) of DEGs that were among the DEGs from DFP 2x IC_50_ treatments. Similarly, 80.6% (3744/4644) and 81.8% (706/863) of DEGs from JIB-04 IC_50_ treatments of MCF-7 and MDA-MB-231 cells were also DEGs from JIB-04 2x IC_50_ treatments of the corresponding cells.

**Figure 6.**
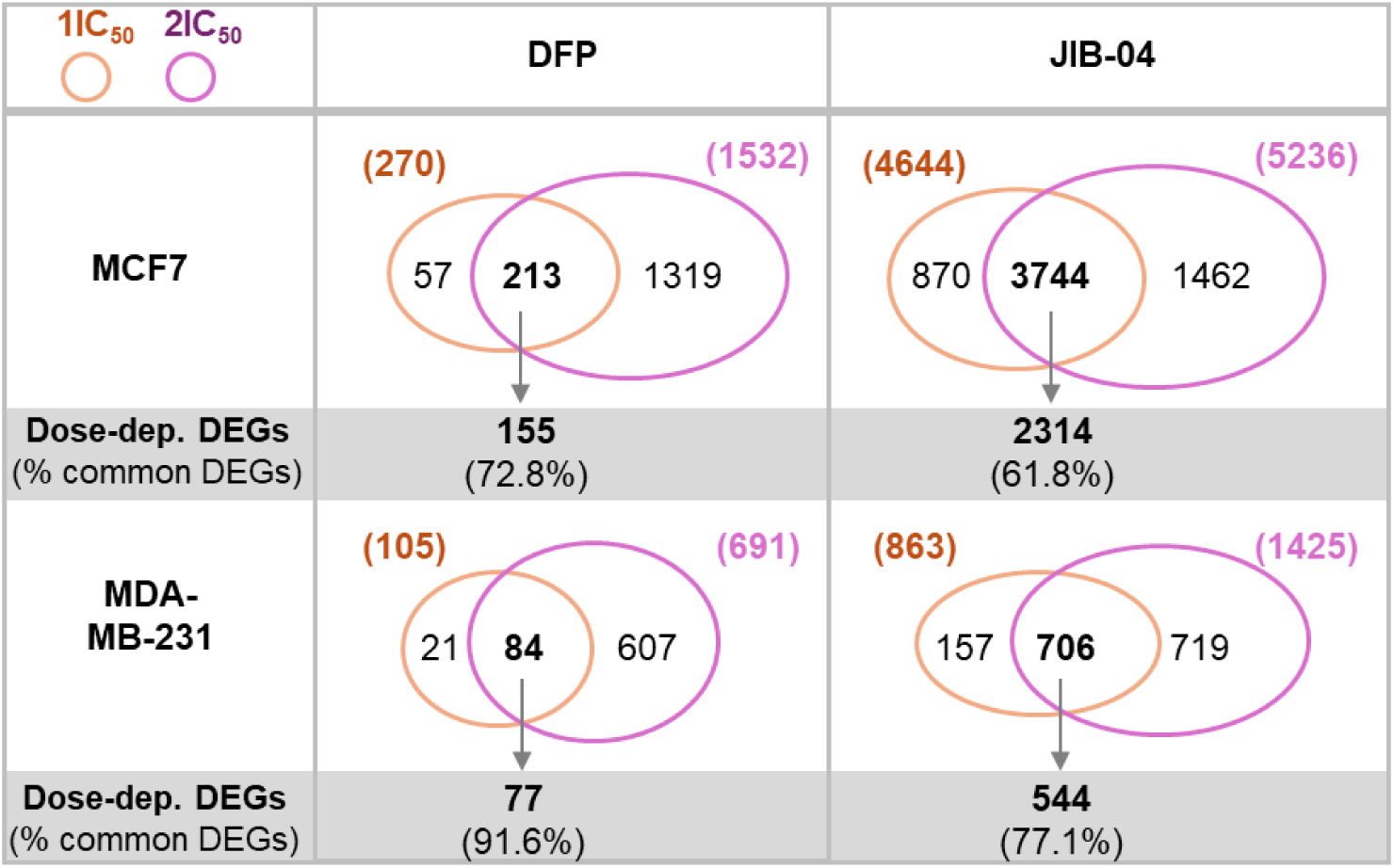
DFP and JIB-04 regulate gene expression in a dose-dependent manner. Venn diagram of DEGs upon drug treatments with expression changes greater than 2-fold (FC>2, Padj<0.05) are shown. Dose-dependent DEGs displayed a higher magnitude of fold-change (FC) in 2x IC_50_ than IC_50_ treatments.

Importantly, the vast majority, ranging from 61.8% to 91.6%, of the common DEGs between IC_50_ and 2x IC_50_ treatments also displayed a higher magnitude of fold-change (FC) in 2x IC_50_ treatments than that from IC_50_ (Figure 6). The dose-dependent responses suggest that these DEGs are direct gene targets regulated by DFP and JIB-04 and that the transcriptional and molecular programs are regulated directly from inhibition on specific KDMs.

### Dose-dependent metastasis DEGs (DDM DEGs) show opposite expression trends in breast cancers as compared to normal breast tissues

To further delineate the effects of DFP caused gene expression changes in relation to breast cancer inhibition, we screened the dose-dependent DEGs for breast cancer metastasis marker genes. Using machine learning approaches, Jung and Yoo predicted 357 breast cancer metastasis marker genes with p<0.01.^13^ Among the dose-dependent DEGs (DD DEGs), 3 (MCF-7/DFP), 6 (MDA-MB-231/DFP), 36 (MCF-7/JIB-04) and 14 (MDA-MB-231/JIB-04) were among the breast cancer metastasis marker genes, which we designated as dose-dependent metastasis DEGs (DDM DEGs).

To further investigate the potential impact of these gene expression changes caused by DFP and JIB-04 treatments, we compared the gene expression of DDM DEGs in cells upon treatments with those of breast cancers from The Cancer Genome Atlas (TCGA database) and the normal breast tissues from the Genotype-Tissue Expression (GTEx) Portal (GTEx database).^14, 15^ Expression levels of each gene of DDM DEGs were examined for 1100 breast cancer samples (TCGA database) and compared with that from 459 normal breast mammary tissues (GTEx database). Many of the DDM DEGs displayed opposite expression changes in breast cancers vs. normal breast tissues to the DEGs elicited in breast cancer cell lines treated with DFP and JIB-04 (Figure 7). For example, RHGB is a downregulated DDM DEG for both DFP and JIB-04 in MCF-7 cells. Its average expression in breast cancers are more than 2-fold higher than in normal breast tissues. DUSP1 is a upregulated DDM DEG for both DFP treatment of MDA-MB-231 cells and JIB-04 treatment of MCF-7 cells, whereas its average expression in breast cancers is more than 2-fold reduced as compared to normal breast tissues (Figure 7). Among the 12 DDM DEGs from JIB-04 treatment of MCF-7 that showed opposite expression changes in TCGA analysis of breast cancers vs. normal breast tissues, 6 were upregulated (SH2D3A, JAG1, DUSP1, HOXA4, JAM2, DENND5A) and 6 were downregulated (RHBG, FAM72D, ATP6AP1L, ZNF233, H2BU1, PLPP6) in MCF-7 cells upon JIB-04 treatment (Figure 7, middle panels). Among the 6 DDM DEGs from JIB-04 treatment of MDA-MB-231 cells that showed opposite expression changes in TCGA analysis of breast cancers vs. normal breast tissues, 5 were upregulated (SLC22A14, ACVR1, FGF18, SVEP1, SERPIND1) and only 1 was downregulated (NAP1L3) in MDA-MB-231 cells upon JIB-04 treatment (Figure 7, bottom panel). These results suggest that DDM DEGs contribute to the reversal of breast cancer cells to normal cells, providing cellular relevance of these drug treatments on the corresponding breast cancers.

**Figure 7.**
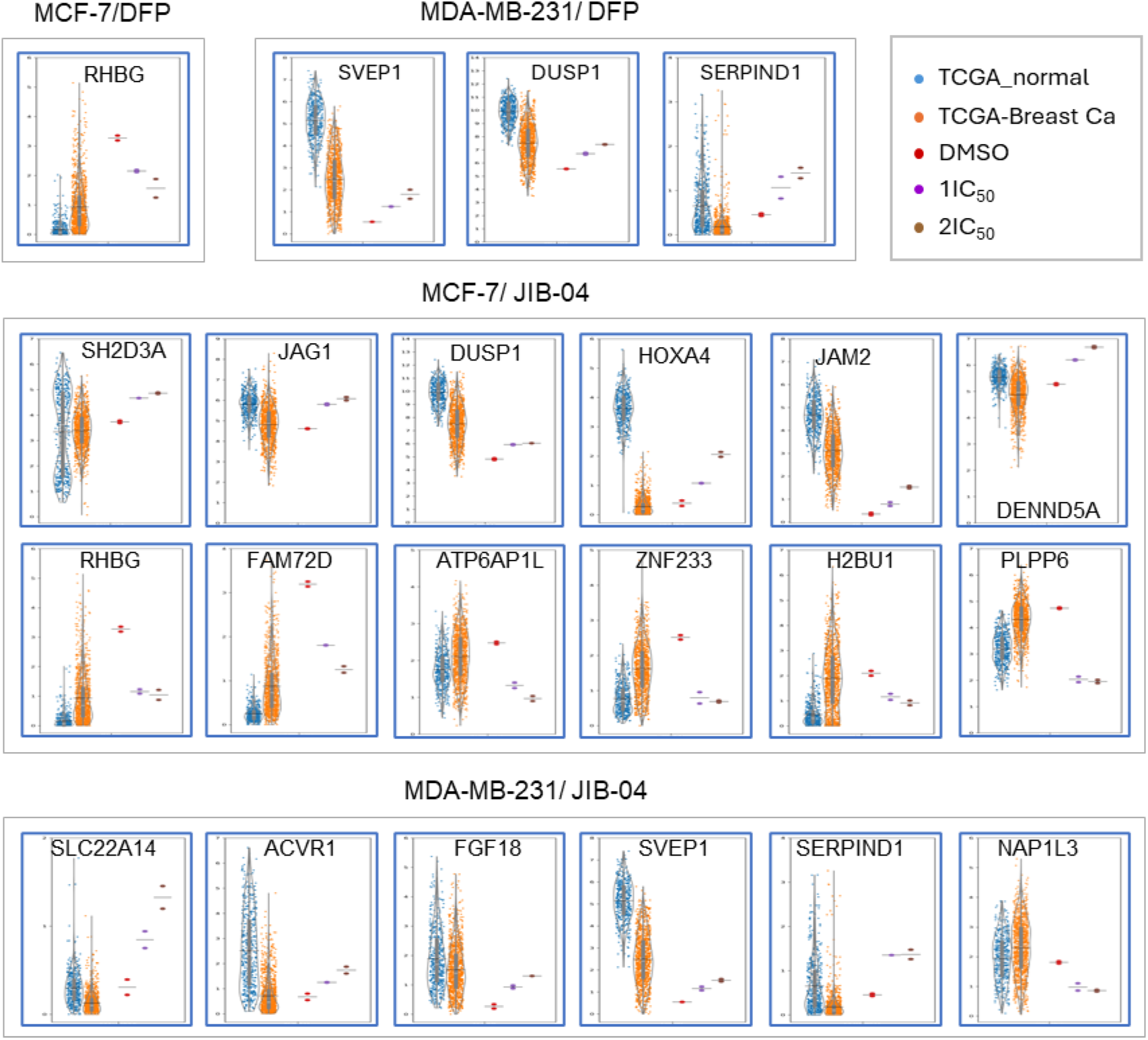
Dose-dependent metastasis DEGs (DDM DEGs) show opposite expression trends in breast cancers as compared to normal breast tissues.

### Western blotting validates RNA seq analysis

These RNA seq observations were validated using Western blotting for the expression levels of key genes whose expressions were significantly perturbed by DFP and JIB-04 in MDA-MB-231 and MCF-7 cell lines. Specifically, we probed the expression of HIF-1α, AURKA, CDKN1A/p21 and CCNB1. Based on the immunoblot results in Figure 8a and Supplemental Figure S5, treatment of MCF-7 cells with graded concentrations of JIB-04 (IC_50_ and 2x IC_50_) and DFP (IC_50_ and 2x IC_50_) for 24 h correspondingly resulted in concentration-dependent downregulatory effects on the expression status of HIF-1α, procaspase 3 and p21. The only exception is that we saw an upregulation of p21 at a higher concentration (4x IC_50_) of DFP under this condition. As for AURKA, EGLN3, and CCNB1, the treatment of MCF-7 cells with either JIB-04 or DFP resulted in a non-significant (*p* > 0.05) upregulation in the levels of the respective proteins. In the case of MDA-MB-231 cells, treatment with either JIB-04 or DFP for 24 h did not induce a significant (*p* > 0.05) effect on HIF-1α levels; however, a slight concentration-dependent decrease in the protein levels was induced by DFP (Figure 8b and Supplemental Figure S6). In addition, while not considered statistically significant at 24 h, treatment of MDA-MB-231 cells with either JIB-04 or DFP induced a concentration-dependent decrease in the levels of EGLN3 and procaspase 3. On the other hand, both JIB-04 and DFP induced a concentration-dependent increase in the expression levels of AURKA and CCNB1 in a non-significant order at 24 h, with the exception of DFP at 4x IC_50_, in which case there was a decrease in the expression levels of both AURKA and CCNB1.

**Figure 8.**
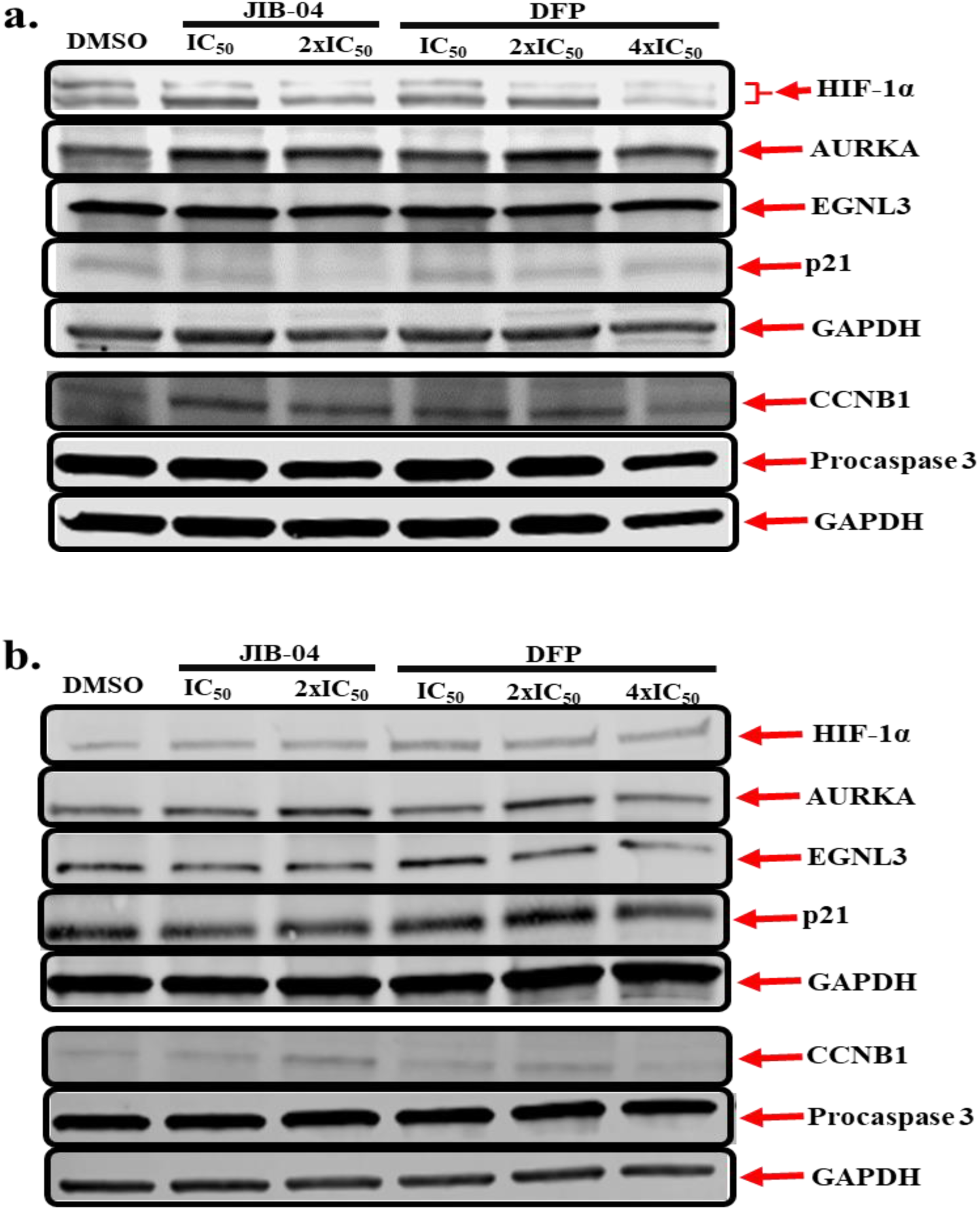
Western blotting analysis of MCF-7 (**a**) and MDA-MB-231 (**b**) cells treated with JIB- 04 and DFP for 24 h reveals the modulatory effects of both compounds on cancer cell survival proteins. Representative full-length gel images are included in the Supplementary Information (Supplemental Figures S7 – S10).

**Figure 9.**
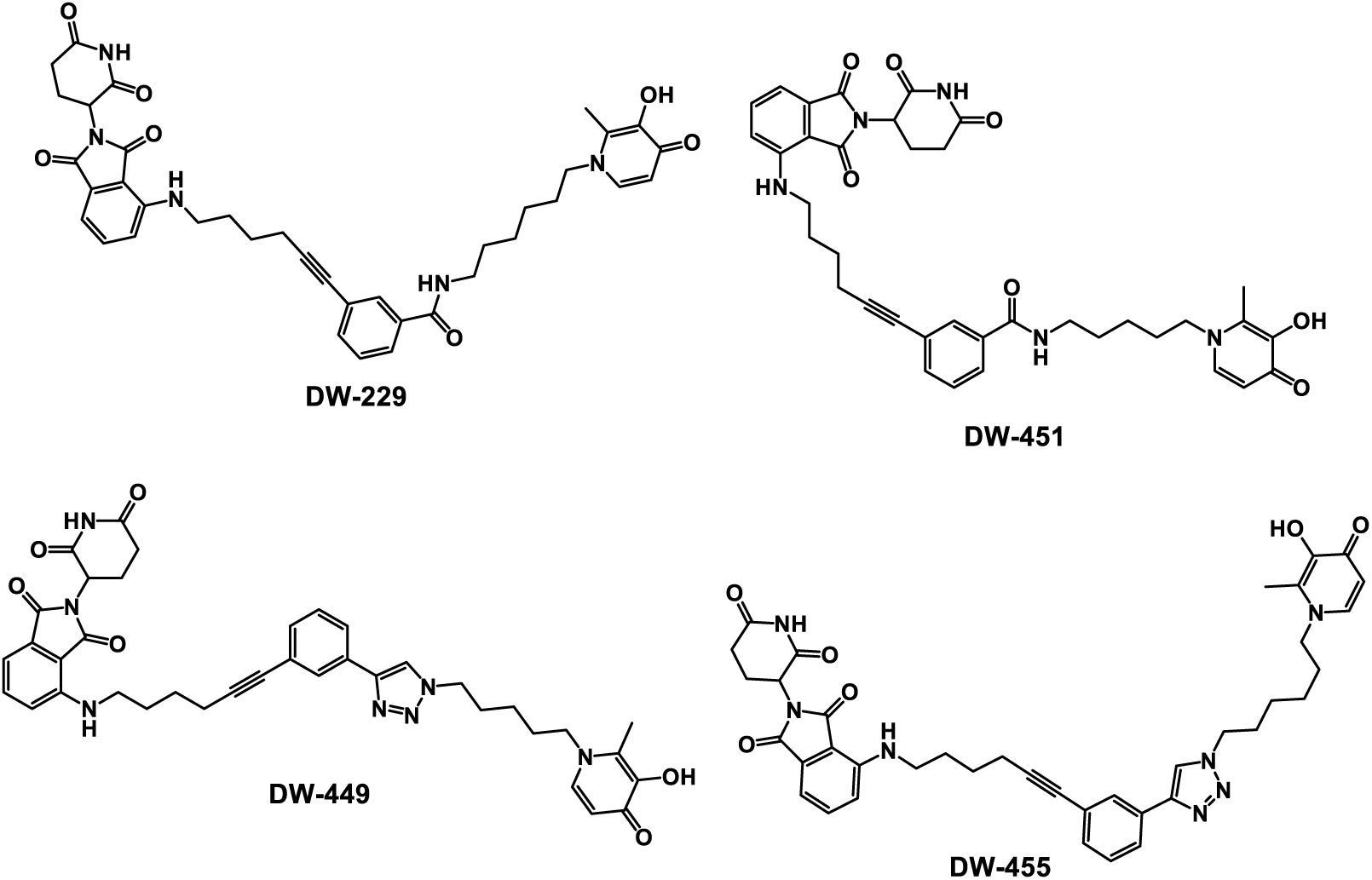
Structures of the designed CRBN E3 ligase DFP-based PROTACs.

### DFP-derived PROTACs elicit enhanced on-target and antiproliferative effects and degrade KDMs

DFP adopts a docked pose at the active sites of several Fe (II)/oxoglutarate dependent KDMs with its N1-methyl group oriented toward the outer rim of the active sites. We have used this information to design DFP-analogs with improved KDM inhibition and antiproliferative activities.^2^ To further confirm our RNA seq and Western blot observations, we used this prior *in silico* result to design two classes of DFP-based PROTACs – amide-linked (**DW-229** and **DW- 451**) and triazole-linked (**DW-449** and **DW-455**) compounds.

We used molecular docking analysis (AutoDock Vina) operating in the flexible mode, as we have described before,^2, 16–18^ to interrogate and verify the interactions of these PROTACs with E3 ligase (PDB:4CI1), and representative Fe (II)/oxoglutarate dependent KDMs KDM6A (PDB:3AVR), KDM5A (PDB: 5CEH), KDM5B (PDB: 6H4Z), KDM3A (PDB: 2Q8C), and KDM2A (PDB: 4QXB). DFP and thalidomide were used as reference compounds with which we compared the docked poses of the DFP-based PROTACS (Figure 10 and Supplemental Figure S11-S14).

**Figure 10.**
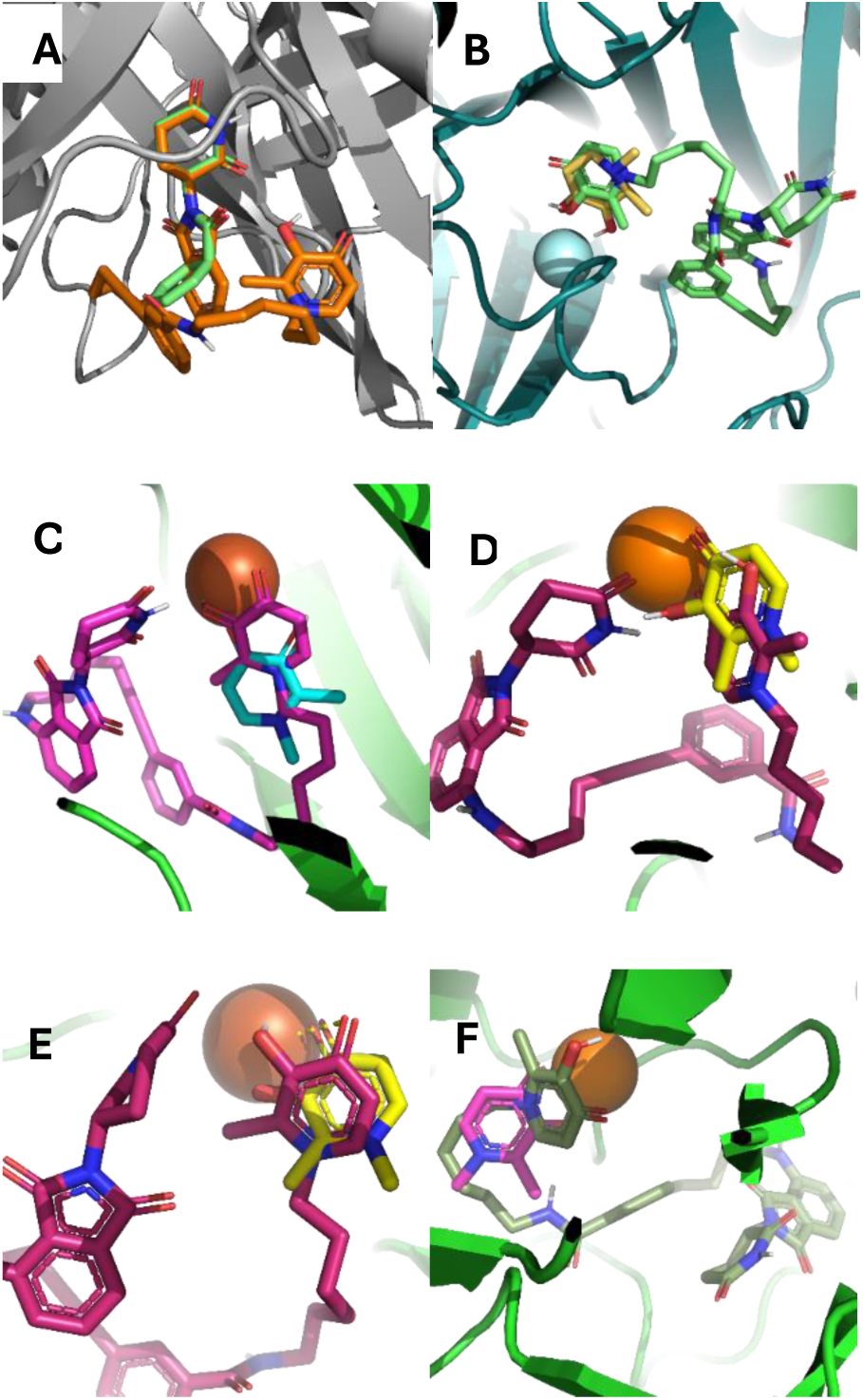
Molecular docking results of compound **DW-229** compared with DFP and thalidomide. (A) Overlay of the E3 Ligase (PDB: 4CI1) dock poses of **DW-229** (orange) and thalidomide (green) shows consistency in the orientation of the thalidomide moiety. (B) At KDM6A (PDB:3AVR) active site, **DW-229** (green) and DFP (yellow) both prefer orientation towards the Fe (II) ion and are at a distance for plausible chelation. (C) At KDM5A (PDB:5CEH) active site, **DW-229** (pink) and DFP (blue) adopt orientations that could enable chelation of the Fe (II) ion. (D) At KDM5B (PDB:6H4Z) active site, **DW-229** (maroon) and DFP (yellow) both bind relatively close to the Fe (II) ion, but with their carbonyl and hydroxyl groups in nearly opposite direction**s**. (E) At KDM3A (PDB:2Q8C) active sites, **DW-229** (maroon) and DFP (yellow) adopt poses with their DFP moiety close to the Fe (II) ion for chelation. (F) At KDM2A (PDB:4QXB) active site, **DW-229** (green) and DFP (pink) adopt different orientations whereby the DFP may not be well positioned for chelation to the Fe (II) ion while the DFP moiety of **DW- 229** binds in close proximity to and engaging in possible cation-π interaction with the Fe (II) ion.

Against E3 ligase, we observed a high degree of conformational similarity between thalidomide and the thalidomide moiety of the PROTACs (Figure 10a). In contrast, DFP and the DFP moiety of the PROTACs adopted varied degrees of docked poses at the active sites of the KDM paralogs investigated. Specifically, DFP and the PROTACS adopted similar docked poses at the active sites of KDM6A and KDM5A (Figure 10b-c). At the active site of KDM5B, the DFP moiety of the PROTACS preferred an orientation in which its carbonyl and hydroxy groups are flipped away from the Fe (II) ion (Figure 10d) while it flipped 180° and at a distance and orientation for plausible chelation with Fe (II) ion at the active sites of KDM3A and KDM2A (Figure 10e-f). In general, the dock scores indicate improvements in binding energies of the PROTACS to the various KDMs relative to DFP, while there is only slight if any change in the dock scores compared with the E3 ligase ligand thalidomide (Supplemental Table S2). These results demonstrate that the DFP and thalidomide moieties of the designed PROTACs can still interact with their target proteins in conformations similar to the free compounds; suggesting that the PROTACs could function as designed.

Encouraged by the foregoing *in silico* results, we synthesize the designed PROTACs **DW- 229, DW-449, DW-451** and **DW-455**. The synthesis of these DFP-based PROTACs is illustrated in Schemes 1a-c. The synthesis of the designed PROTACs **DW-229, DW-449, DW-451** and **DW- 455** was based on the pomalidomide ligand. The E3 ligand with linker was synthesized through a S_N_Ar reaction of 2-(2-dioxo-piperidine-3-yl)-4-fluoro-isoindole-1-dione (**1**) with hex-5-yn-1- amine in DMF and DIPEA, resulting in compound **2** (scheme 1a). The synthesis of the triazole based PROTACs began with azido intermediate compounds **3a-b** built on a previously developed protocol using maltol^2^. The CuSO_4_-mediated cycloaddition reaction of PMB-protected azido maltols with compound **2** furnished the penultimate intermediates **4a-b** in excellent yield. Subsequently, Sonogashira coupling of compounds **4a-b** with compound **2** resulted in intermediate compounds **5a-b.** The next step involved PMB deprotection carried out in 5% TFA in CH_2_Cl_2_ to give the desired triazole-based PROTACs compounds **DW-449** and **DW-451** in moderate yield. The synthesis of amide-based PROTACs began with the reduction of azido maltols **3a-b** under Staudinger conditions to afford the corresponding amines, which subsequently were reacted with 3-iodobenzoic acid in DMF using TBTU to afford the requisite intermediate compounds **6a-b**. Sonogashira reaction of **6a-b** with compound **1** afforded intermediates **7a-b,** which were deprotected with TFA to furnish the amide-based PROTACs compounds **DW-229** and **DW-451.**

**Scheme 1.**
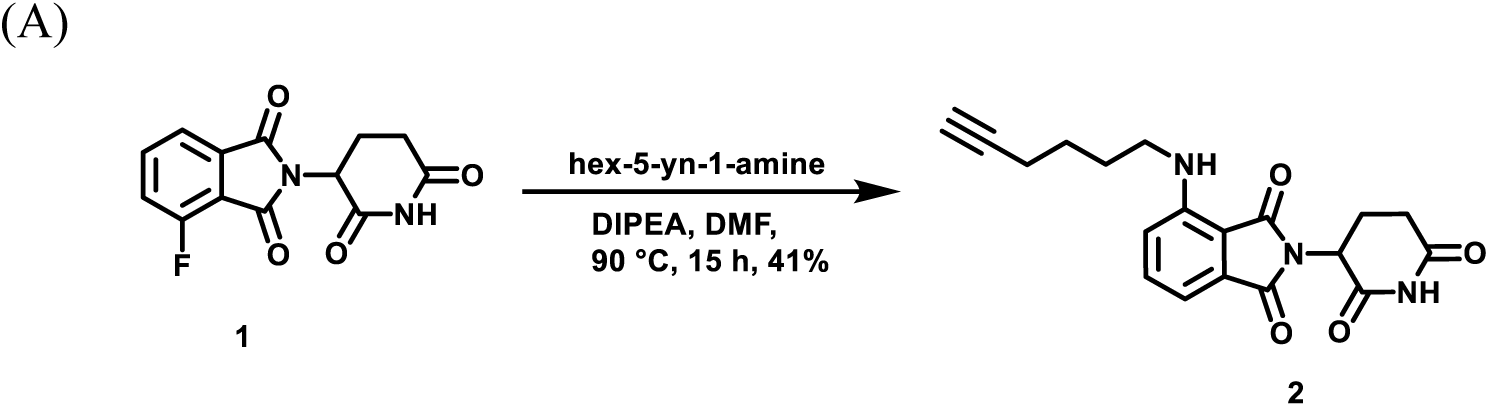
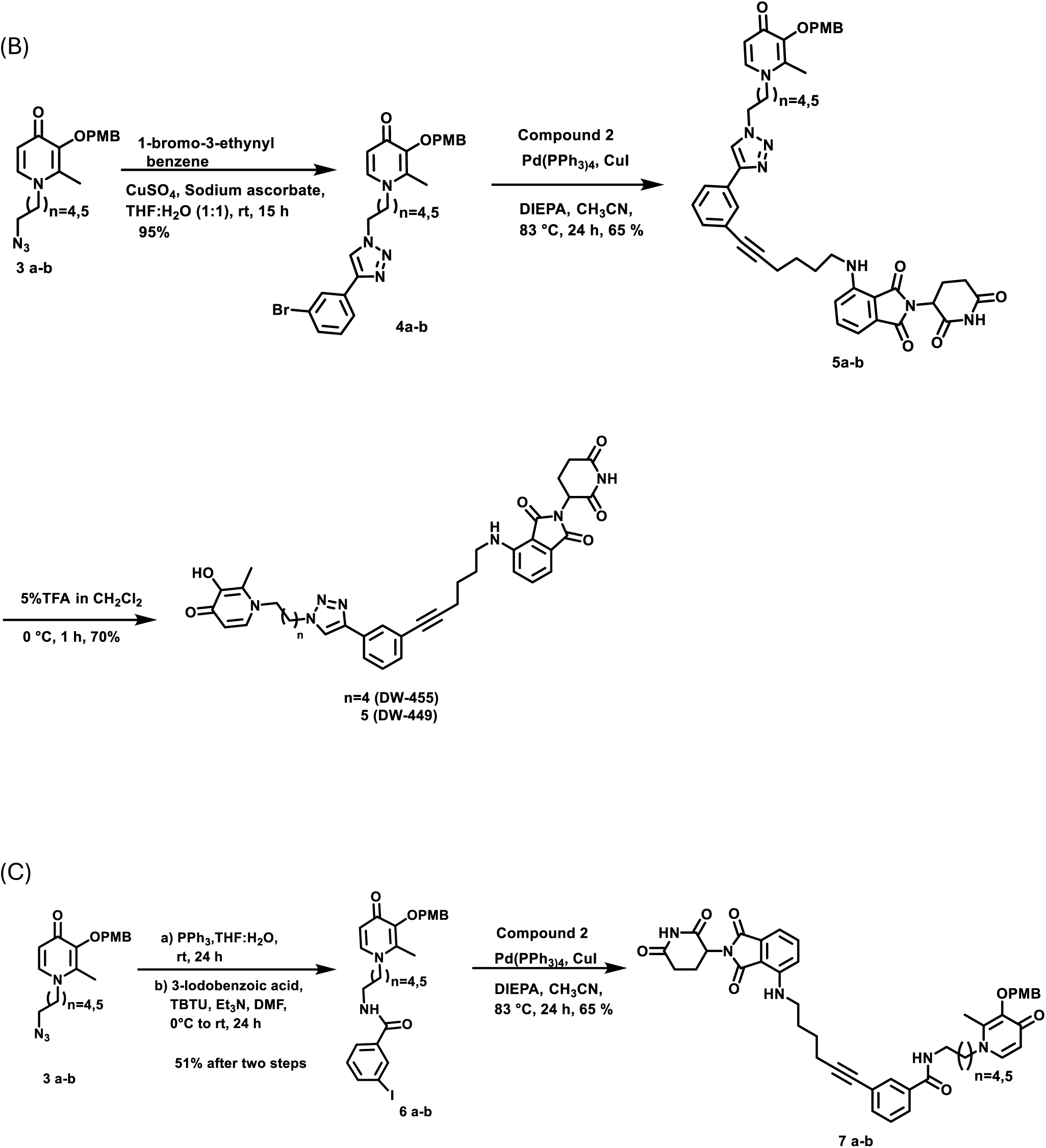
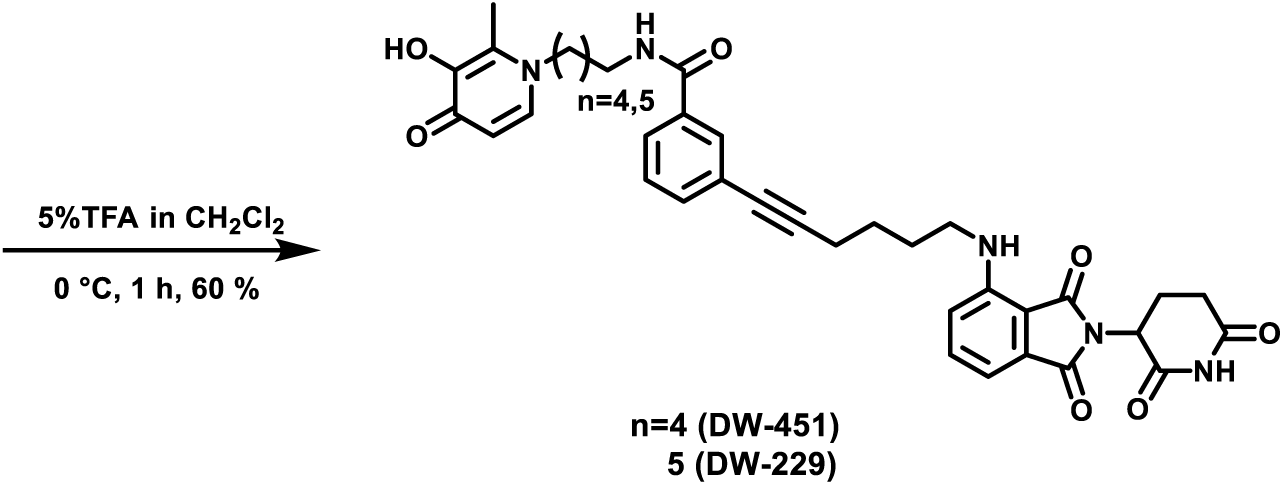
(A) Synthesis of E3 linker; (B) Synthesis of triazole-based series of KDMi PROTACs **DW-449** and **DW-451**, (C) Synthesis of amide-based series of KDMi PROTACs **DW-229** and **DW-451**.

We first evaluated these DFP-based PROTACs in chromatin *in vivo* assay (CiA), a cell-based assay that directly measures the demethylase activities of KDMs within live cells.^19^ We observed that these compounds elicited potent dose-dependent intracellular KDM inhibition activities with **DW-451** showing a somewhat attenuated potency relative to the others. Also, at lower concentrations, **DW-229** transiently induced heterochromatin formation, an effect that is analogous to stimulating KDM activities. However, this effect dissipated at higher concentrations as **DW-229** showed a similar pattern of KDM inhibition as the other PROTACs (Figure 11).

**Figure 11.**
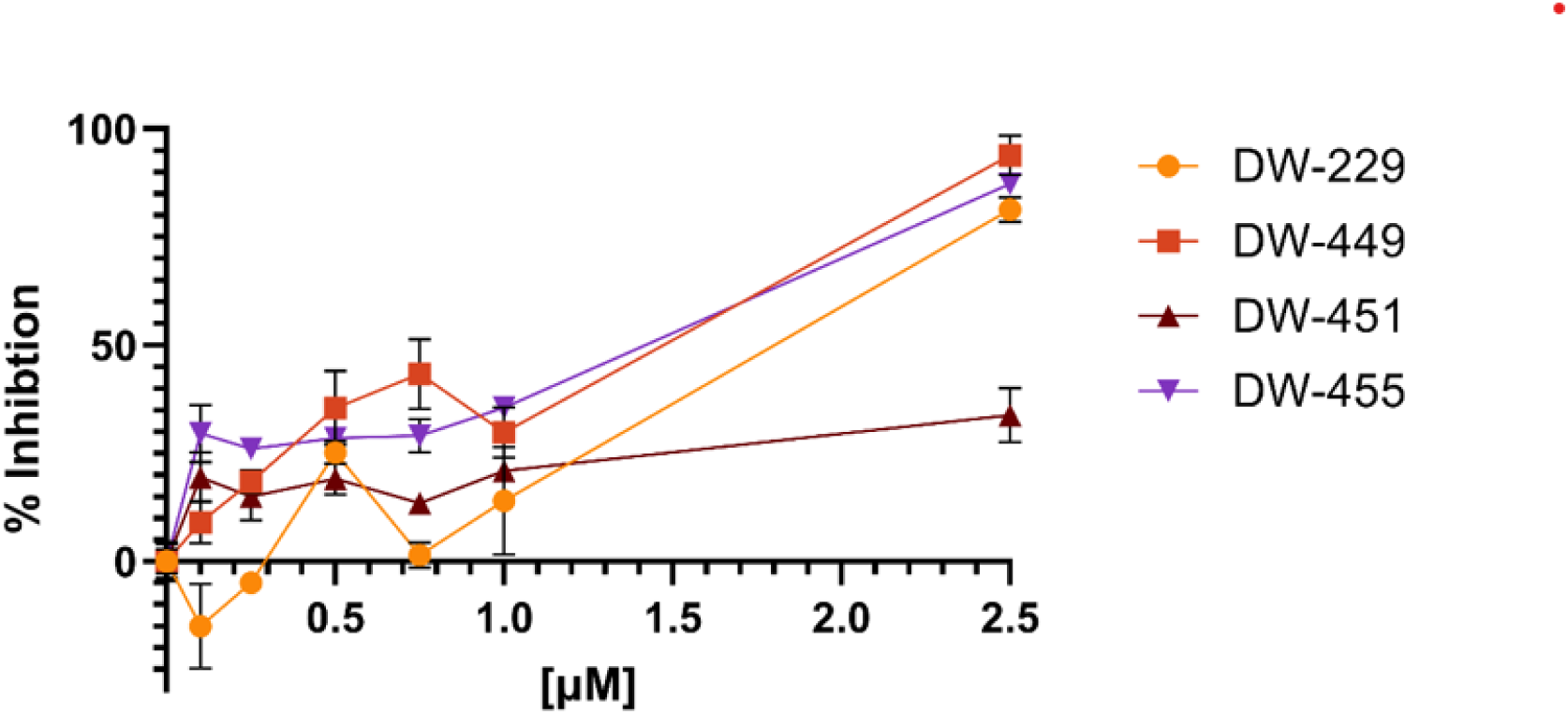
Chromatin *in vivo* assay (CiA) in mES treated with DFP-based PROTACs **DW-229, DW-449, DW-451** and **DW-455** for 48h revealed a dose-dependent inhibition of KDMs. Error bars represent the standard deviation of three biological replicates.

Additionally, these PROTACs inhibit the proliferation of selected cancer cells, with potency enhancement that is >260-fold relative to DFP. More specifically, they elicited significant (*p* < 0.05) cytotoxicity against the tested cancer cell lines – breast (MCF-7, MDA-MB-231, and MDA-MB-453), liver (HepG2 and SK-HEP-1), prostate (DU-145 and LNCaP), and lung (A549) cancer cell lines; and with minimal cytotoxicity against Vero, a normal cell line (Table 1). To probe further into the intracellular mechanism of action of these DFP-based PROTACs we used immunoblotting to determine the effects of representative examples, **DW-229** and **DW-449**, on the expression levels of representative KDM isoforms in MCF-7 and MDA-MB-231 cells. In MCF-7 cells, we observed that **DW-229** significantly (*p* < 0.05) downregulated the nuclear expression levels of KDM isoforms 5B and 6A, while having minimal effects on KDM1A (cytoplasmic and nuclear) and KDM6B (cytoplasmic) (Figure 12a and Supplemental Figures S15-S16). Similarly, **DW-449** also demonstrated a concentration-dependent downregulation of nuclear KDM5B along with little effect on the cytoplasmic expression levels of KDM6B especially at the higher concentrations of treatment in MCF-7 cells (Figure 12b and Supplemental Figures S17-S18).

**Figure 12.**
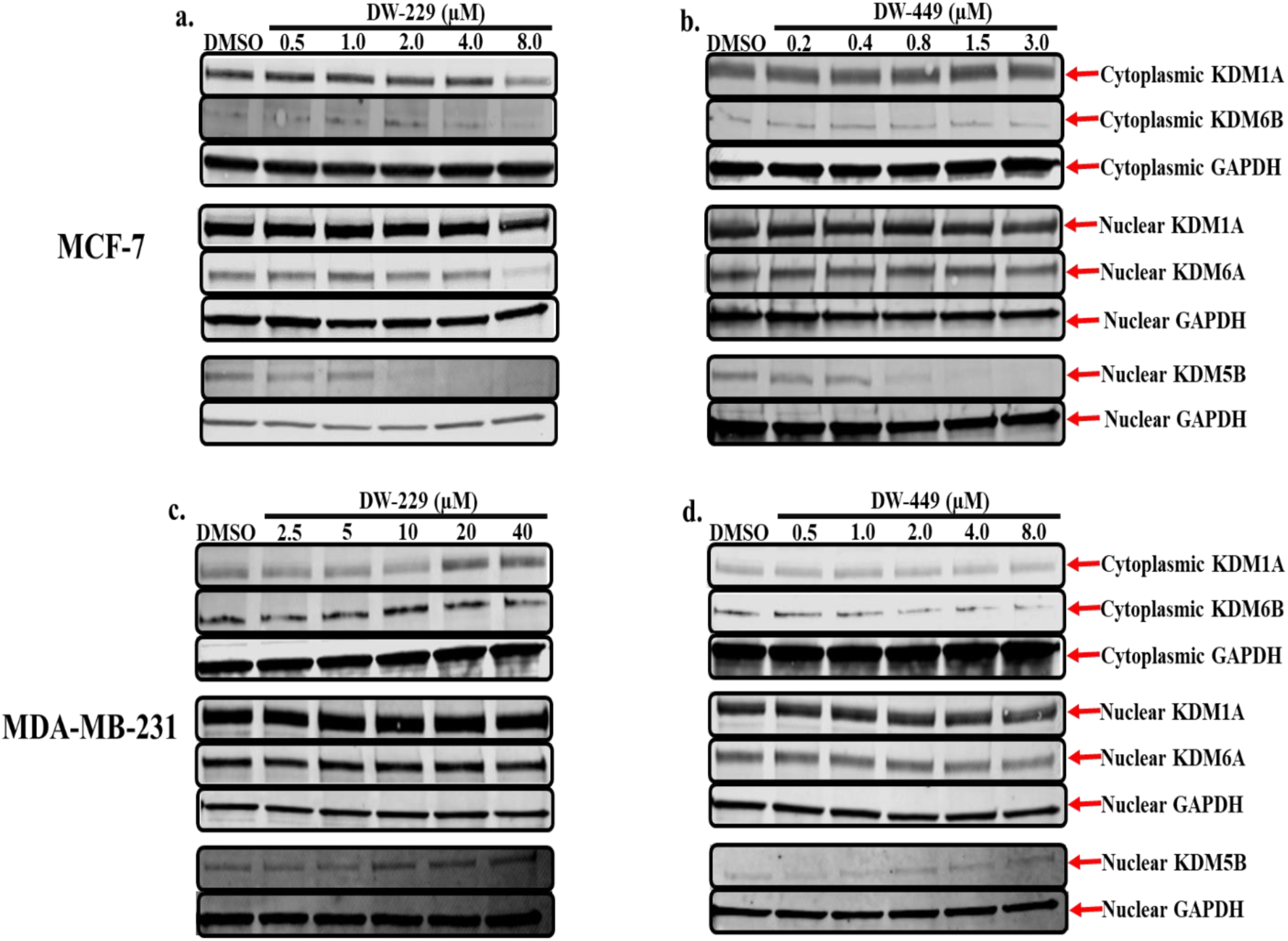
Western blotting analysis of cytoplasmic and nuclear fractions of lysates from MCF-7 (**a**-**b**) and MDA-MB-231 (**c**-**d**) cells treated with graded concentrations of compounds **DW-229** and **DW-449** reveals the modulatory effects of these PROTACs on selected lysine demethylase expression levels. Representative full-length gel images are included in the Supplementary Information (Supplemental Figures S23-S30).

**Table 1.**
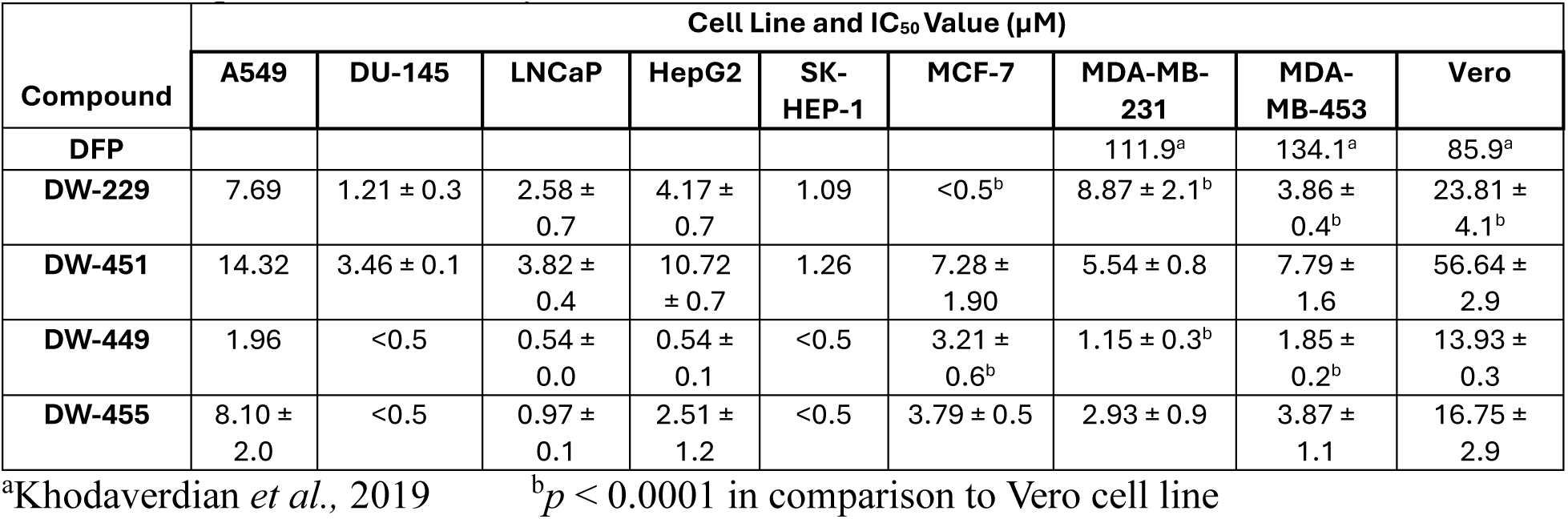
Antiproliferative activity of DFP and DFP-based PROTACs.

| Compound | Cell Line and IC<sub>50</sub> Value (μM) |  |  |  |  |  |  |  |  |
| --- | --- | --- | --- | --- | --- | --- | --- | --- | --- |
|  | A549 | DU-145 | LNCaP | HepG2 | SK-HEP-1 | MCF-7 | MDA-MB-231 | MDA-MB-453 | Vero |
| **DFP** |  |  |  |  |  |  | 111.9<sup>a</sup> | 134.1<sup>a</sup> | 85.9<sup>a</sup> |
| **DW-229** | 7.69 | 1.21 ± 0.3 | 2.58 ± 0.7 | 4.17 ± 0.7 | 1.09 | <0.5<sup>b</sup> | 8.87 ± 2.1<sup>b</sup> | 3.86 ± 0.4<sup>b</sup> | 23.81 ± 4.1<sup>b</sup> |
| **DW-451** | 14.32 | 3.46 ± 0.1 | 3.82 ± 0.4 | 10.72 ± 0.7 | 1.26 | 7.28 ± 1.90 | 5.54 ± 0.8 | 7.79 ± 1.6 | 56.64 ± 2.9 |
| **DW-449** | 1.96 | <0.5 | 0.54 ± 0.0 | 0.54 ± 0.1 | <0.5 | 3.21 ± 0.6<sup>b</sup> | 1.15 ± 0.3<sup>b</sup> | 1.85 ± 0.2<sup>b</sup> | 13.93 ± 0.3 |
| **DW-455** | 8.10 ± 2.0 | <0.5 | 0.97 ± 0.1 | 2.51 ± 1.2 | <0.5 | 3.79 ± 0.5 | 2.93 ± 0.9 | 3.87 ± 1.1 | 16.75 ± 2.9 |

<sup>a</sup>Khodaverdian *et al.*, 2019
<sup>b</sup> $p < 0.0001$ in comparison to Vero cell line

Interestingly, however, the immunoblot results for MDA-MB-231 cells treated with **DW-229** showed a biphasic effect with lower concentrations of **DW-229** being downregulatory, while higher concentrations were upregulatory) on the cytoplasmic expression status of KDM1A with its corresponding nuclear expression levels not affected except at the highest concentration of treatment which is upregulatory. In addition, although not significant, **DW-229** perturbs the cytoplasmic expression status of KDM6B with the effects being biphasic. Going further, **DW- 449** exhibited a similar trend to **DW-229** by non-significantly perturbing the expression of both cytoplasmic KDM6B and nuclear KDM5B (Figure 12c-d and Supplemental Figures S19-S22).

Finally, we used RNAseq to validate the effects of **DW-229**, relative to DMSO and DFP, on the transcriptomic level expression KDMs in MCF-7 cells. We chose MCF-7 for this experiment due to the pronounced and consistent effect elicited by these PROTACs on the expression status of the KDMs investigated in this cell line. We observed that **DW-229**, relative to DFP, downregulated the expression levels of KDM5B (log2 fold change = −1.7), KDM3A (−1.5), and KDM2A (-0.8). Moreover, **DW-229** elicited moderate degradation of KDMs 4A-C, 5C and 6B (Figure 13a-c and Supplemental Table S).

**Figure 13.**
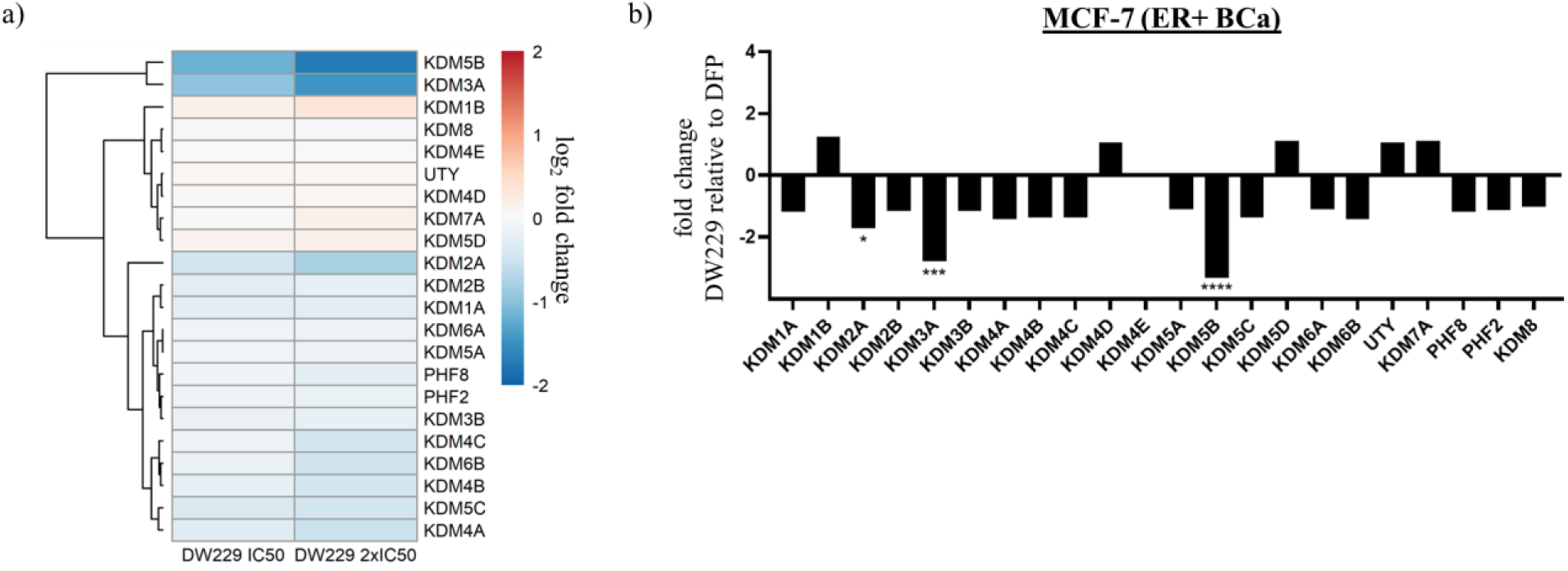
Effects of **DW-229** relative to DFP on KDM expression in MCF-7 cells. a) Log2 fold change heatmap of KDM expression levels in MCF-7 cells treated with **DW-229** relative to DFP. KDM5B and KDM3B were significantly downregulated by **DW-229**. b) Fold change bar graphs of DW229 2x IC50 (0.9µM) relative to DFP 2x IC50 (200µM). * p < 0.05, ** p < 0.01, *** p < 0.001, **** p < 0.0001.

## Discussion

DFP is an FDA approved iron chelator.^20, 21^ We previously reported that DFP attenuates the proliferation of breast cancer cells through pan-selective inhibition of Fe(II)/α-ketoglutarate dependent KDMs, inhibiting six KDM isoforms at low micromolar IC50s, while being considerably less active/inactive against other eleven KDM isoforms.^2^ Here, we further probed the contribution of KDM inhibition to the antiproliferative activities of DFP by first using RNA seq to compare its effects on the transcriptome relative to that of JIB-04, an established KDMi, in breast cancer cells. We initially focused our analysis on the genes adduced to contribute to KDM inhibition activity of JIB-04 in Ewing sarcoma cells.^9^ RNA seq analysis revealed that both JIB- 04 and DFP perturbed the expression pattern of these genes in a similar manner in MCF and MDA-MB-231 cells, with a few exceptions (Figure 2). A more in-depth analysis revealed that DFP and JIB-04 similarly perturbed the expression of genes implicated in hypoxia, cycle progression, microtubule organization and dynamics, and apoptosis, although the effects of JIB- 04 are generally more pronounced (Figures 3-5). Although hypoxia is one of the most significantly upregulated pathways by DFP and JIB-04, we observed that both compounds significantly downregulated HIF-1α in both breast cancer lines.

HIF-1α is a transcription factor that activates several genes important for tumor cell survival, proliferation and invasion. Overexpression of HIF-1α, associated with the aggressiveness of several cancers,^22–26^ has been observed in multiple tumor types including BCa.^27–30^ HIF-1α stability is regulated in response to the intracellular oxygen levels. However, downstream targets, including NF-κβ signaling, RAS-RAF-MEK-ERK, PI3K/Akt/mTOR signaling, and JAK-STAT3 pathways can influence HIF-1α expression independent of intracellular oxygen levels.^31, 32^ Under normoxic conditions, EGLN3 and EGLN1 negatively regulate HIF-1α. Direct interaction of EGLN1 suppressed HIF-1α activity while the hydroxylation of HIF-1α by EGLN3 promotes its proteasomal degradation.^31, 33^ The fact that DFP and JIB-04 have minimal effect on NF-κβ signaling, RAS-RAF-MEK-ERK, PI3K/Akt/mTOR signaling, and JAK-STAT3 pathways but significantly upregulate the expression of EGLN3 and EGLN1 strongly suggests that the KDM inhibition activities of these compounds, due to their attenuation of oxygen consumption by KDMs, mimic normoxic conditions that trigger the degradation of HIF-1α by the oxygen-dependent prolyl hydroxylation activity of the EGLNs. A support for our inference is the observation that the expression levels of EGLN3 and EGLN1 increase under hypoxic conditions in preparation for HIF-1α degradation upon the return of normal oxygen levels.^31^ Another supportive evidence is the observations in the literature demonstrating that the expression of HIF-1α could be modulated through histone demethylation mediated by KDMs.^22, 34, 35^ Specifically, the depletion or inactivation of KDM4A and KDM5B has been linked with the downregulation of HIF-1α and attenuation of tumor aggressiveness.^22, 34^

Genome-wide transcriptome analysis of differential gene expression demonstrates that the majority of DEGs elicited by IC_50_ treatments of DFP and JIB-04 were regulated in a dose-dependent manner (Figure 6), suggesting potent and direct effects in gene expression by KDM inhibition. Furthermore, we uncovered many dose-dependent metastasis DEGs (DDM DEGs) by DFP and JIB-04 show reversal expression trends in breast cancers as compared to normal breast tissues (Figure 7), suggesting potential beneficial effects of these drug treatments to breast cancers.

We subsequently used Western blotting to probe the expression status of protein products of representative genes whose expressions were suggested by RNA seq to be significantly perturbed by DFP and JIB-04 in MDA-MB-231 and MCF-7 cell lines. We focused our analysis on the expressions of HIF-1α, EGNL3, AURKA, CDKN1A/p21 and CCNB1. Western blot data from the treatment of MCF-7 cells with DFP and JIB-04 more closely paralleled the RNA seq data with respect to the expression of HIF-1α and EGNL3, and to some extent p21 at higher dose of DFP. In contrast, the Western blot data revealed that DFP and JIB-04 have no or modest upregulating effects on the expression of AURKA and CCNB1 (Figure 8a). In MDA-MB-231 cells, the RNA seq and Western blotting data for DFP are in agreement as it upregulated the expression EGNL3 at 2x IC_50_ while downregulating that of CCNB1 at 4x IC_50_. JIB-04 has no effect on EGNL3 expression while it stimulated CCNB1 2x IC_50_ (Figure 8b). Both compounds have little or no effect on the expression status of HIF-1α, AURKA and p21 in MDA-MB-231 cells. However, DFP at 4x IC_50_, caused a decrease in the expression levels of both AURKA and CCNB1. For several reasons, including posttranscriptional regulations and the mismatch in the dynamics RNA and protein stability that could be cell-type dependent, it is not always the case that there will be a direct correspondence between RNA seq and Western blot data.

To obtain direct evidence for intracellular engagement with the target KDMs, we used DFP to design two classes of PROTACs – amide-linked (**DW-229** and **DW-451**) and triazole-linked (**DW-449** and **DW-455**) compounds. Analyses of these PROTACs in chromatin in vivo assay (CiA) and cell proliferation assay revealed that their potent KDM inhibition and/or degradation activities translated to a significantly potent antiproliferative effects against the tested cancer cell lines, with potency enhancement ranging from 13-97-fold relative to DFP against the triple-negative breast cancer cell lines. One of the PROTACs, **DW-229**, showed a transient induction of heterochromatin at lower concentrations. This could be due to the initial offsetting gene expression repressive effect of alternative isoforms of target KDMs lacking the demethylase activity, which is eventually overcome by further depletion of the target full-length KDMs at high concentrations of **DW-229**.^36–38^ Subsequent probing of the effects of two representative PROTACs – **DW-229** and **DW-449** – on the expression status of selected KDMs in MCF-7 and MDA-MB-231 cells using Western blotting revealed that they preferentially degraded the Fe(II)/α-ketoglutarate dependent KDMs 5B, 6A and 6B while showing less effect on the FAD-dependent KDM1A. The concentration dependence of the KDM degradation activities of these PROTACs is more consistent in MCF-7 with significant degradation of nuclear KDM5B (Figure 12a-b), which has been reported to play a significant role in breast cancer progression and metastatic behavior.^39^ In MDA-MB-231, **DW-229** degraded KDM6B while it has little effects on KDM5B (Figure 12c). **DW-449** elicited a similar concentration dependent degradation of KDM6B (Figure 12d). In contrast however, **DW-449** only showed evidence of degradation of KDM5B at lower concentrations and restored its levels back to that of no treatment control at higher concentrations. This observation suggests that, against KDM5B in MDA-MB-231 cells, **DW-449** is subject to the hook effect, an attenuation of degradation efficiency commonly seen with high concentrations of PROTACs.^40^

Finally, analysis of RNA seq data from MCF-7 treated with **DW-229** provide additional support for the KDM degradation activities of these PROTACs. **DW-229** non-uniformly downregulated several KDMs in MCF-7 cells, strongly degrading KDMs 2A, 3A and 5B, and moderately degrading KDMs 4A-C, 5C and 6B (Figure 13). Several of these KDMs have been implicated in the survival and aggressiveness of several types of cancer including BCa. Upregulation of KDM2A activity caused silencing of tumor suppressors genes in BCa, promoting tumor growth and stemness.^41–43^ Genetic downregulation and pharmacological inhibition of KDM3A and KDM5B caused significant decrease in the proliferation and invasiveness of BCa.^11, 39, 44–47^ KDM4A−C overexpression has been observed in BCa and the inhibition of the demethylase activities of the KDM4 subfamily has been suggested to be a potential therapeutic strategy for BCa.^48–50^ KDM6B expression is significantly increased in invasive BCa and it is required for TGF-β-induced EMT and BCa invasion via the H3K27me3 marks erasing activity of KDM6B. Knockdown of KDM6B inhibited EMT while overexpression of KDM6B induced the expression of mesenchymal genes and promoted EMT.^51^ In contrast, the roles of KDM5C in BCa etiology are context dependent, with evidence supporting tumor suppressor and oncogenic activities.^52^

In conclusion, we present evidence supporting KDM inhibition as a key mechanism of anticancer activity of DFP. Moreover, we disclosed novel, highly potent DFP-derived PROTACs whose anticancer activities merit further investigation.

## Experimental Section

### Chemicals and reagents

Anhydrous solvents and reagents were purchased from Sigma-Aldrich (St. Louis, MO, USA), Acros, VWR International (Radnor, PA, USA) or Thermo Fisher Scientific (Waltham, MA, USA) and were used without further purification. Analtech silica gel plates (60 F254) were utilized for analytical TLC, and Analtech preparative TLC plates (UV254, 2000 μm) were used for purification. Silica gel (200−400 mesh) was used in column chromatography. TLC plates were visualized using UV light, anisaldehyde, and iodine stains. [HPLC analyses of products were carried out using Phenomenex Luna 91 5 µm C8(2) 100 Å LC column (4.6×250 mm) using Agilent 1260 Infinity II HPLC system. Water (solvent A) and MeCN (solvent B) with 0.1 % TFA were used as the mobile phase at a flow rate of 0.5 mL·min-1 93 with the following gradient: 0–5 min: 5% B, 5–30 min: linear gradient to 100% B, 30–34 min: 100% B, 94 34–35 min: linear gradient to 5% B, 35–36 min: 5 % B, 36–37 min: linear gradient to 100% B, 37–38 min: 95 100% B, 38– 39 min: linear gradient to 5% B. The detection wavelength is at 254 nm and has a flow rate of 0.5 mL/min. Sample concentrations were 250 μM – 1mM, injecting 30 μL. NMR spectra were obtained on a Varian-Gemini 400 MHz and 700MHz magnetic resonance spectrometer. ^1^H-NMR spectra were recorded in parts per million (ppm) relative to the residual peaks of CHCl_3_ (7.24 ppm) in CDCl_3_ or CHD_2_OD (4.78 ppm) in CD_3_OD or DMSO-*d5* (2.49 ppm) in DMSO-*d6*. ^13^C spectra were recorded relative to the solvents peaks with complete hetero-decoupling. MestReNova (version 11.0) was used to process the original NMR “fid” files. High-resolution mass spectra were recorded at Georgia Institute of Technology’s Systems Mass Spectrometry Core facility.

### Synthesis Procedure and Characterization

#### 1) Synthesis of 2-(2,6-dioxopiperidin-3-yl)-4-(hex-5-yn-1-ylamino)isoindoline-1,3-dione (**2**)

A mixture of a 2-(2,6-Dioxopiperidin-3-yl)-4-fluoroisoindoline-1,3-dione 1 (1.55 g, 1 equiv.), dry DMF (10 mL) and DIPEA (1.42 mL, 1.5 equiv.) was added and stirred at room temperature (rt). Subsequently hex-5-yn-1-amine (0.64 mL, 1 equiv.) was added, under positive argon pressure at room temperature. The reaction mixture heated to 90^◦^C for 15 h. The reaction was monitored by TLC and after reaching its completion, the reaction mixture was diluted with water and extracted with EtOAc (DMF was removed carefully by successive washing of brine solution to the organic layer). The organic layer was dried over Na_2_SO_4_, filtered and concentrated under reduced pressure to give a crude product. The crude product was purified by column chromatography, eluting with EtOAc: Hexane (2:8 to 4:6), to give compound **2** as yellow solid. (800 mg, 42 %).

^1^H NMR (400 MHz, CDCl_3_) δ 8.27 (s, 1H), 7.49 (t, 1H), 7.09 (d, *J* = 6.8 Hz, 1H), 6.89 (d, *J* = 8.6 Hz, 1H), 6.25 (t, 1H), 4.90 (m, 1H), 3.31 (t, *J* = 7.1 Hz, 2H), 2.88 (m, *J* = 12.0 Hz, 1H), 2.76 (m, *J* = 12.1 Hz, 2H), 2.25 (m, 2H), 2.13 (m, 1H), 1.98 (S, 1H), 1.80 (m, 2H), 1.65 (m, *J* = 9.8 Hz, 2H).

#### 2. Synthesis of compound **3a-b**

The synthesis of compounds **3a** and **3b** was achieved using a previously published protocol.^2^

#### 3) Synthesis of compound **4a-b**

1-Bromo-3-Ethynyl benzene (1.1 equiv.) and azides **3 a-b** (0.1 g, 1 equiv.) were dissolved in THF (2 mL) and water (2 mL) and stirred at room temperature (rt). Copper sulfate (4 mg, 0.05 equiv.) and sodium ascorbate (5.5 mg, 0.1 equiv.) were added to the reaction mixture, and stirring continued for 15 h. The reaction mixture was diluted with CH_2_Cl_2_ (3X 20 mL) and washed with 1:4 NH_4_OH/saturated NH_4_Cl (3 × 25 mL) and again with saturated NH_4_Cl (25 mL). The organic layer was dried over Na_2_SO_4_ and concentrated under vacuum. The crude product was purified by column chromatography, eluting with CH_2_Cl_2_: MeOH in 10:1 to give **4a-b** as white solids in ∼95 % yield.

**Compound 4a:** ^1^H NMR (400 MHz, CDCl_3_) δ 7.91 (s, 1H), 7.85 (s, 1H), 7.66 (d, J = 9.4 Hz, 1H), 7.34 (d, J = 9.0 Hz, 1H), 7.20 (m, 2H), 7.09 (d, J = 7.4 Hz, 1H), 6.74 (d, 2H), 6.24 (d, 1H), 5.04 (s, 2H), 4.29 (t, J = 6.9 Hz, 2H), 3.68 (s, 3H), 3.60 (t, 2H), 1.95 (s, 3H), 1.84 (m, 2H), 1.53 (m, 2H), 1.22 (m, 2H).

**Compound 4b:** ^1^H NMR (400 MHz, CDCl_3_) δ 7.94 (s, 1H), 7.80 (s, 1H), 7.70 (d, J = 12.2 Hz, 1H), 7.41 (d, 1H), 7.24 (d, 2H), 7.11 (d, J = 7.5 Hz, 1H), 6.77 (d, J = 8.6 Hz, 2H), 6.31 (d, J = 7.5 Hz, 1H), 5.09 (s, 2H), 4.34 (t, 2H), 3.72 (s, 3H), 3.65 (t, 2H), 1.98 (s, 3H), 1.88 (m, 2H), 1.54 (m, 2H), 1.27 (m, 4H).

#### 4) Synthesis of compound **5a-b**

Compound **2** (104 mg, 1 equiv.) and either of the intermediate compounds **4a-b** (100 mg, 1 equiv.), were dissolved in dry acetonitrile (5 mL) under argon. Subsequently, Pd(PPh_3_)_4_ (10 mg 0.05 equiv.) and CuI (3 mg, 0.06 equiv.) were added, followed by Hunig’s base (0.5 mL). The reaction mixture was heated at 75 ^0^C overnight. The reaction mixture was quenched with water (10 mL) and extracted with CH_2_Cl_2_ (3×20 mL) and washed with NH_4_OH/NH_4_Cl 1:1 (10 mL), the two layers separated, and the organic layer was washed sequentially with conc. NH_4_OH/NH_4_Cl 1:1 (2 x 10 mL), brine (30 mL) and dried over Na_2_SO_4,_ and then filtered. The solvent was removed using a rotary evaporator, and the crude material was purified using preparative TLC, eluting with CH_2_Cl_2_: MeOH (10:1), v/v; to afford intermediate product compound **5 a-b** as yellow solid.

**Compound 5a:** ^1^H NMR (400 MHz, CDCl_3_) δ 9.23 – 9.02 (s, 1H), 7.82 – 7.75 (d, *J* = 15.1 Hz, 2H), 7.76 – 7.70 (s, 1H), 7.44 – 7.35 (d, *J* = 7.0 Hz, 1H), 7.32 – 7.27 (d, *J* = 17.5 Hz, 3H), 7.14 –7.07 (d, *J* = 7.5 Hz, 1H), 7.02 – 6.95 (d, *J* = 4.3 Hz, 1H), 6.88 – 6.84 (d, *J* = 8.6 Hz, 1H), 6.82 –6.75 (d, *J* = 6.5 Hz, 2H), 6.38 – 6.31 (d, *J* = 7.5 Hz, 1H), 6.28 – 6.22 (s, 1H), 5.15 – 5.01 (s, 2H), 4.93 – 4.78 (s, 1H), 4.38 – 4.25 (s, 2H), 3.80 – 3.68 (s, 3H), 3.68 – 3.59 (s, 2H), 3.33 – 3.22 (s, 2H), 2.87 – 2.67 (m, 3H), 2.53 – 2.41 (s, 2H), 2.10 – 2.00 (s, 1H), 2.00 – 1.94 (s, 3H), 1.90 – 1.77 (s, 4H), 1.75 – 1.67 (s, 2H), 1.62 – 1.54 (s, 2H), 1.27 – 1.14 (s, 2H).

**Compound 5b:** ^1^H NMR (500 MHz, CDCl_3_) δ 8.75 (s, 1H), 7.83 (s, 1H), 7.74 (s, 2H), 7.44 (t, 1H), 7.34 – 7.27 (m, 4H), 7.11 (d, *J* = 7.5 Hz, 1H), 7.03 (d, *J* = 7.0 Hz, 1H), 6.89 (d, *J* = 8.5 Hz, 1H), 6.80 (d, *J* = 9.5 Hz, 2H), 6.37 (d, *J* = 7.5 Hz, 1H), 6.27 (s, 1H), 5.13 (s, 2H), 4.90 (m,1H), 4.35 (t, 2H), 3.75 (s, 3H), 3.67 (t, 2H), 3.31 (m, 2H), 2.86 – 2.66 (m, 3H), 2.48 (t, *J* = 6.9 Hz, 2H), 2.13 – 2.06 (m, 1H), 2.01 (s, 3H), 1.92 – 1.79 (m, 4H), 1.72 (m, 2H), 1.56 (m, 2H), 1.28 (m, 4H).

#### 5) Synthesis of compound **DW-449** and **DW-455**

The intermediate compounds **5a-b** were separately deprotected by adding 5% TFA in CH_2_Cl_2_ (4 mL) at 0◦C and the reaction was allowed to warm to rt for 1h. The completion of the reaction was indicated by TLC. The reaction mixture was quenched with 10% NaHCO_3_ solution (20 mL) and extracted with CH_2_Cl_2_ (2×20 mL). The organic phases were combined, dried over Na_2_SO_4_, and then filtered. The solvent was removed using a rotary evaporator, and the crude material was purified using preparative TLC [CH_2_Cl_2_: MeOH (10:2), v/v] to afford a compound **DW-449** and **DW-455** as light-yellow solid in ∼70 % yield.

**DW-449:** ^1^H NMR (700 MHz, CDCl_3_) δ 7.82 (s, 2H), 7.43 (s, 1H), 7.32 (s, 2H), 7.13 (s, 1H), 7.02 (s, 1H), 6.88 (s, 1H), 6.78 (S, 1H), 6.31 (d, 1H), 4.92 (m, 1H), 4.36 (bs, 2H), 3.74 (s, 2H), 3.31 (s, 2H), 2.78 (m, 2H), 2.47 (s, 2H), 2.29 (s, 3H), 1.86 – 1.54 (m, 6H), 1.31 (m, 4H). ^13^C NMR (176 MHz, CDCl_3_) δ 171.8, 169.5, 169.3, 168.9, 168.0, 167.7, 167.4, 166.6, 158.8, 158.0, 149.3, 147.0,146.3, 138.3, 136.8, 134.9, 134.5, 132.3, 131.1, 130.7, 129.8, 128.7, 124.9, 124.3, 120.1, 116.6, 113.3,111.3, 109.9,88.7, 80.1, 55.2, 53.5, 50.1, 48.9, 42.0, 34.6, 31.4, 30.6, 30.0, 29.7, 28.3, 26.0, 25.8, 22.7, 19.1, 11.8. HRMS (EI) m/z Calcd. for C_39_H_42_N_7_O_6_ [M+H]^+^: 704.3197, found 704.3195.

**DW-455:** ^1^H NMR (700 MHz, CDCl_3_) δ 7.81 (s, 2H), 7.44 (s, 1H), 7.33 (s, 2H), 7.13 (s, 1H), 7.03 (s, 1H), 6.88 (s, 1H), 6.79 (s, 1H), 6.34 (s, 1H), 6.26 (s, 1H), 4.90 (m, 1H), 4.38 (s, 2H), 3.78 (s, 2H), 3.32 (s, 2H), 2.87 – 2.66 (m, 4H), 2.49 (s, 2H), 2.31 (s, 3H), 2.09 (s, 1H), 1.95 (m, 2H), 1.85 (m, 2H), 1.71 (m, J = 7.4 Hz, 4H), 1.33 (m, 2H). ^13^C NMR (176 MHz, CDCl_3_) δ 171.6, 169.6, 169.4, 168.9, 167.7,158.9, 149.4, 147.2, 146.9, 146.3, 136.8, 136.2, 132.5, 131.3, 130.6, 128.9, 128.8, 128.1, 125.0, 124.4, 120.0, 116.7 114.0, 111.4, 111.3, 110.0, 90.0, 81.0, 55.3, 53.6, 50.0, 49.0, 42.2, 31.5, 30.3, 29.8, 28.4, 25.9, 23.4, 22.7, 19.2, 11. 9. HRMS (EI) m/z Calcd. for C_38_H_40_N_7_O_6_ [M+H]^+^: 690.3040, found:690.3080

#### 6) Synthesis of compound **6a-b**

Either of the azide compounds **3a-b** (200 mg. 1 equiv.) was dissolved in 10 mL of THF: H_2_O (8:2) and cooled to 0 °C. Then triphenylphosphine (176 mg, 1.2 equiv.) was added slowly and reaction mixture was stirred at rt for 24 h. After completion of reaction (monitored by TLC), it was extracted with ethyl acetate (5 x 2 mL). The organic layer was washed with brine, dried over Na_2_SO_4_, filtered and concentrated under reduced pressure to give a crude amine product.

A mixture of the crude amine (0.176 g 1 equiv.) and 3-iodobenzoic acid (122 mg, 1 equiv.) in dry DMF (6 mL) was stirred at 0 °C in an argon atmosphere. Subsequently, TBTU (0.20 g, 1.1 equiv.) and DIPEA (0.27 mL, 3 equiv.) were added, and the reaction was stirred at rt for 15 h. TLC indicated the completion of the reaction. The reaction was quenched with water (50 mL), extracted with CH_2_Cl_2_ (2×30 mL), and the combined organic layer washed with a saturated solution of NaHCO_3_ and brine (30 mL). The organic layer was dried over Na_2_SO_4_, and then filtered. The solvent was removed using a rotary evaporator and the crude material was purified on silica gel column chromatography, eluting with CH_2_Cl_2_: MeOH (12:1), v/v, to afford solid the desired compounds **6 a-b** in ∼51 % yield after two steps.

**Compound 6a**: ^1^H NMR (400 MHz, CDCl_3_) δ 8.15 (bs, 1H), 7.78 (t, *J* = 8.6 Hz, 2H), 7.24 (d, *J* = 8.8 Hz, 2H), 7.19 (d, *J* = 7.3 Hz, 1H), 7.11 (m, 1H), 6.80 (d, *J* = 10.4 Hz, 2H), 6.34 (d, *J* = 7.3 Hz, 1H), 5.07 (s, 2H), 3.76 (s, 3H), 3.71 (t, 2H), 3.40 (m, 2H), 2.04 (s, 3H), 1.66 – 1.57 (m, 4H), 1.31 (m, 2H).

**Compound 6b**: ^1^H NMR (400 MHz, CD_3_OD) δ 8.48 (bs, 1H), 8.14 (m, 3H), 7.77 (d, *J* = 7.6 Hz, 1H), 7.59 (d, *J* = 6.3 Hz, 2H), 7.53 (s, 2H), 7.18 (d, *J* = 12.6 Hz, 2H), 6.78 (d, *J* = 7.6 Hz, 1H), 5.38 (s, 2H), 4.17 (t, 2H), 4.11 (s, 3H), 3.69 (t, 2H), 2.41 (s, 3H), 1.98 – 1.91 (s, 4H), 1.74 – 1.65 (s, 4H).

#### 7) Synthesis of compound **7a-b**

Compound **2** (104 mg, 1 equiv.) and either of the intermediate compounds **6a-b** (100 mg, 1 equiv.) were dissolved in dry acetonitrile (5 mL) under argon. Subsequently, Pd(PPh_3_)_4_ (10 mg 0.05 equiv.) and CuI (3 mg, 0.06 equiv.) were added, followed by Hunig’s base (0.5 mL). The reaction mixture was heated at 75 °C overnight. The reaction mixture was quenched with water (10 mL) and extracted with CH_2_Cl_2_ (3×20 mL) and the combined organic layer washed sequentially with conc. NH_4_OH/NH_4_Cl 1:1 (2 x 10 mL), brine (30 mL) and dried over Na_2_SO_4_, and then filtered. The solvent was removed using a rotary evaporator, and the crude material was purified using preparative TLC, eluting with CH_2_Cl_2_: MeOH (10:1), v/v; to afford intermediate compounds **7 a-b** as yellow solid.

**Compound 7 a:** ^1^H NMR (400 MHz, CDCl_3_) δ 8.98 (s, 1H), 7.82 (s, 1H), 7.71 (d, *J* = 7.9 Hz, 1H), 7.45 (d, *J* = 5.6 Hz, 2H), 7.33 – 7.26 (m, 3H), 7.15 (d, *J* = 7.5 Hz, 1H), 7.03 (d, *J* = 7.0 Hz, 1H), 6.86 (d, *J* = 8.6 Hz, 1H), 6.80 (d, *J* = 8.8 Hz, 2H), 6.35 (d, *J* = 7.5 Hz, 1H), 6.23 (t, 1H), 5.09 (s, 2H), 4.86 (m, 1H), 3.75 (s, 3H), 3.65 (t, 2H), 3.40 – 3.25 (m, 4H), 2.85 – 2.67 (s, 3H), 2.44 (t, 2H), 2.12 – 2.01 (s, 4H), 1.81 (m, 2H), 1.71-1.51 (m, 6H), 1.28 (m, 2H).

**Compound 7 b** 1H NMR (400 MHz, CDCl3) δ 8.50 (s, 1H), 7.75 (s, 1H), 7.67 (d, J = 7.9 Hz, 1H), 7.49 (m, 2H), 7.35 – (d,1H), 7.30(d, 2H), 7.15 (d, J = 7.5 Hz, 1H), 7.07 (d, J = 7.7 Hz, 1H), 6.89 (d, J = 8.7 Hz, 1H), 6.82 (d, J = 8.8 Hz, 2H), 6.41 (d, J = 7.5 Hz, 1H), 6.25 (m, 2H), 5.16 (s, 2H), 4.90 (m, 1H), 3.77 (s, 3H), 3.71 (t, 2H), 3.44 – 3.24 (m, 4H), 2.90 – 2.68 (m, 3H), 2.48 (t, 2H), 2.11 (m, J = 4.4 Hz, 1H), 2.04 (s, 3H), 1.85 (m, 2H), 1.73 (m, 2H), 1.61 – 1.49 (m, 4H), 1.39 – 1.22 (m, 4H).

#### 8) Synthesis of compound **DW-451** and **DW-229**

The intermediate compounds **7a-b** were separately deprotected by adding 5% TFA in CH_2_Cl_2_ (4 mL) at 0◦C and the reaction was allowed to warm to rt for 1h. The completion of the reaction was indicated by TLC. The reaction mixture was quenched with 10% NaHCO_3_ solution (20 mL) and extracted with CH_2_Cl_2_ (2×20 mL). The organic phases were combined, dried over Na_2_SO_4_, and then filtered. The solvent was removed using a rotary evaporator, and the crude material was purified using preparative TLC [CH_2_Cl_2_: MeOH (10:2), v/v] to afford a compound **DW-451** and **DW-229** as light-yellow solid in ∼60 % yield.

**Compound DW-451:** ^1^H NMR (400 MHz, DMSO-*d6*) δ 8.55 (s,1H), 7.83 (s, 2H), 7.64 – 7.35 (m, 5H), 7.13 (s, 1H), 7.01 (s, 1H), 6.62 (s, 1H), 6.09 (s, 1H), 5.07 (m, 1H), 3.91 (s, 2H), 3.37 (s, 2H), 3.24 (s, 2H), 2.87 (s, 1H), 2.56 (s, 2H), 2.27 (s, 3H), 2.03 (s, 2H), 1.78 – 1.50 (m, 6H), 1.38 – 1.20 (s, 2H). ^13^C NMR (176 MHz, CDCl_3_) δ 171.3, 169.6, 169.4, 168.7, 167.7, 167.2, 147.0, 146.4, 136.9, 136.3, 134.7, 134.5, 132.6, 130.5, 129.9, 129.0, 128.7, 128.6, 124.4, 116.7, 114.0, 111.6, 110.1, 90.6, 80.6, 55.3, 53.9, 49.0, 42.3, 39.7, 34.8, 32.0, 31.5, 30.5, 29.8, 29.3, 28.4, 25.9, 23.7, 22.9, 22.8, 19.2, 14.2, 12. HRMS (EI) m/z Calcd. for C_37_H_40_N_5_O_7_ [M+H]^+^: 666.2928, found 666.2994.

**Compound DW-229**: ^1^H NMR (700 MHz, CDCl_3_) δ 7.77 (s, 2H), 7.44 (m, 3H), 7.30 (s, 1H), 7.13 (s, 1H), 7.04 (s, 1H), 6.88 (s, 1H), 6.80 (d, 1H), 6.36 (s, 1H), 6.25 (s, 1H), 4.91 (m, 1H), 3.82 (s, 2H), 3.41 (s, 2H), 3.31 (s, 2H), 2.88 – 2.69 (m, 4H), 2.47 (s, 2H), 2.33 (s, 3H), 2.09 (s, 1H), 1.83 (m, 2H), 1.70 (m, 4H), 1.56 (m, 2H), 1.35 (m, 4H). ^13^C NMR (176 MHz, CDCl_3_) δ 171.6, 169.6, 169.4, 168.9, 167.7, 167.0, 159.9, 146.9, 146.6, 136.9, 136.2, 135.0, 134.3, 132.6, 130.0, 129.0, 128.6, 126.4, 124.3, 116.7, 114.0, 111.5, 110.1, 90.5, 80.6, 56.4, 55.3, 54.0, 51.3, 49.0, 42.2, 39.8, 38.2, 31.5, 30.8, 29.8, 29.5, 28.4, 26.5, 26.0, 22.9, 19.2, 14.2, 11.9. HRMS (EI) m/z Calcd. for C_38_H_42_N_5_O_7_ [M+H]^+^: 680.3084, found 680.3085.

### Cell Culture

MCF-7 cells were cultured in non-phenol red DMEM (Corning, 17-205-CV) supplemented with 10% FBS, 1% Pen/Strep, and 1% L-glutamine, while MDA-MB-231 and Vero cells were cultured in phenol red-containing DMEM (Corning, 10-013-CV) supplemented with 10% FBS, and 1% pen/strep. The other cell lines HepG2, SK-HEP-1, DU-145 were cultured in MEM (Corning) supplemented with 10% FBS, 1% Pen/Strep, and 1% L-glutamine while A549 cells were cultured the same medium as Vero, but without Na pyruvate. In addition, LNCaP and MDA-MB-453 cells were cultured in a complete RPMI-1640 medium. mES CiA cells were cultured in DMEM (Corning) with 15% FBS supplemented with 100 units/mL penicillin/streptomycin, non-essential amino acids (NEAA; Gibco 11140-050), 10 mM HEPES buffer (Corning, 25-060-CI), 55 µM β-mercaptoethanol, leukemia inhibitory factor (LIF), 7.5µg/mL blasticidin (InvivoGen, ant-bl-1), and 1.5µg/mL puromycin (InvivoGen, ant-pr-1).

### RNA seq

MDA-MB-231/MCF-7 cells were seeded in 6-well plates at a density of 1-2 x 10^5^ cells per well in complete media and allowed to grow for 48 h. Each cell line was then treated with the equivalent IC_50_ and 2x IC_50_ concentration of each test agent solution dissolved in 100% DMSO and diluted with the respective media, ensuring that the final DMSO concentration was 1%. After a 24 h incubation period, the cells were trypsinized, followed by centrifugation for 5 min at 500 RCF. The resulting cell pellets were resuspended in cold 1X PBS and centrifuged again for 5 min at 500 RCF, with this washing step repeated twice. Next, using RNeasy Plus Mini Kit and Qiagen QIAcube Connect RNA Purification system, RNA samples were extracted from the cells and their concentrations determined using a DeNovix spectrophotometer. Subsequently, quality control of the RNA samples was carried out using Agilent RNA 6000 Nano Kit which is designed for use with Agilent 2100 Bioanalyzer instrument only. High quality RNA samples having RNA integrity numbers of 7 or above were used for the next step. RNA-seq libraries were prepared based on NEBNext Ultra II Directional RNA Library Prep Kit for Illumina using NEBNext Poly(A) mRNA Magnetic Isolation Module (NEB #E7490). RNA sequencing was conducted on Illumina NovaSeq X Plus PE150 bp reads at the Molecular Evolution Core, Georgia Institute of Technology. The quality of sequencing was measured using Phred quality score (Q score) and it was established that more than 99% of the sequencing reads attained over 99.99% base call accuracy (base call error rate of 1 x 10^-39^). The resulting paired end fastq raw read files were processed using the custom RNAseq Pipeline scripts from the Berkely Gryder Lab on Case Western Reserve University’s (CWRU) high performance computing cluster. Reads were aligned to the GRCh38 reference genome using STAR version 2.5.3a. Transcripts per million (TPM) read counts were measured via RSEM, the base average TPM read counts and normalized log2 fold change values were calculated for each gene using the R package DESeq2. Significant differential expression was defined as |log2 fold change| ≥ 1 and a false discovery rate ≤ 0.25. Rank lists of the differential expressed genes from each sample were submitted to the Molecular Signatures Database (MSigDB) Gene Set Enrichment Analysis (GSEA) tool. Results were exported and visualized using publicly available custom R scripts released by Berkely Gryder from CWRU (https://github.com/GryderArt/VisualizeRNAseq).

### Western Blotting

Cells were seeded into 6-well plates at a density of 1 x 10^6^ cells per well and allowed to incubate for 24 h. Following treatment for 24 h, cells were lysed using radioimmunoprecipitation assay (RIPA) buffer (150 μL) supplemented with phosphatase inhibitor and protease inhibitor on ice for 15 minutes. The lysates were sonicated for 90 s and then centrifuged at 16000 x*g* for 10 min, and the supernatants were collected. Total protein concentration was determined using a BCA protein assay kit. Based on the protein concentration, the lysates were normalized to achieve equal protein concentrations; the denatured lysate was loaded onto TGX MIDI 4–15% gels and electrophoresed at 150 V for 70 min. Subsequently, the gel was transferred onto a Turbo PDVF membrane, blocked with 5% BSA, and incubated overnight at 4 °C with the dsired antibodies. On the following day, the membrane was washed with TBS-T, incubated with secondary antibody, and bands were quantified using ImageJ software.

### Docking Methods

The docking studies were performed using Autodock Vina through PyRx, and using ChemDraw 3D for ligand preparation, Autodock Tools for protein preparation and PyMol for visualization^53–55^. The general procedures for preparation of crystal structure and molecular docking were done following the procedure that was previously published^56^. The one deviation from the procedure was all docking was done with an exhaustiveness of 32.

### Chromatin *in vivo* Assay (CiA)

For all experiments, mouse embryonic stem cells (mES) containing the CiA components were generated as previously described.^8, 19, 57^ Briefly, cells containing a replacement of a single Oct4 allele with a green fluorescent protein (GFP) gene were infected with lentivirus to stably integrate plasmids N118 (LV EF-1α-Gal-FKBPx1-HA-PGK-Blast) and N163 (nLV EF-1α-HP1α (CS)-Frbx2(Frb+FrbWobb)-V5-PGK-Puro). mES cells where then were seeded into gelatin coated 96 well plates at 10k cells / well. 24 h after seeding, the media was changed and replaced with media only (positive control), media with 6nM rapamycin (negative control), or media with 6nM rapamycin and a varying concentration of a compound of interest (0.1-10 µM). Each condition was done in three biological triplicates. Rapamycin was obtained from LC Laboratories (Woburn, Massachusetts). After 48 h of exposure, cells were washed with PBS, collected using 0.25% trypsin-EDTA, quenched with growth media, and transferred to a non-tissue culture treated U-bottom 96-well plate before being analyzed with an Attune NxT Acoustic Focusing Flow Cytometer with an autosampler (ThermoFisher). GFP signal was measured with a 488nM laser with a 530/30 filter; autofluorescence was also measured with a 637nM laser with a 670/14 filter. Collected mES cells were gated to include only live, single cell populations displaying no autofluorescence. A bifurcating gate was applied to generate a GFP positive (%GFP+) and GFP negative population for each sample, represented as a percentage of cells. Mean %GFP+ were calculated for the negative and positive control populations. Then, a percent inhibition value was calculated for each sample well and the negative control wells; % inhibition is defined as 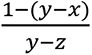 where x = the %GFP+ of the sample, y = the mean %GFP+ of the positive control wells, and z = the mean %GFP+ of the negative control wells. CiA data was recorded at the University of North Carolina at Chapel Hill’s Flow Cytometry Core Facility.

### MTT Cell Viability Assay

Using standard protocol, with minor modifications,^58^ cells, at a density of 4.5 × 10^3^/well and a volume of 100 μL, were seeded in 96-well transparent tissue culture plates and allowed to adhere for 24 h. Thereafter, the culture medium was aspirated, and the cells were treated for 72 h with 100 μL of varying concentrations (0.5 – 100 μM) of the compounds, dissolved in dimethyl sulfoxide (DMSO) but diluted with the respective culture media to ensure a final maximum DMSO concentration of 1%. Subsequently, 10 μL of MTT reagent (5 mg/mL) was added to the culture medium and the plates were incubated for 3 h, after which the MTT reagent/medium was carefully aspirated. Finally, the formed formazan crystals were dissolved in 100% DMSO (100 μL/well) and absorbance values were determined at 570 nm using multimode plate reader (Tecan Infinite M200 Pro, Männedorf, Switzerland), and the percent cell viability was computed with respect to untreated controls.

### Declaration

The authors declare no competing financial interest or personal relationships that could be perceived as influencing this study.

### Author Contributions

A.J., J.O.O. and D.W. contributed equally to the manuscript. Experimental Design: A.J., J.O.O., D.W., B.W., R.K., T.J.N., N.A.H., Y.F. and A.K.O. Data Generation: A.J., J.O.O., D.W., A.B., B.W., R.K., R.Y., T.J.N., B.J.C., and J.M. Data Analysis: A.J., J.O.O., D.W., A.B., B.W., R.K. and T.J.N. Supervision: N.A.H., Y.F. and A.K.O. Manuscript Writing: A.J., J.O.O., D.W., B.W., T.J.N., N.A.H., Y.F. and A.K.O. Manuscript Review: All. Funding: N.A.H., Y.F. and A.K.O.

## Supporting information

DFP_RNA Seq_Suppl Info

## Acknowledgements

This project was financially supported by NIH grants R01CA252720 and T32CA244125 (UNC- Integrated Translational Oncology Program to UNC/TJN). The UNC Flow Cytometry Core Facility (RRID:SCR_019170) is supported in part by a Cancer Center Core Support Grant (P30 CA016086) to the UNC Lineberger Comprehensive Cancer Center.

**Notes:** The authors declare no competing financial interest.

## Graphical Abstract

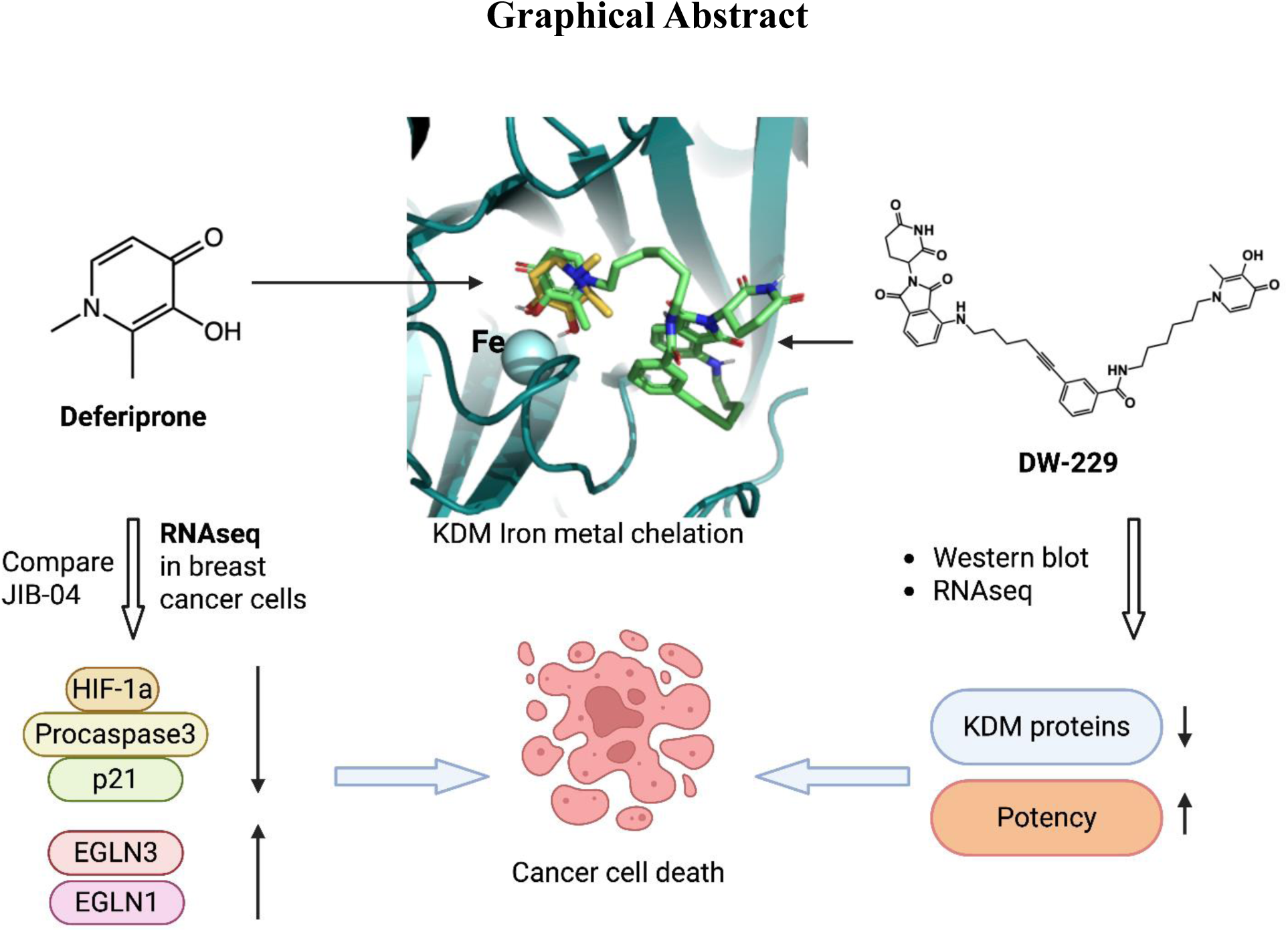

## Notes

### Competing Interest Statement

The authors have declared no competing interest.

