## Supplementary material for "Transcriptomic Signature and PROTAC Strategy Revealed Histone Lysine Demethylase as a Target of Anticancer Activity of Deferiprone": DFP_RNA Seq_Suppl Info

[1]School of Chemistry and Biochemistry, Georgia Institute of Technology, Atlanta, GA 30332-0400, USA; [2]School of Biological Sciences, Georgia Institute of Technology, Atlanta, GA, 30332-0400, USA; [3]The University of North Carolina Eshelman School of Pharmacy, Chapel Hill, NC, 27599, USA; [4]Parker H. Petit Institute for Bioengineering and Bioscience, Georgia Institute of Technology, Atlanta, GA 30332-0400, USA.

[¶]These authors contributed equally to the manuscript.

**Keywords:** Histone lysine demethylase; Deferiprone; RNA seq; PROTAC.

#### Supporting information

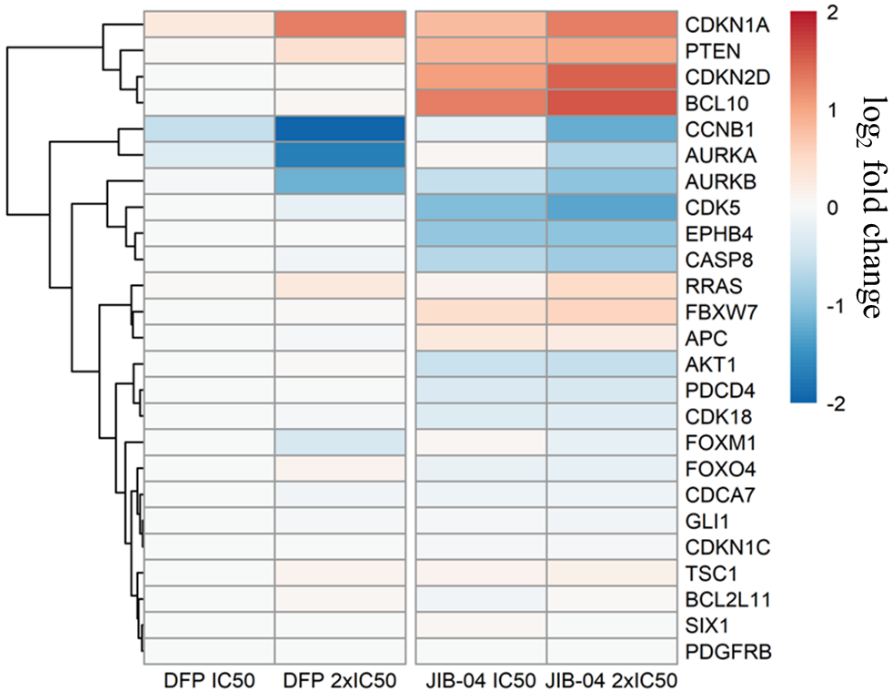


**Figure S1.** Effects of DFP and JIB-04 on genes involved in KDM inhibition in MCF-7 cells. Log2 fold change heatmap of genes significant to JIB-04 inhibition in Ewing Sarcoma displaying upregulation of CDKN1A (p21) by DFP 2x $IC_{50}$ (log2 fold change = +1.3) and JIB-04 2x $IC_{50}$ (log2 fold change = +1.3), downregulation of CCNB1 (DFP 2x $IC_{50}$ log2 fold change = -2.3, JIB-04 2x $IC_{50}$ log2 fold change = -1.2), downregulation of AURKA (DFP 2x $IC_{50}$ log2 fold change = -1.7, JIB-04 1μM log2 fold change = -0.7), and downregulation of AURKB (DFP 2x $IC_{50}$ log2 fold change = -1.2, JIB-04 2x $IC_{50}$ log2 fold change = -1.0). Additionally, JIB-04 2x $IC_{50}$ upregulated PTEN, CDKN2D, and BCL10 (log2 fold change = +1.0, +1.5, +1.6, respectively) and downregulated CDK5 and EPHB4 (log2 fold change = -1.3, -1.0, respectively).

**Table S1.** Effects of DFP and JIB-04 on genes involved in JIB-04 KDM inhibition. The significance of log2 fold change DFP 2x $IC_{50}$ and JIB-04 2x $IC_{50}$ treatment p-values of DFP 2x $IC_{50}$ and JIB-04 2x $IC_{50}$ treatment relative to DMSO in MDA-MB-231 and MCF-7 cells. Gene names arranged in descending order of expression levels resulting from JIB-04 treatment to MDA-MB-231 cells. ns = not significant.

| | **MDA-MB-231 (TNBC)** | | **MCF-7 (ERα+ BCa)** | |
|---|---|---|---|---|
| GeneID | **DFP 2x $IC_{50}$** | **JIB-04 2x $IC_{50}$** | **DFP 2x $IC_{50}$** | **JIB-04 2x $IC_{50}$** |
| CDKN1A | 6.3e-63 | 4.0e-88 | 1.7e-23 | 2.3e-26 |
| CCNB1 | 2.8e-28 | 3.1e-37 | 8.4e-19 | 9.0e-09 |
| AURKA | 2.0e-19 | 2.3e-26 | 5.2e-14 | 6.9e-05 |
| BCL10 | ns | 9.0e-03 | ns | 1.2e-05 |
| CDKN2D | ns | 3.3e-02 | ns | 1.8e-03 |
| PDCD4 | ns | ns | ns | ns |
| FBXW7 | ns | ns | ns | ns |
| AKT1 | ns | ns | ns | 7.1e-05 |
| TSC1 | ns | ns | ns | ns |
| BCL2L11 | ns | ns | ns | ns |
| CDCA7 | 2.5e-02 | 2.5e-03 | ns | ns |
| AURKB | 4.3e-02 | 5.4e-03 | 7.6e-05 | 7.7e-04 |
| SIX1 | ns | ns | ns | ns |
| RRAS | ns | ns | ns | 4.7e-02 |
| CASP8 | ns | ns | ns | 3.0e-02 |
| PDGFRB | ns | ns | ns | ns |
| GLI1 | ns | ns | ns | ns |
| FOXO4 | ns | ns | ns | ns |
| CDK18 | ns | ns | ns | ns |
| CDKN1C | ns | ns | ns | ns |
| APC | ns | ns | ns | ns |
| CDK5 | ns | ns | ns | 2.3e-04 |
| EPHB4 | ns | ns | ns | 3.22e-05 |
| PTEN | ns | ns | 4.0e-02 | 6.0e-04 |
| FOXM1 | ns | ns | 3.6e-02 | ns |

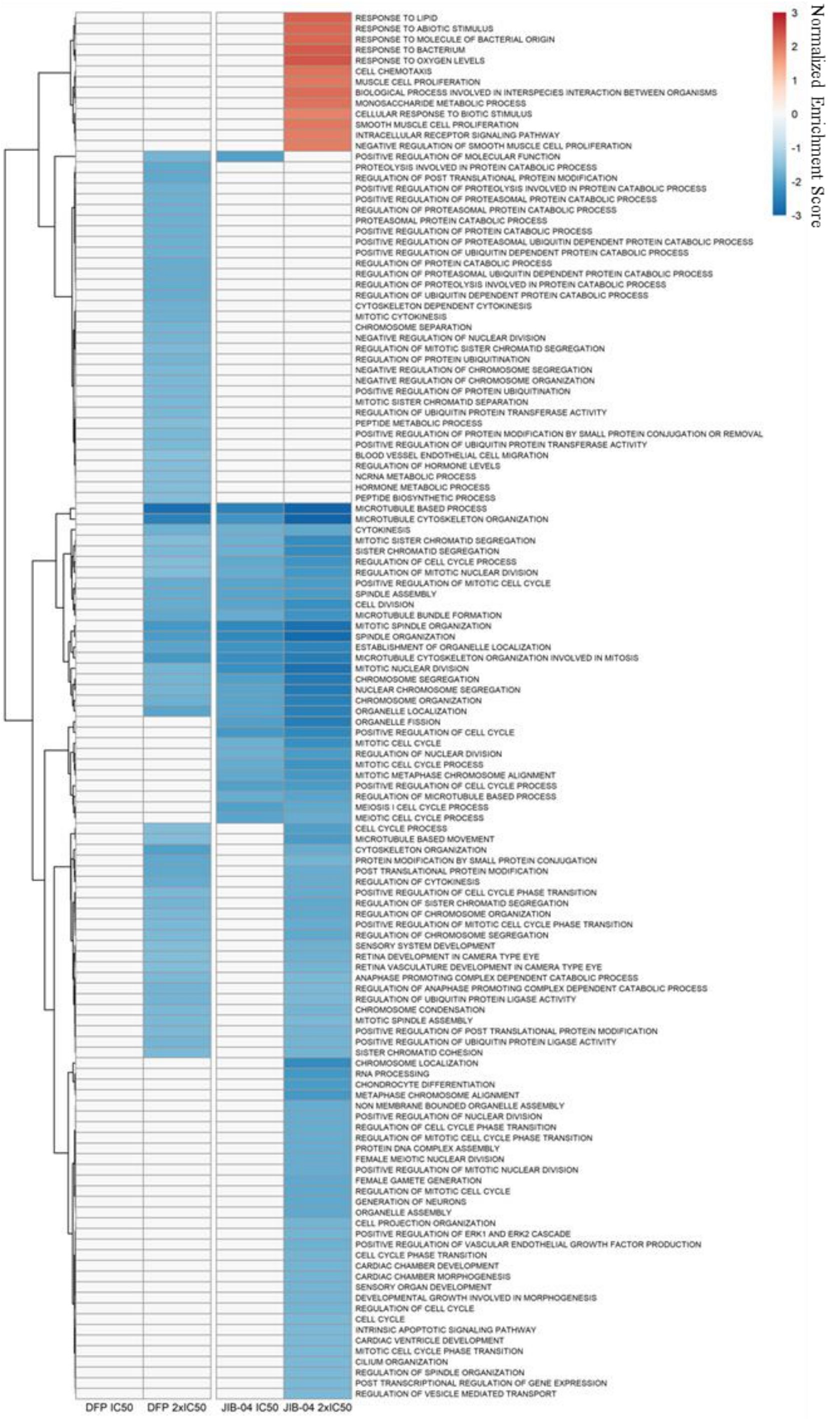


**Figure S2.** Full Gene Ontology Biological Processes (GOBP) Gene Set Enrichment Analysis (GSEA) heatmap of DFP and JIB-04 treatment of MDA-MB-231 cells. Significant gene set enrichment was defined by $p < 0.05$ and false discovery rate (FDR) < 0.25. DFP $IC_{50}$ did not significantly enrich GOBP gene sets, DFP 2x $IC_{50}$, JIB-04 $IC_{50}$, and JIB-04 2x $IC_{50}$ negatively enriched gene sets related to cell cycle processes, chromosomal arrangement, protein modification, cellular localization, and response to stimuli. Gene sets relating to cellular responses were positively enriched by JIB-04 2x $IC_{50}$.

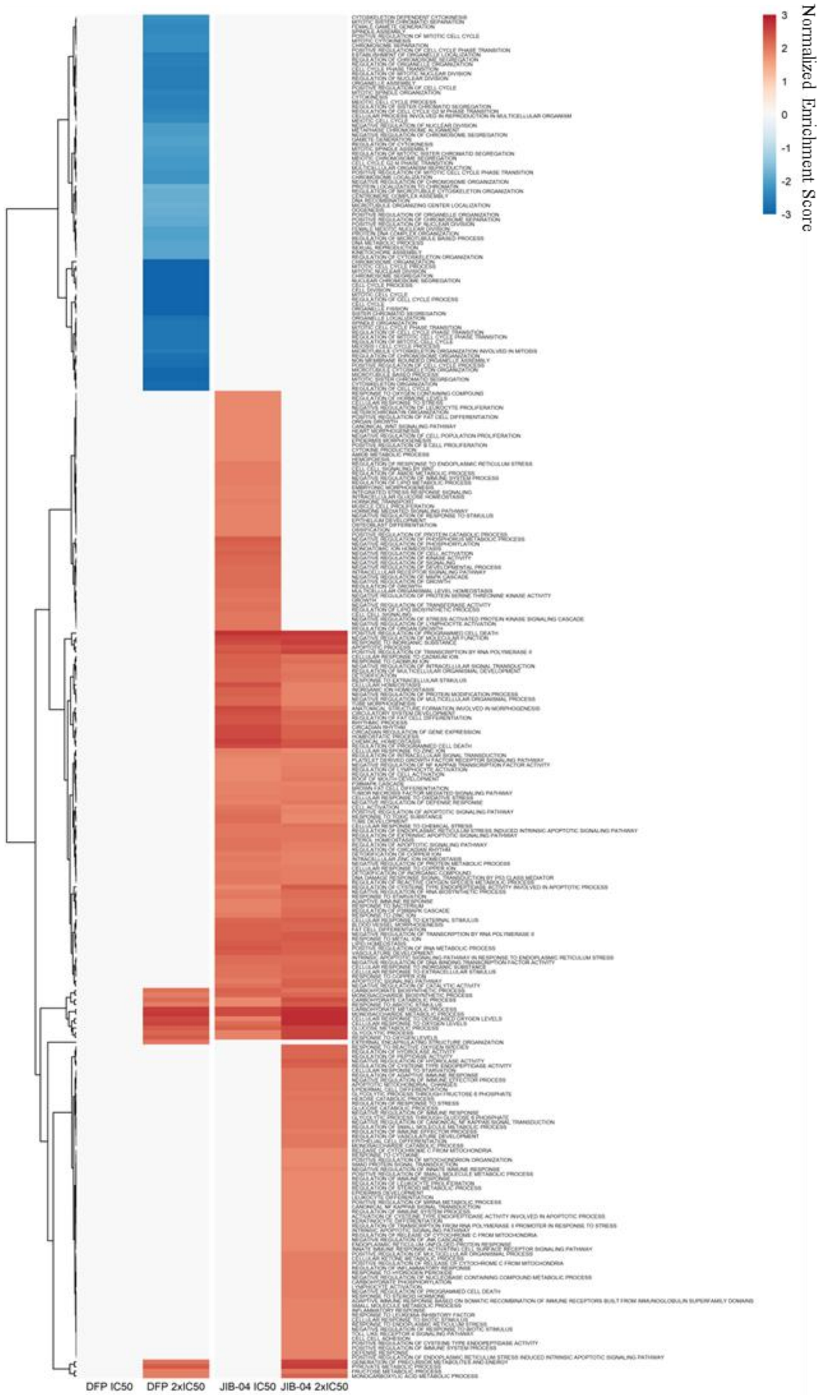


**Figure S3.** Full Gene Ontology Biological Processes (GOBP) Gene Set Enrichment Analysis (GSEA) heatmap of DFP and JIB-04 treatment to MCF-7 cells. Significant gene set enrichment was defined by $p < 0.01$ and false discovery rate (FDR) < 0.10. DFP $IC_{50}$ did not significantly enrich GOBP gene sets. DFP 2x $IC_{50}$, JIB-04 $IC_{50}$, and JIB-04 2x $IC_{50}$ positively enriched gene sets related to oxygen level response and metabolic processes. Individually, DFP 2x $IC_{50}$ negatively enriched gene sets related to cell cycle processes and chromosomal arrangement, JIB-04 at $IC_{50}$ and 2x $IC_{50}$ positively enriched gene sets related to cellular responses to metal ions, apoptotic and tumor necrotic factor signaling pathways.

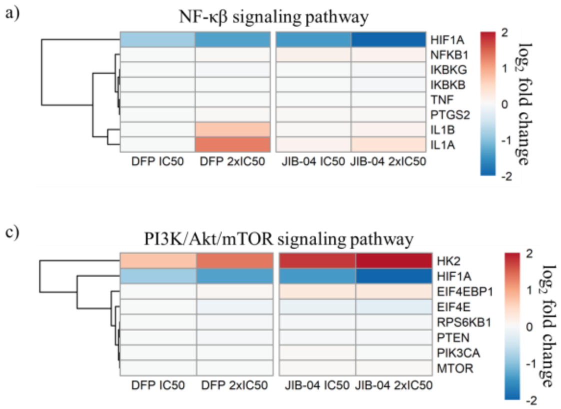


**Figure S4.** Effects of DFP and JIB-04 on traditional hypoxia inducible factor 1α (HIF-1α) regulation pathways were minimally affected by the downregulation of HIF-1α a) NF-κβ signaling pathway, b) RAS-RAF-MEK-ERK pathway, c) PI3K/Akt/mTOR signaling pathway, and d) JAK-STAT3 pathway.

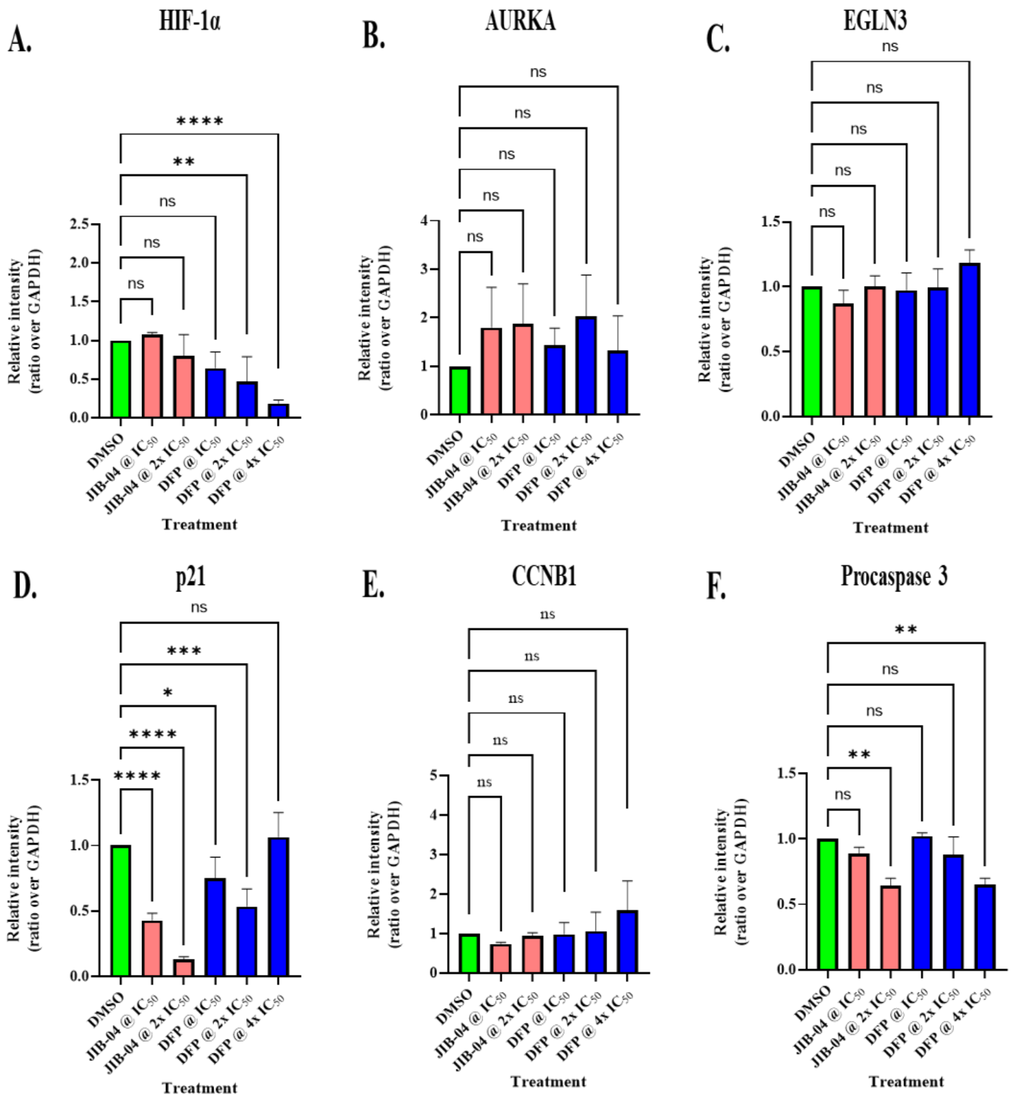


**Figure S5.** Western blot densitometric analyses (A-F) of MCF-7 cells treated with JIB-04 and DFP for 24 h. Quantification bars display means plus standard deviations; ordinary one-way ANOVA of each treatment was compared with the DMSO control group. **p* < 0.05; ***p* < 0.0048; ****p* = 0.0001; *****p* < 0.0001.

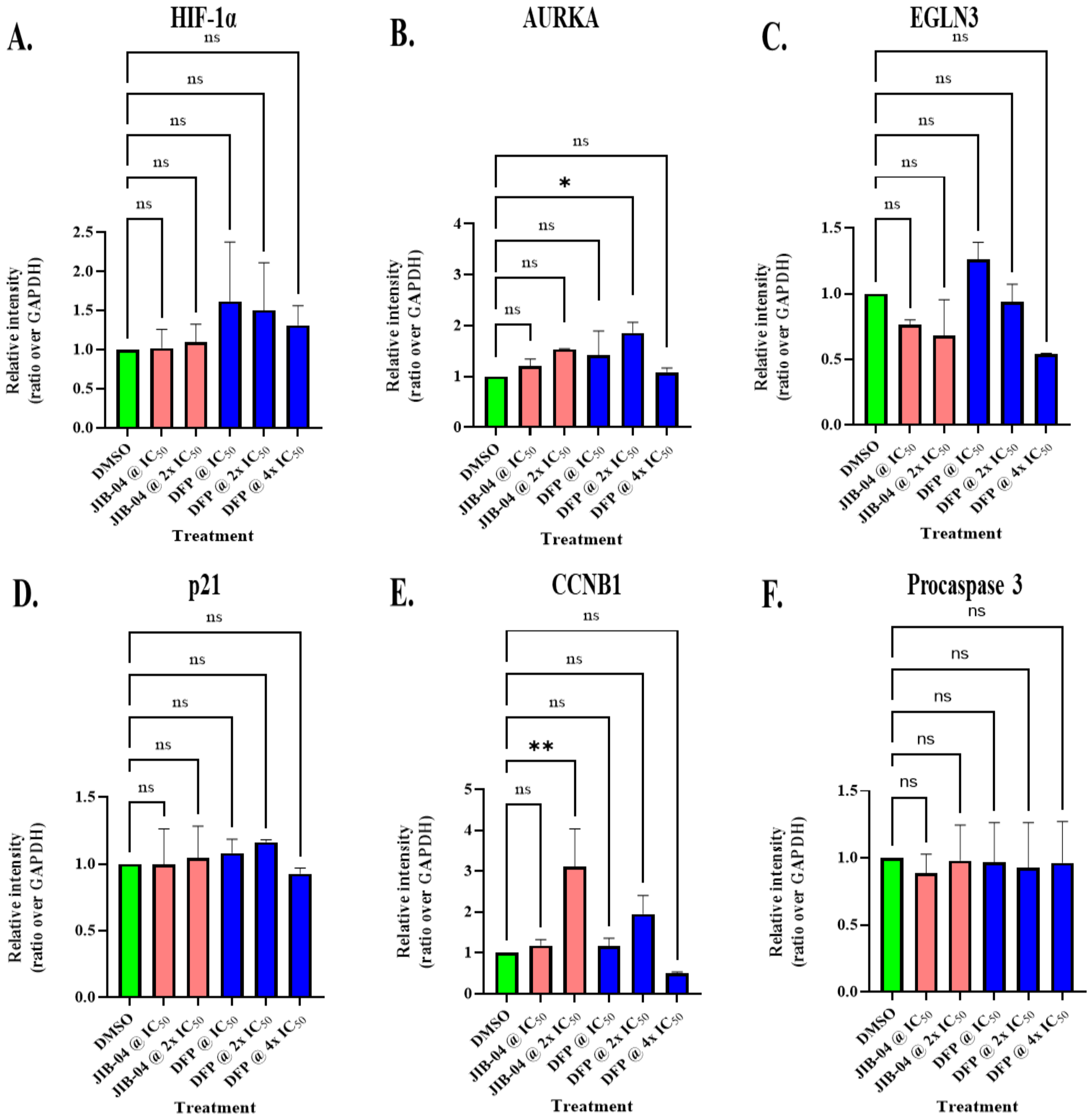


**Figure S6.** Western blot densitometric analyses (A-F) of MDA-MB-231 cells treated with JIB-04 and DFP for 24 h. Quantification bars display means plus standard deviations; ordinary one-way ANOVA of each treatment was compared with the DMSO control group. *$p < 0.05$; **$p < 0.001$.

**Full Western blot gels with labels**

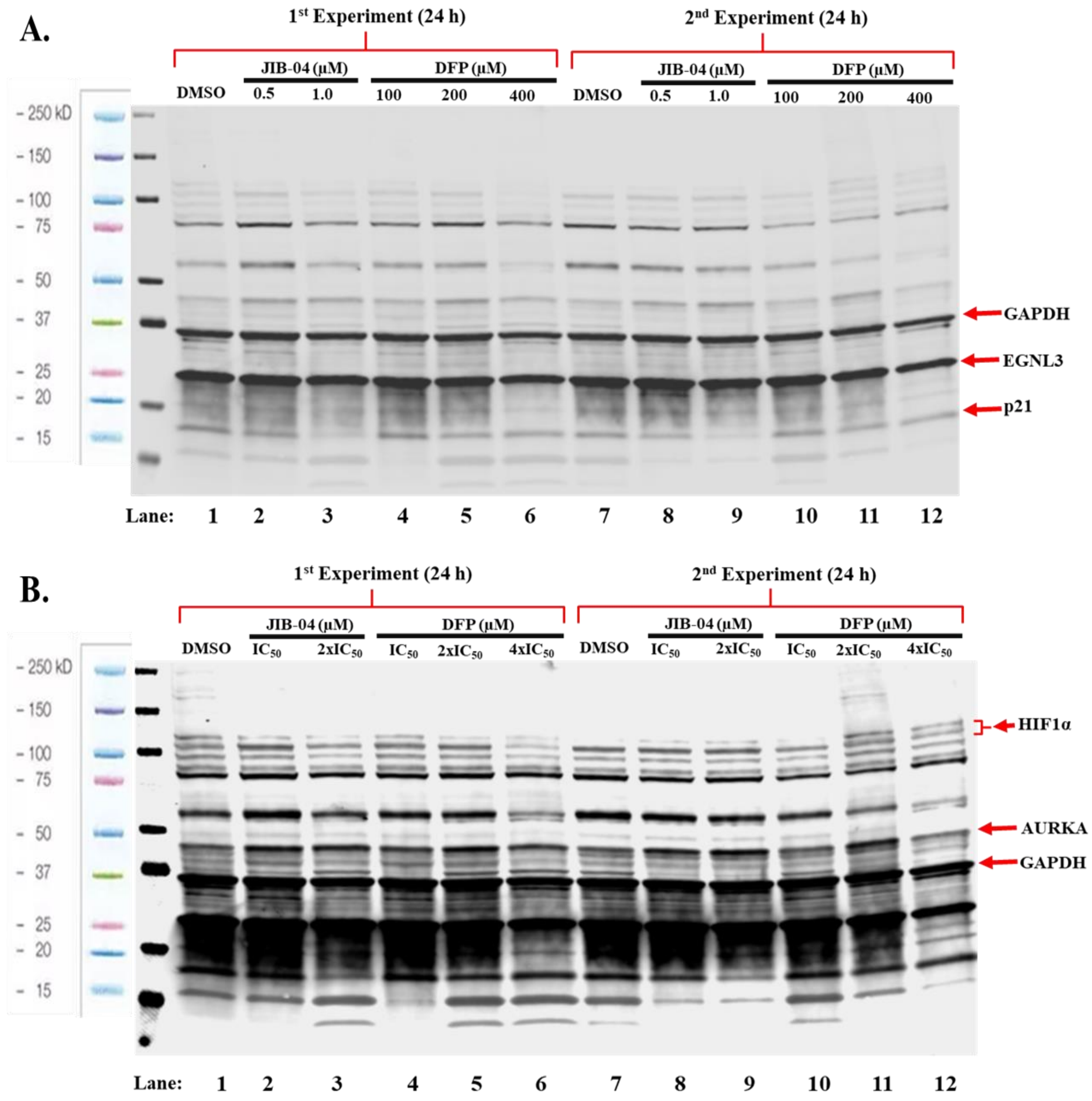


**Figure S7.** Full immunoblot gel images (A-B) of MCF-7 cells treated with JIB-04 and DFP for 24 h: *HIF1α antibody cocktail.*

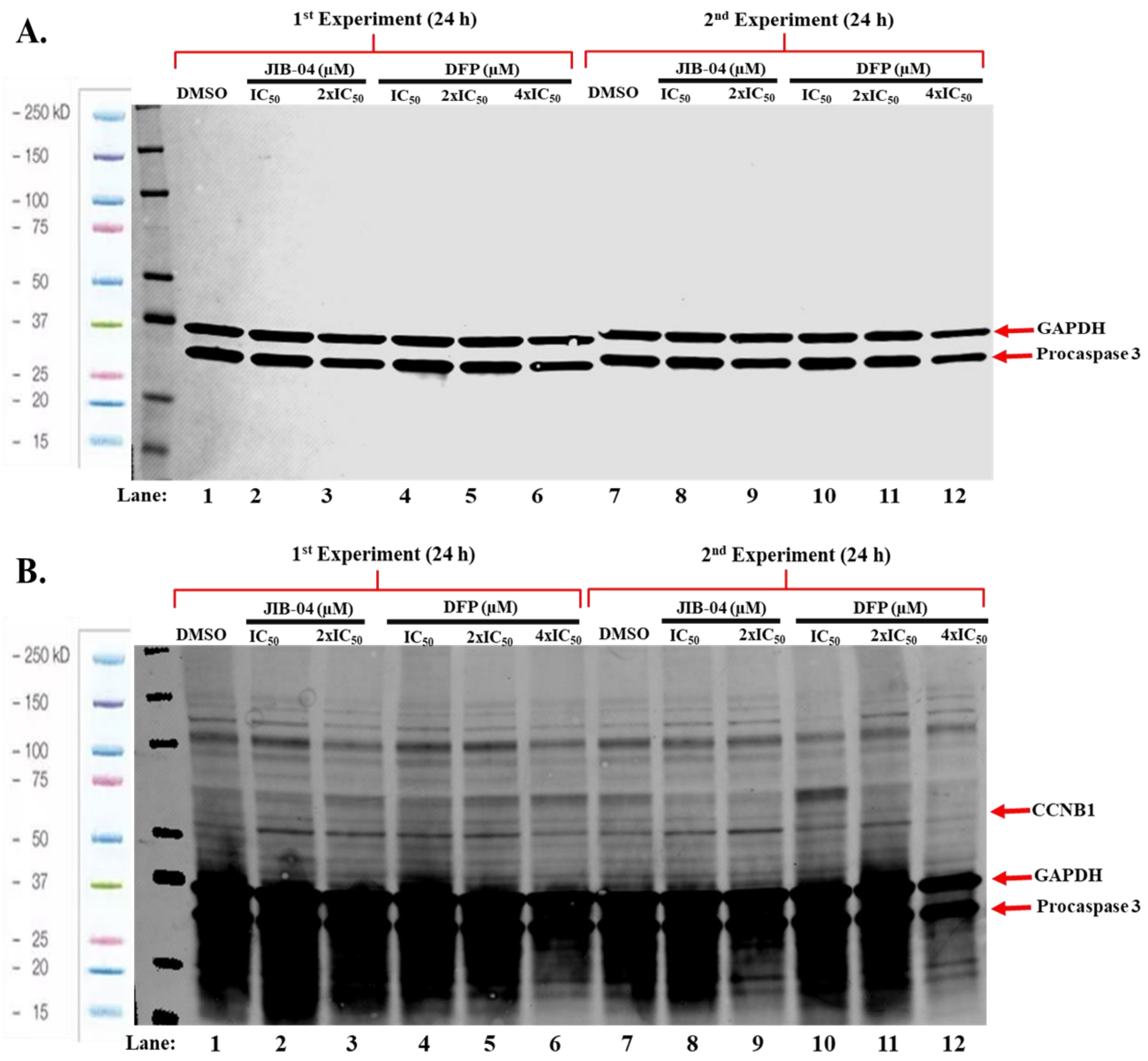


**Figure S8.** Full immunoblot gel images (A-B) of MCF-7 cells treated with JIB-04 and DFP for 24 h: *CCNB1 antibody cocktail.*

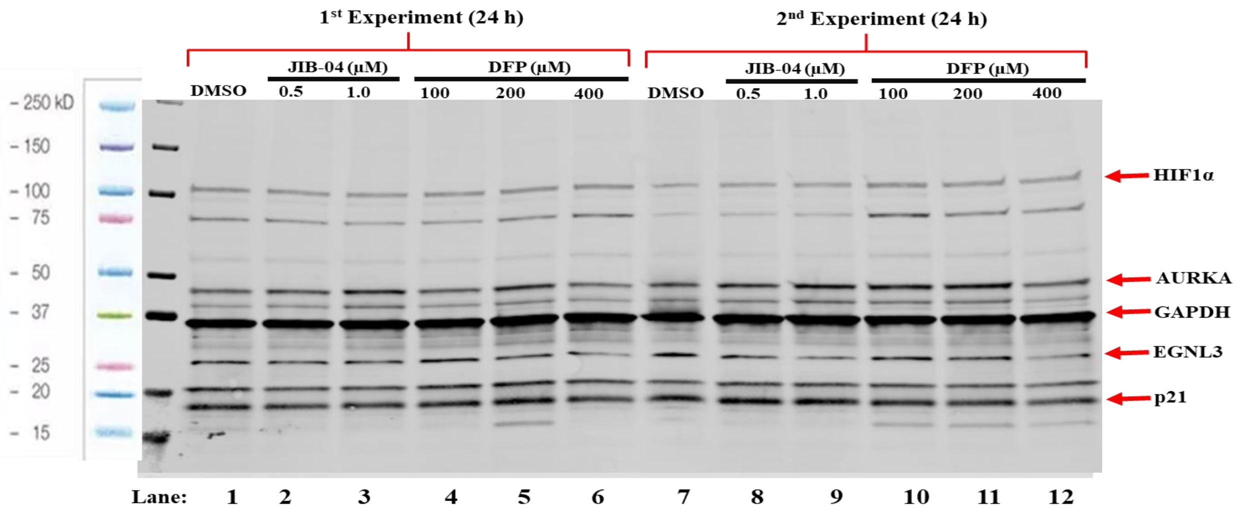


**Figure S9.** Full immunoblot gel images of MDA-MB-231 cells treated with JIB-04 and DFP for 24 h: *HIF1α antibody cocktail.*

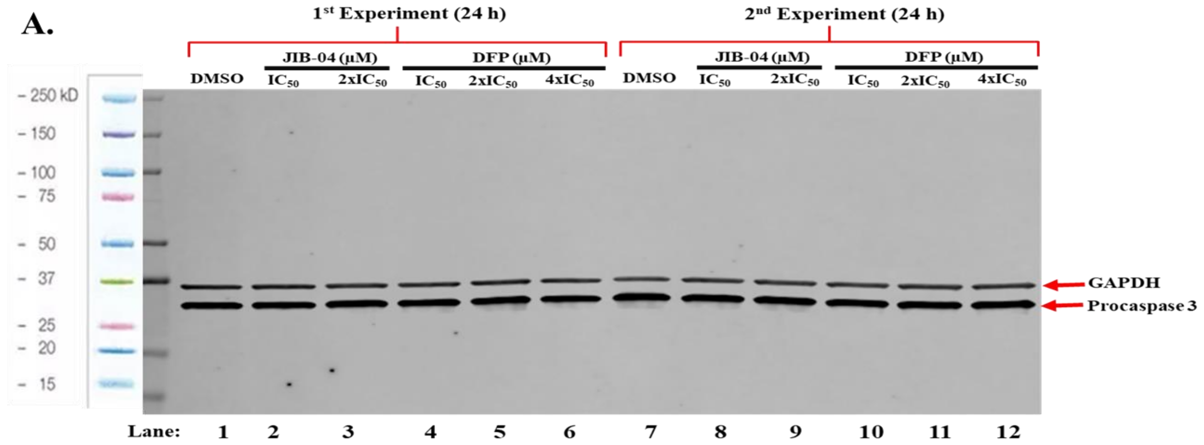


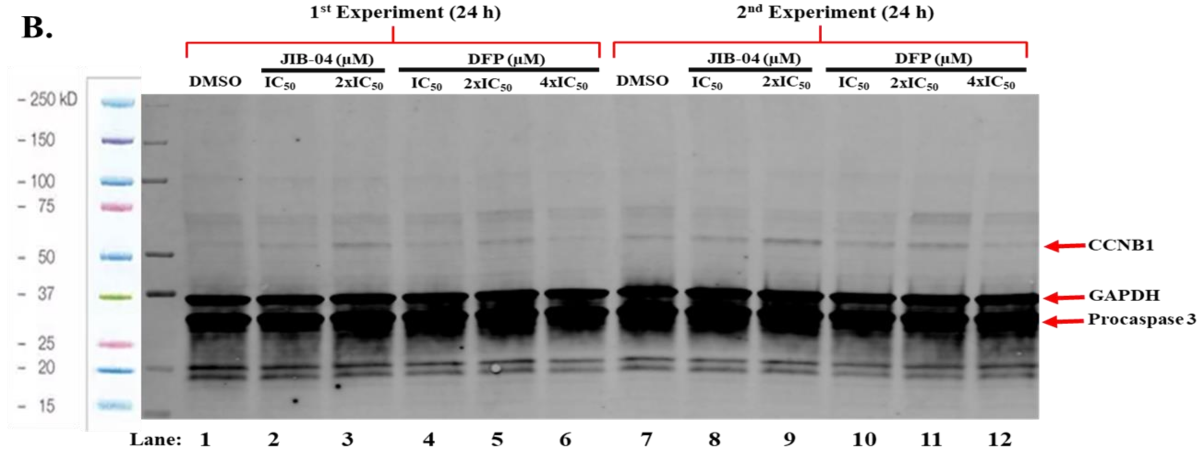


**Figure S10.** Full immunoblot gel images (A-B) of MDA-MB-231 cells treated with JIB-04 and DFP for 24 h: *CCNB1 antibody cocktail.*

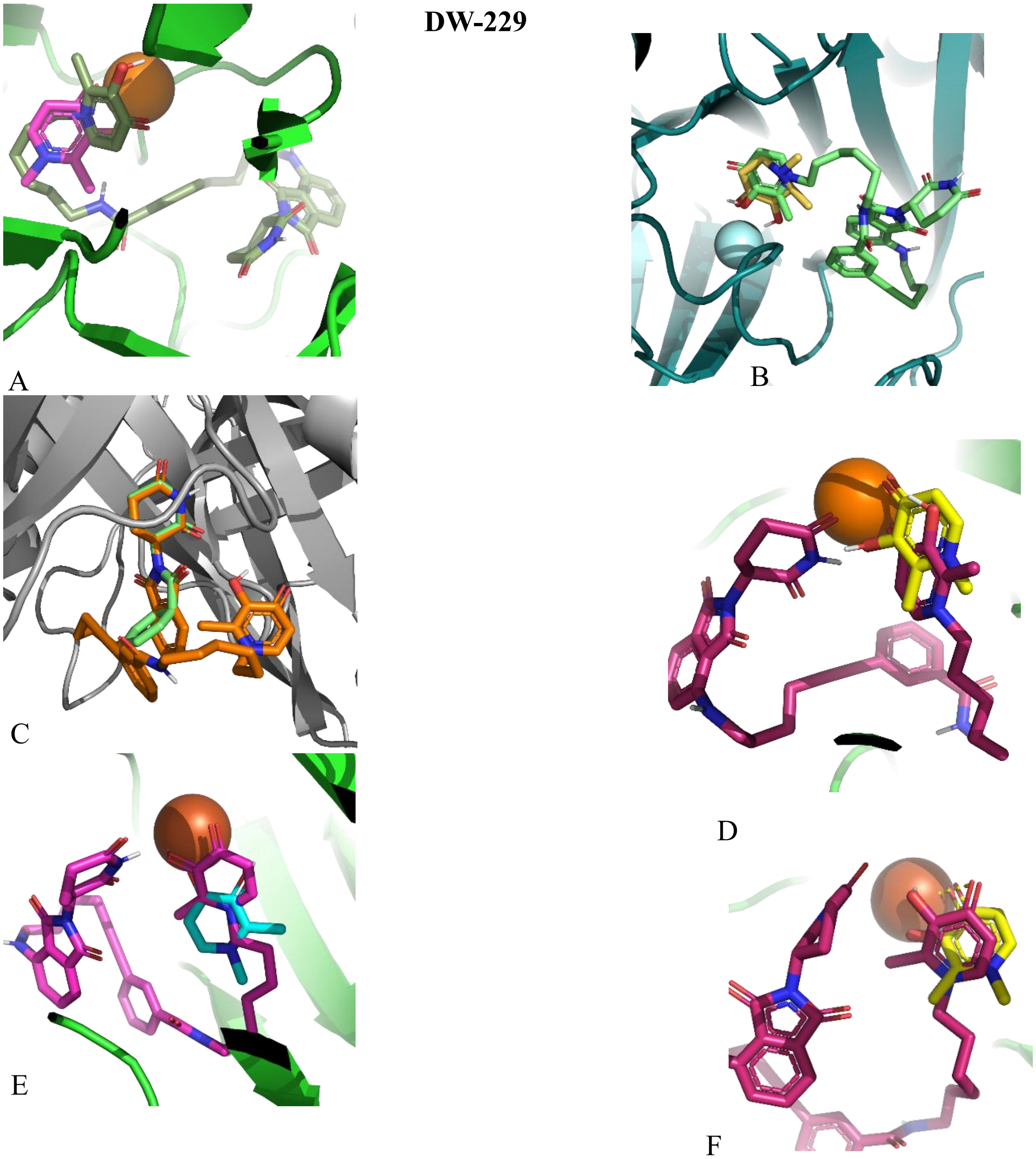


**Figure S11.** Molecular docking results of **DW-229** compared with DFP and thalidomide. (A) E3 Ligase (PDB: 4CI1) overlaying **DW-229** (orange, -9.507 kcal/mol) and thalidomide (green, -9.51 kcal/mol) show a large consistency in overlap between the active components of thalidomide. (B) KDM6A (PDB:3AVR) **DW-229** (green, -8.891 kcal/mol) DFP (yellow, -4.972 Kcal/mol) both show orientation towards the Fe (II) ion and are a plausible distance for chelation. (C) KDM5A (PDB:5CEH) **DW-229** (pink, -10.576 kcal/mol) and DFP (blue, -5.067 kcal/mol) demonstrate plausible coordination and potential chelation of the Fe (II) ion. (D) KDM5B (PDB:6H4Z) **DW-229** (Maroon, -10.821) and DFP (yellow-5.387 kcal/mol) both have their DFP moiety relatively close to the Fe (II) ion, but **DW-229** is flipped 180°. (E) KDM3A (PDB:2Q8C) **DW-229** (maroon, -10.644 kcal/mol ) and DFP (yellow, -5.568 kcal/mol) both have the DFP moiety plausibly close to the Fe (II) ion for chelation. (F) KDM2A (PDB:4QXB) **DW-229** (green, -9.479 kcal/mol) and DFP (pink, -5.435 kcal/mol), the position of **DW-229** DFP moiety seems not be well-oriented to optimally chelate the Fe (II) ion.

**DW-449**

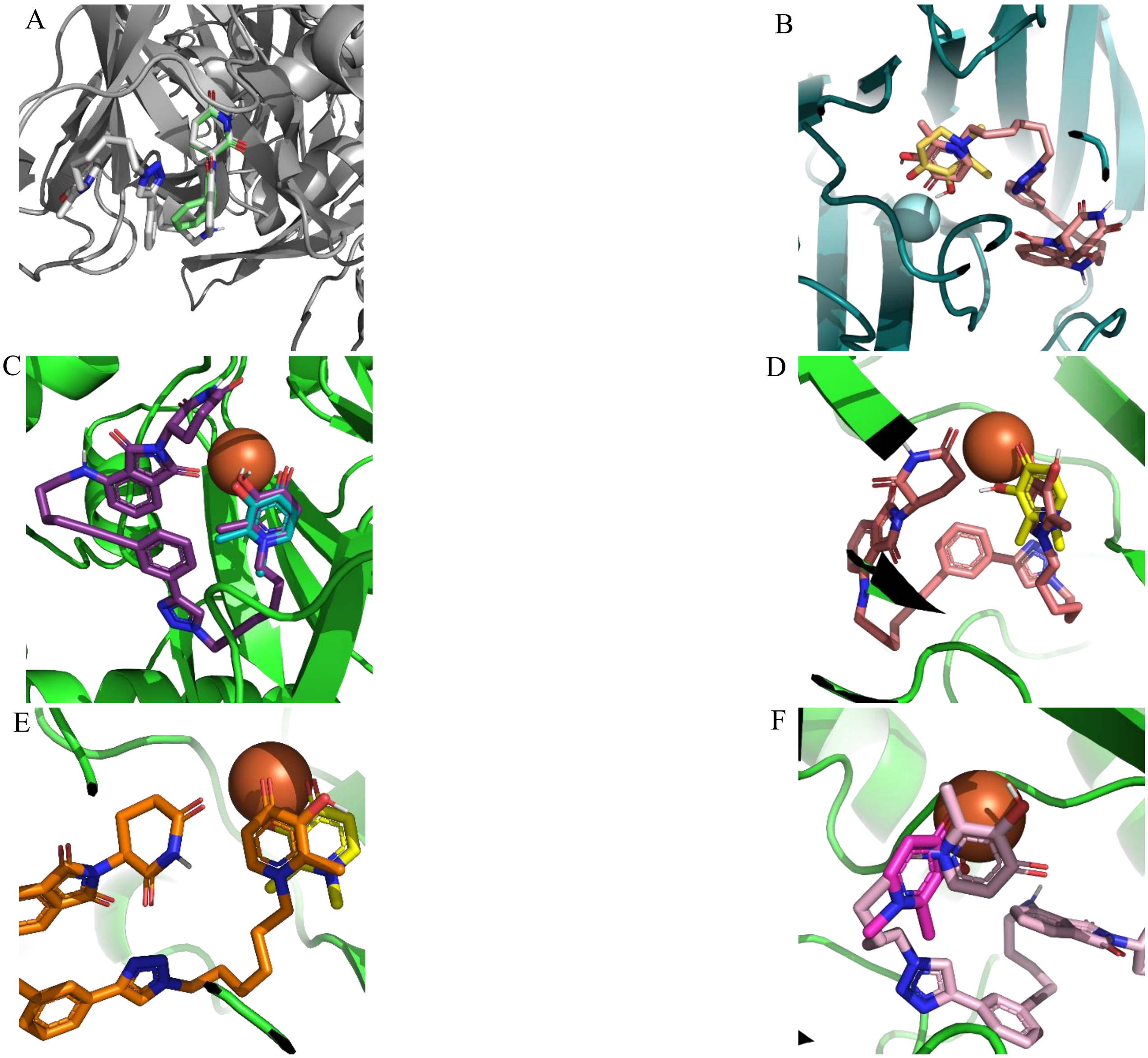


**Figure S12.** Molecular docking results of **DW-449** compared with DFP and thalidomide. (A) E3 Ligase (PDB: 4CI1) overlaying **DW-449** (white, -9.438 kcal/mol) and thalidomide (green, -9.51 kcal/mol) show a large consistency in overlap between the active components of thalidomide. **(**B) KDM6A (PDB:3AVR) **DW-449** (salmon, -9.773 kcal/mol) and DFP (yellow, -4.972 Kcal/mol) both show strong orientation towards the iron ion and are a plausible distance for chelation. (C) KDM5A (PDB:5CEH) **DW-449** (purple, -10.498 kcal/mol) and DFP (teal, -5.067 kcal/mol), the docked output shows a large degree of overlap between DFP and the corresponding portion of **DW-449**, with plausible chelation. (D) KDM5B (PDB:6H4Z) docked output of **DW-449 (**salmon, -10.707 kcal/mol) and DFP (yellow, -5.387 kcal/mol) shows that their iron-binding groups are relatively close to the Fe (II) ion, although **DW-449** is not oriented for optimal chelation. (E) KDM3A (PDB:2Q8C) **DW-449** (orange, -9.859 kcal/mol) and DFP (yellow, -5.568 kcal/mol) both adopt orientations showing their iron-binding groups are plausibly close to the Fe (II) ion but not oriented for optimal chelation. (F) KDM2A (PDB:4QXB) **DW-449 (**light pink, -10.768 kcal/mol) and DFP (dark pink, -5.435 kcal/mol) both adopt orientations showing their iron-binding groups are plausibly close to the Fe (II) ion but not oriented for optimal chelation.

**DW-451**

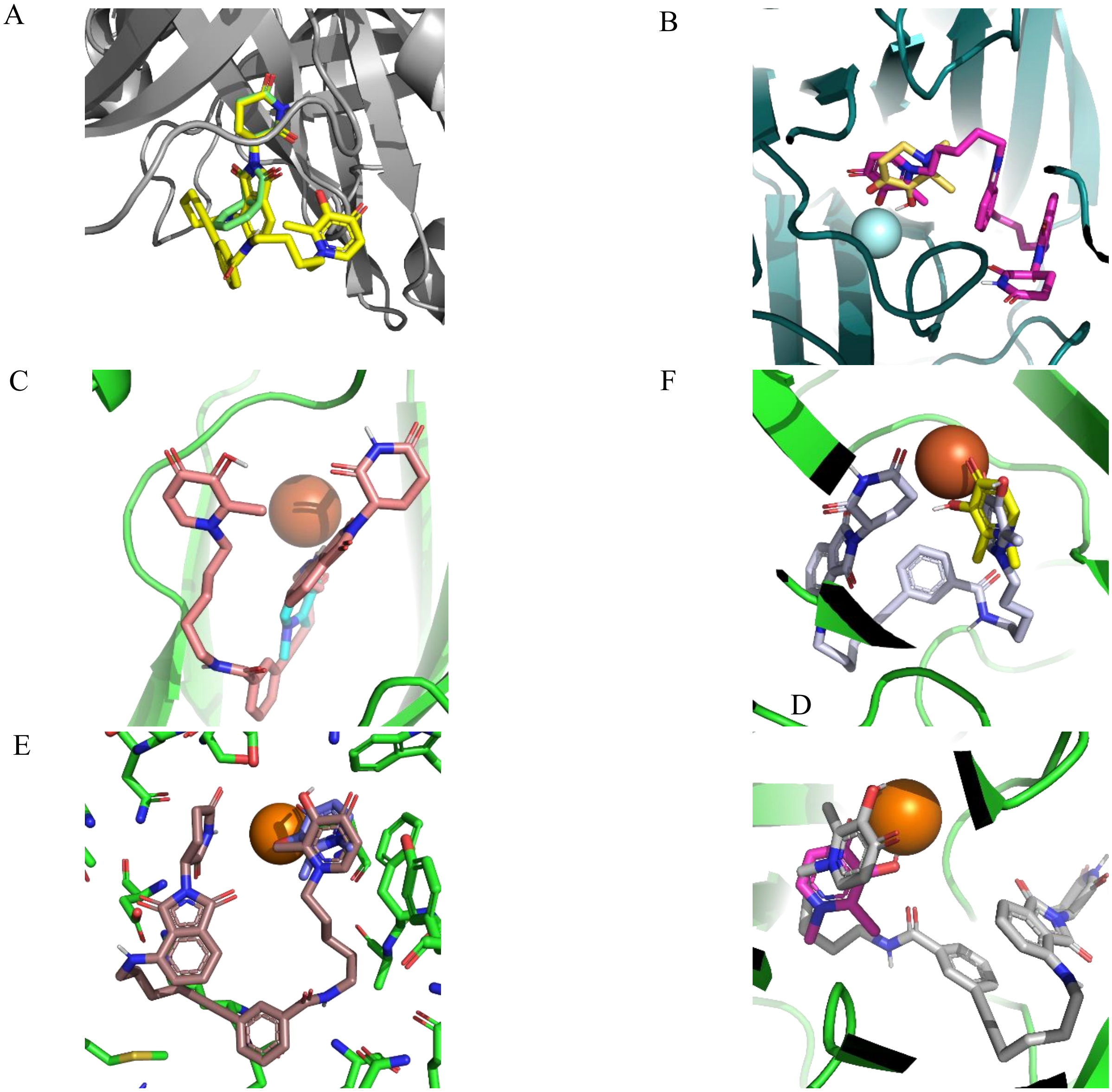


**Figure S13.** Molecular docking results of **DW-451** compared with DFP and thalidomide. (A) E3 Ligase (PDB: 4CI1) overlaying **DW-451** (yellow, -10.026 kcal/mol) and thalidomide (green, -9.51 kcal/mol) show a large consistency in overlap between the active components of thalidomide. (B)

KDM6A (PDB:3AVR) **DW-451** (pink, -8.143 kcal/mol) and DFP (yellow, -4.972 kcal/mol) both show strong orientation towards the Fe (II) ion and are at a plausible distance for chelation. **(**C) KDM5A (PDB:5CEH) **DW-451** (salmon, -10.945 kcal/mol) and DFP (blue, -5.067 kcal/mol) adopt docked orientations showing that their iron-binding groups are in proximity for plausible chelation with the Fe (II) ion. However, these orientations may not afford optimal chelation. **(**D) KDM5B (PDB:6H4Z) **DW-451** (gray, -11.186 kcal/mol) and DFP (yellow, -5.387 kcal/mol) adopt docked orientations showing that their iron-binding groups are in proximity for plausible chelation with the Fe (II) ion. However, **DW-451** is flipped from chelation. (E) KDM3A (PDB:2Q8C) **DW-451** (brown, -10.379 kcal/mol) and DFP (blue, -5.568 kcal/mol) the iron-binding group of **DW-451** is close to the Fe (II) ion, but not optimally oriented for chelation. (F) KDM2A (PDB:4QXB) **DW-451** (grey, -9.765 kcal/mol) and DFP (pink, -5.435 kcal/mol) the iron-binding group of **DW-451** is close to the Fe (II) ion, but not optimally oriented for chelation.

**DW-455**

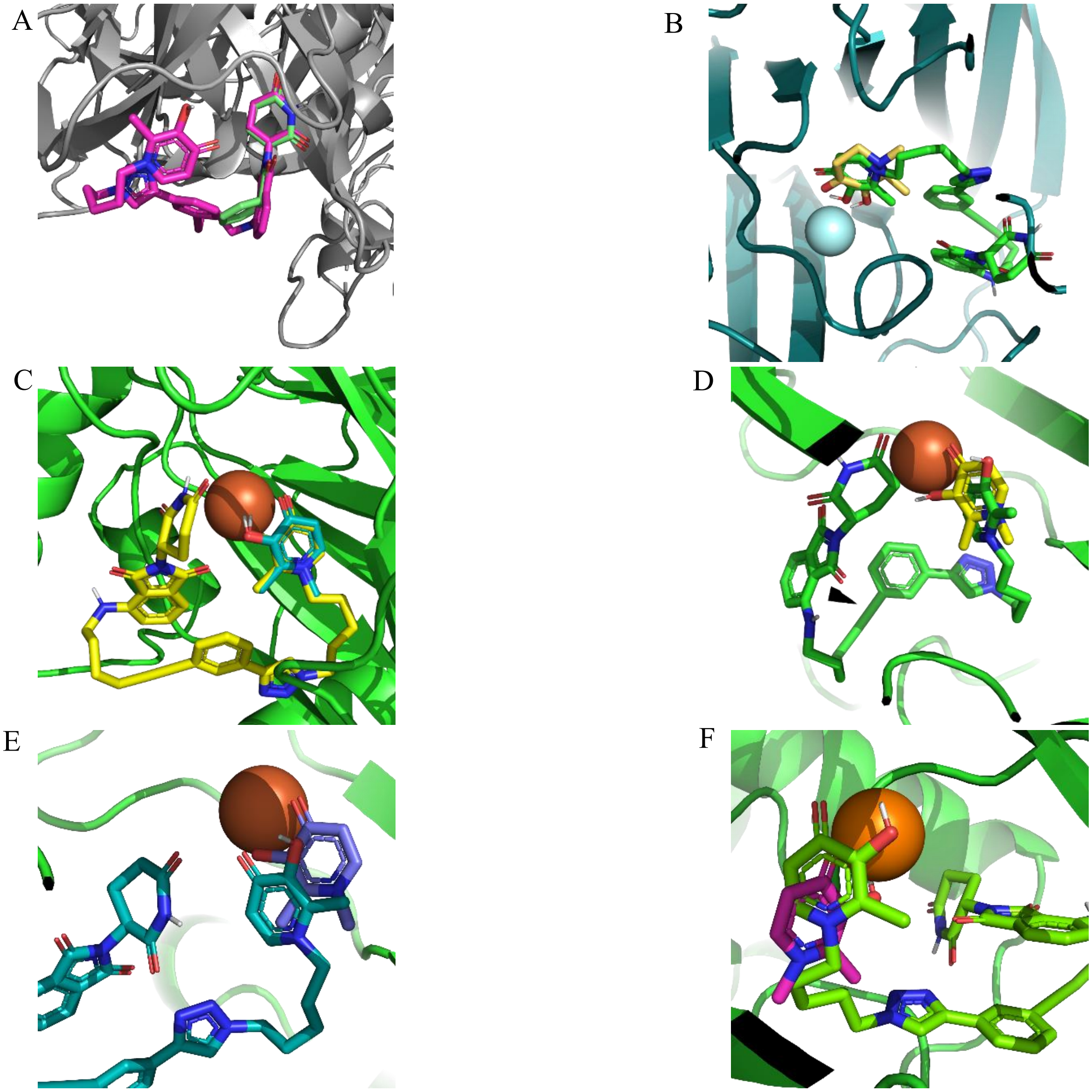

**Figure S14.** Molecular docking results of **DW-455** compared with DFP and thalidomide. (A) E3 Ligase (PDB: 4CI1) overlaying **DW-455** (pink, -10.675 kcal/mol) and thalidomide (green, -9.51 kcal/mol) show a large consistency in overlap between the active components of thalidomide. (B) KDM6A (PDB:3AVR) **DW-455** (green, -9.018 kcal/mol) and DFP (yellow, -4.972 kcal/mol) both show strong orientation towards the Fe (II) ion and are at a plausible distance for chelation. (C) KDM5A (PDB:5CEH) **DW-455** (yellow, -10.11 kcal/mol) and DFP (blue, -5.067 kcal/mol) the docked output shows a large degree of overlap between DFP and the corresponding portion of **DW-455**. (D) KDM5B (PDB:6H4Z) **DW-455** (green, -10.112 kcal/mol) and DFP (yellow, -5.387 kcal/mol) both adopt docking poses such that their iron binding groups are oriented toward the Fe (II) ion, but **DW-455** is not oriented for optimal chelation. (E) KDM3A (PDB:2Q8C) **DW-455** (teal, -10.395 kcal/mol) and DFP (blue, -5.568 kcal/mol) iron binding groups are oriented toward the Fe (II) ion and are within a distance for chelation. (F) KDM2A (PDB:4QXB) **DW-455** (green, -9.665 kcal/mol) and DFP (pink, -5.435 kcal/mol) position their iron binding groups close to the Fe (II) ion. However, **DW-455** iron binding group is not oriented for optimal chelation.

**Table S2.** Docking scores of the designed CRBN E3 ligase DFP-based PROTACs.

| **Protein (PDB)** | **Docking Scores Kcal/mol)** | | | | | |
|---|---|---|---|---|---|---|
| | CRBN | DFP | **DW-229** | **DW-449** | **DW-451** | **DW-455** |
| CRBN (4CI1) | -9.51 | | -9.507 | -9.438 | -10.026 | -10.675 |
| KDM6A (3AVR) | | -4.972 | -8.891 | -9.773 | 8.143 | -9.018 |
| KDM5A (5CEH) | | -5.067 | -10.576 | -10.498 | -10.945 | -10.11 |
| KDM5B (6H4Z) | | -5.387 | -10.821 | -10.707 | -11.186 | -10.112 |
| KDM3A (2Q8C) | | -5.568 | -10.664 | -9.859 | -10.379 | -10.395 |
| KDM2A (4QXB) | | 5.435 | -9.479 | -10.768 | -9.765 | -9.665 |

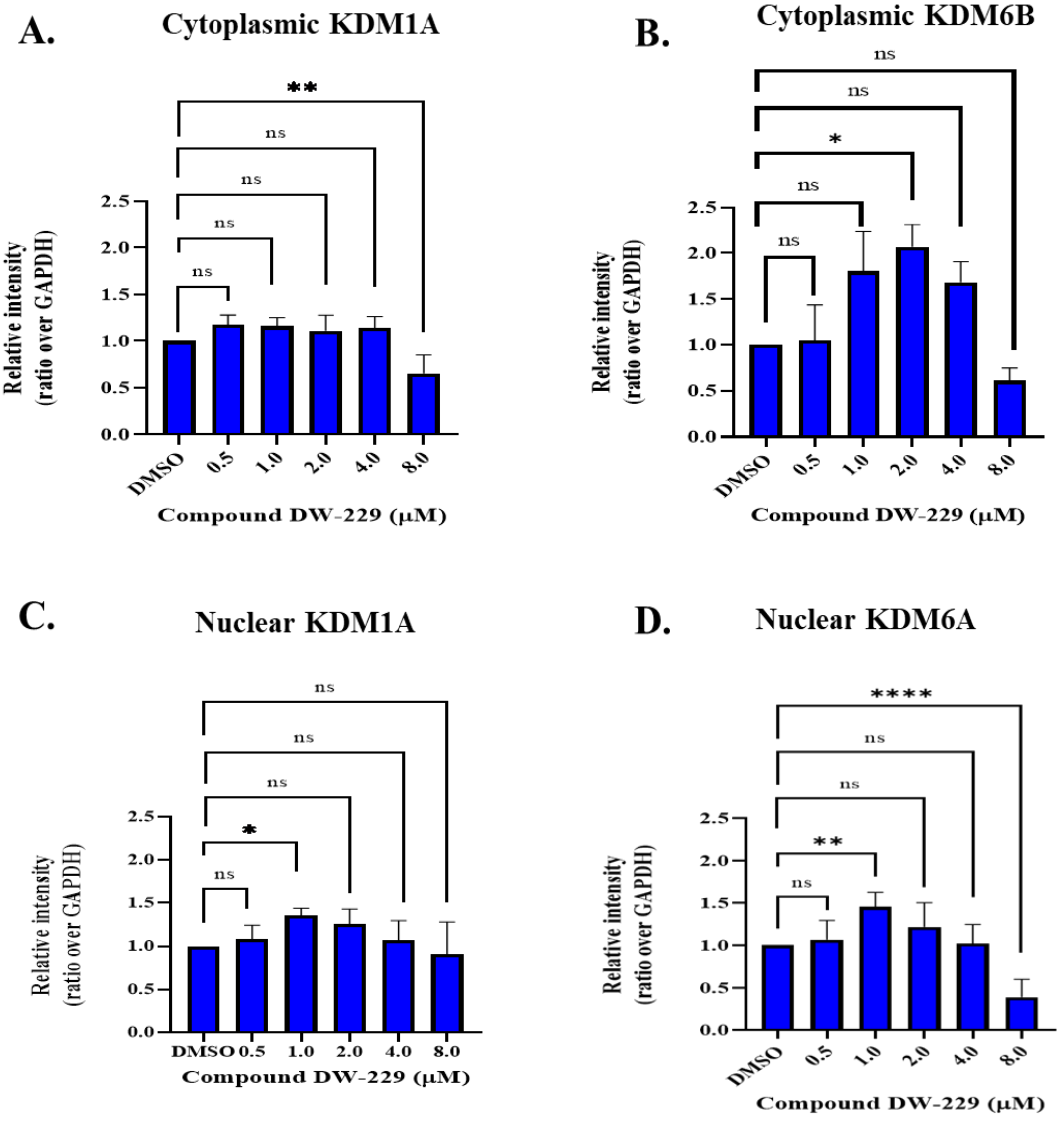


**Figure S15.** Western blot densitometric analyses (A–D) of MCF-7 cells treated with **DW-229** for 24 h: *KDM1A, KDM6A, and KDM6B antibody cocktail.* Quantification bars display means plus standard deviations of data obtained from at least two independent experiments; ordinary one-way ANOVA of each treatment was compared with the DMSO control group. *$p < 0.05$; **$p < 0.005$; ***$p = 0.001$; ****$p < 0.0001$).

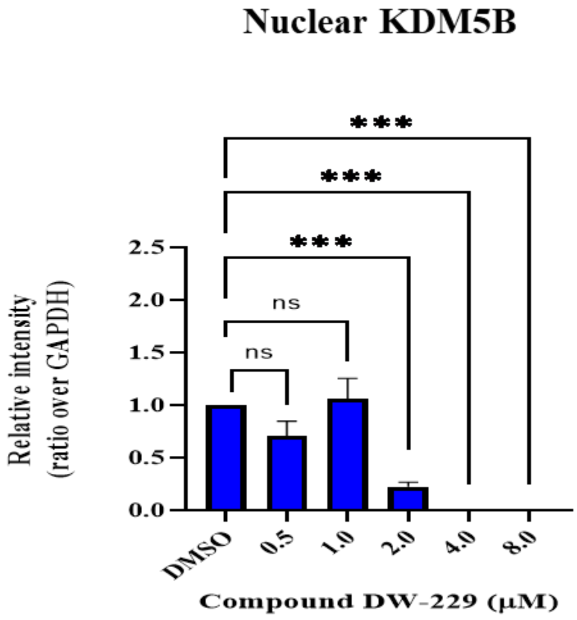


**Figure S16.** Western blot densitometric analysis of MCF-7 cells treated with **DW-229** for 24 h: *KDM5B antibody cocktail.* Quantification bars display means plus standard deviations of data obtained from at least two independent experiments; ordinary one-way ANOVA of each treatment was compared with the DMSO control group. ***$p$ = 0.001.

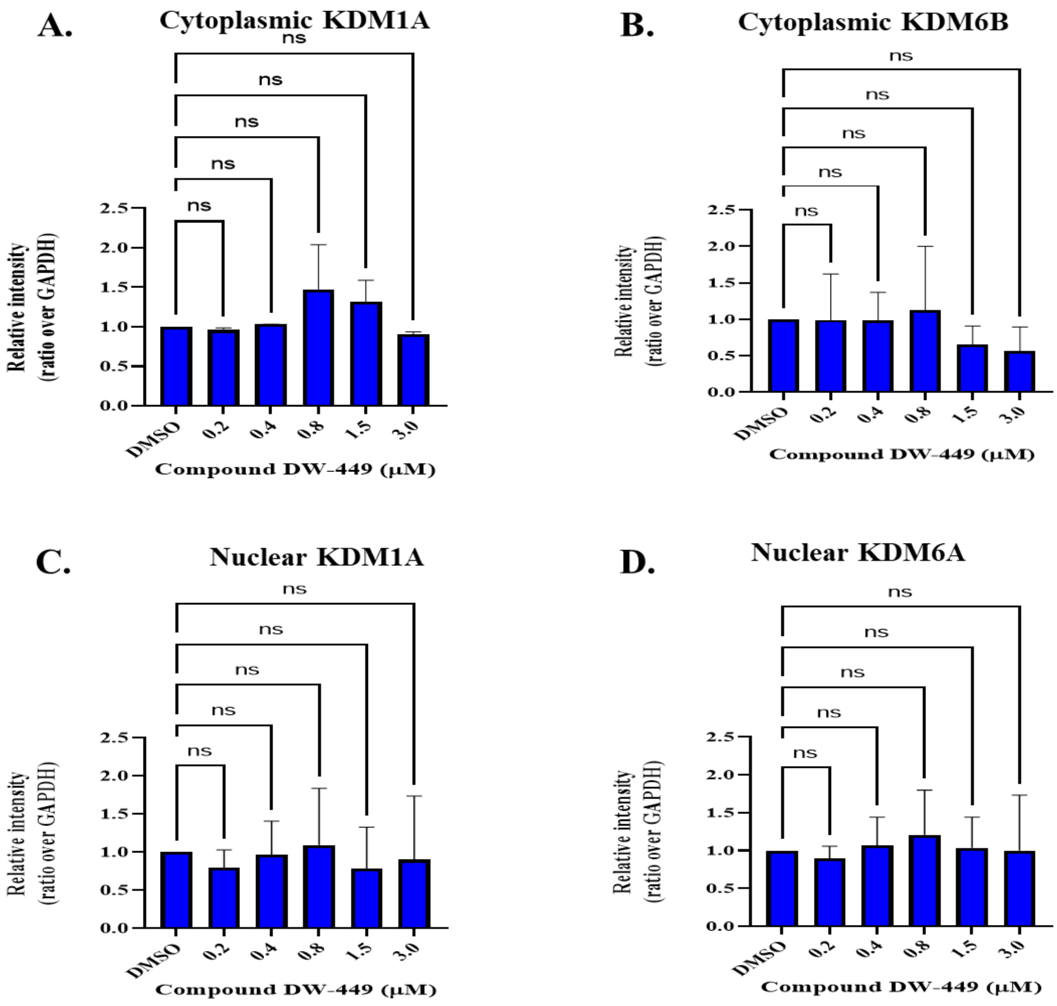


**Figure S17.** Western blot densitometric analyses (A–D) of MCF-7 cells treated with **DW-449** for 24 h: *KDM1A, KDM6A, and KDM6B antibody cocktail.* Quantification bars display means plus standard deviations of data obtained from at least two independent experiments; ordinary one-way ANOVA of each treatment was compared with the DMSO control group.

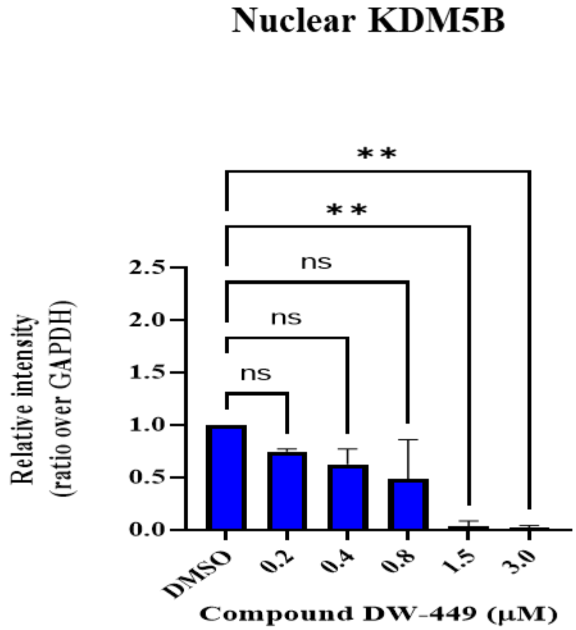


**Figure S18.** Western blot densitometric analysis of MCF-7 cells treated with **DW-449** for 24 h: *KDM5B antibody cocktail.* Quantification bars display means plus standard deviations of data obtained from at least two independent experiments; ordinary one-way ANOVA of each treatment was compared with the DMSO control group. $^{**}p < 0.005$.

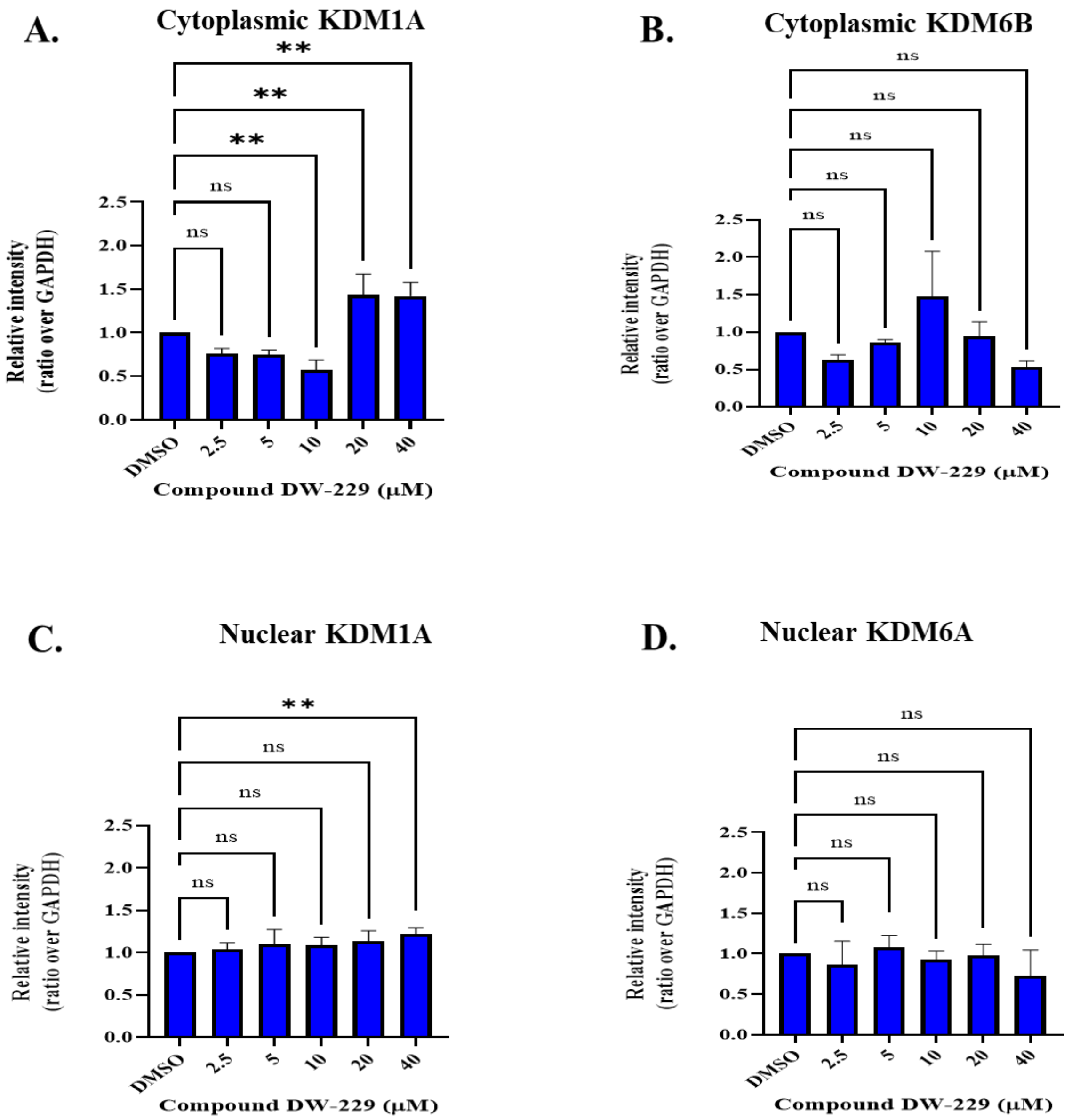


**Figure 19.** Western blot densitometric analyses (A–D) of MDA-MB-231 cells treated with **DW-229** for 24 h: *KDM1A, KDM6A, and KDM6B antibody cocktail.* Quantification bars display means plus standard deviations of data obtained from at least two independent experiments; ordinary one-way ANOVA of each treatment was compared with the DMSO control group. $^{**}p < 0.005$.

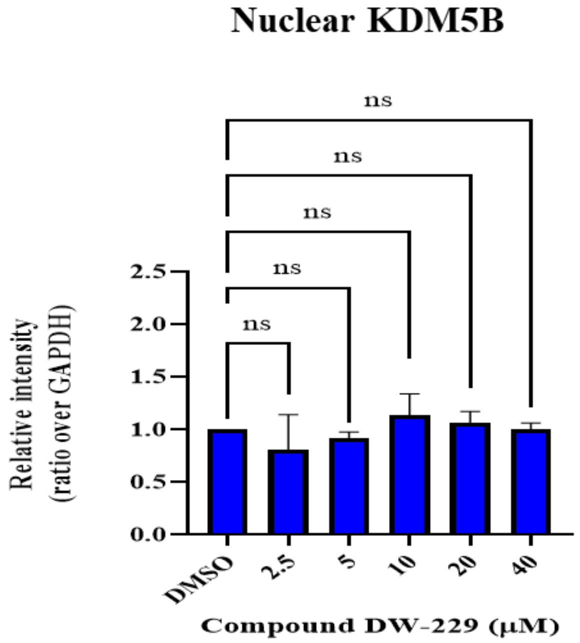


**Figure 20.** Western blot densitometric analysis of MDA-MB-231 cells treated with **DW-229** for 24 h: *KDM5B antibody cocktail.* Quantification bars display means plus standard deviations of data obtained from at least two independent experiments; ordinary one-way ANOVA of each treatment was compared with the DMSO control group.

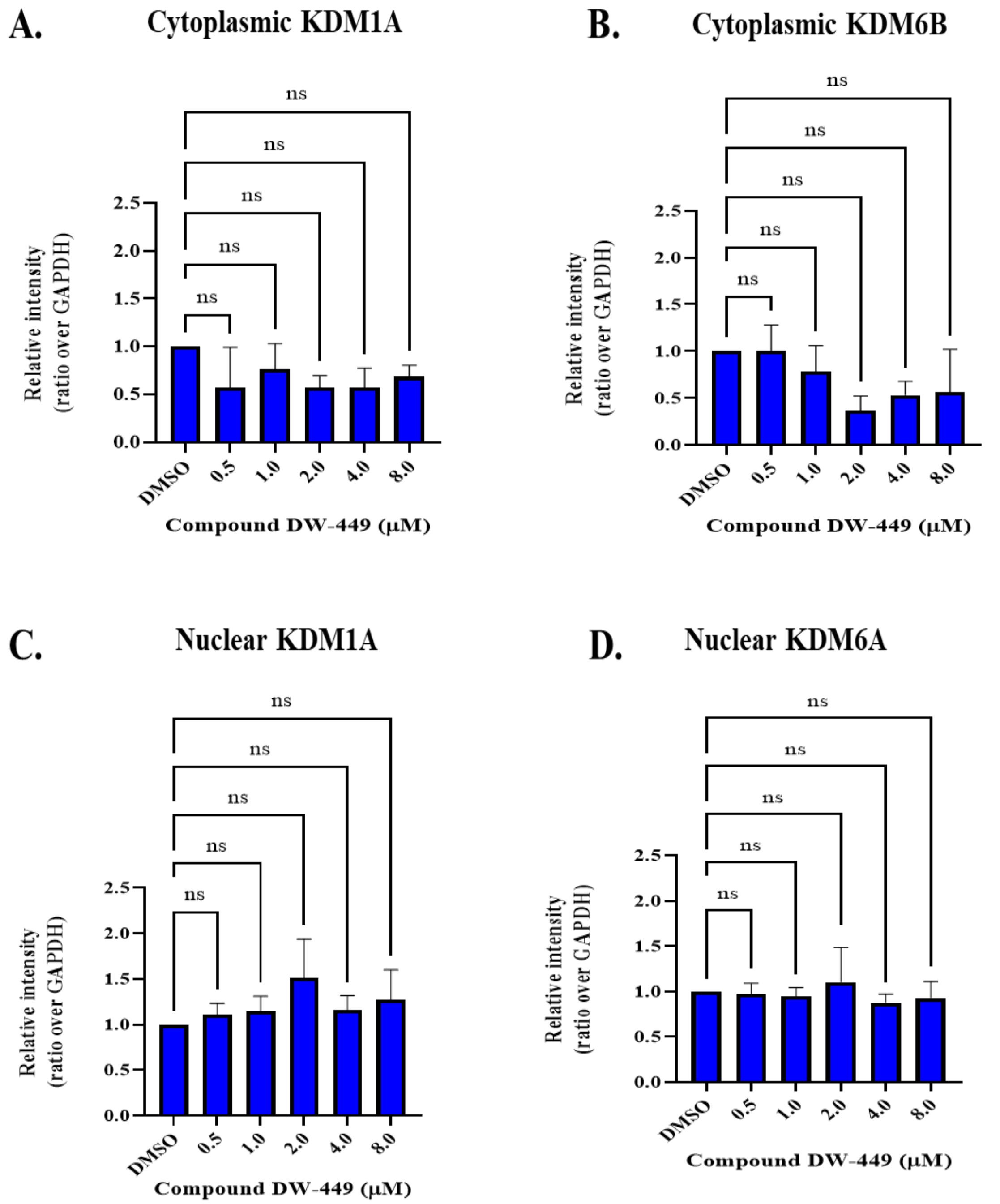


**Figure 21.** western blot densitometric analysis (A–D) of MDA-MB-231 cells treated with **DW-449** for 24 h: *KDM1A, KDM6A, and KDM6B antibody cocktail.* Quantification bars display means plus standard deviations of data obtained from at least two independent experiments; ordinary one-way ANOVA of each treatment was compared with the DMSO control group.

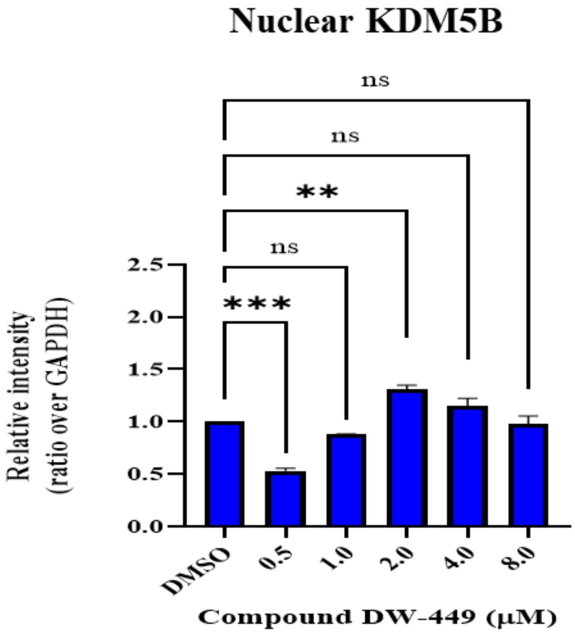


**Figure 22.** Western blot densitometric analysis of MDA-MB-231 cells treated with **DW-449** for 24 h: *KDM5B antibody cocktail.* Quantification bars display means plus standard deviations of data obtained from at least two independent experiments; ordinary one-way ANOVA of each treatment was compared with the DMSO control group. $^{**}p < 0.005$; $^{***}p = 0.001$.

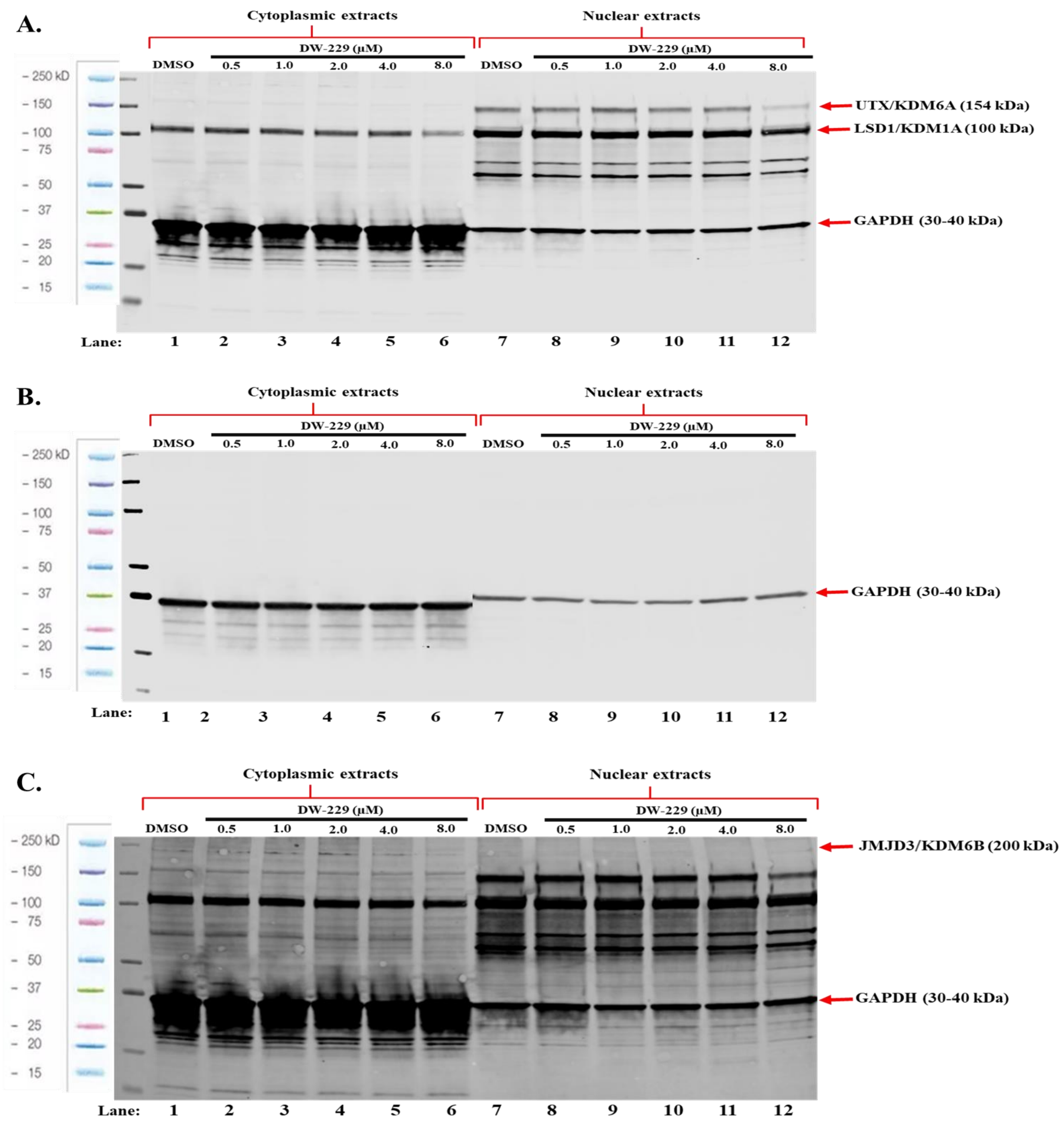


**Figure S23.** Full immunoblot gel images (A-C) of MCF-7 cells treated with **DW-229** for 24 h: *KDM1A, KDM6A, and KDM6B antibody cocktail.*

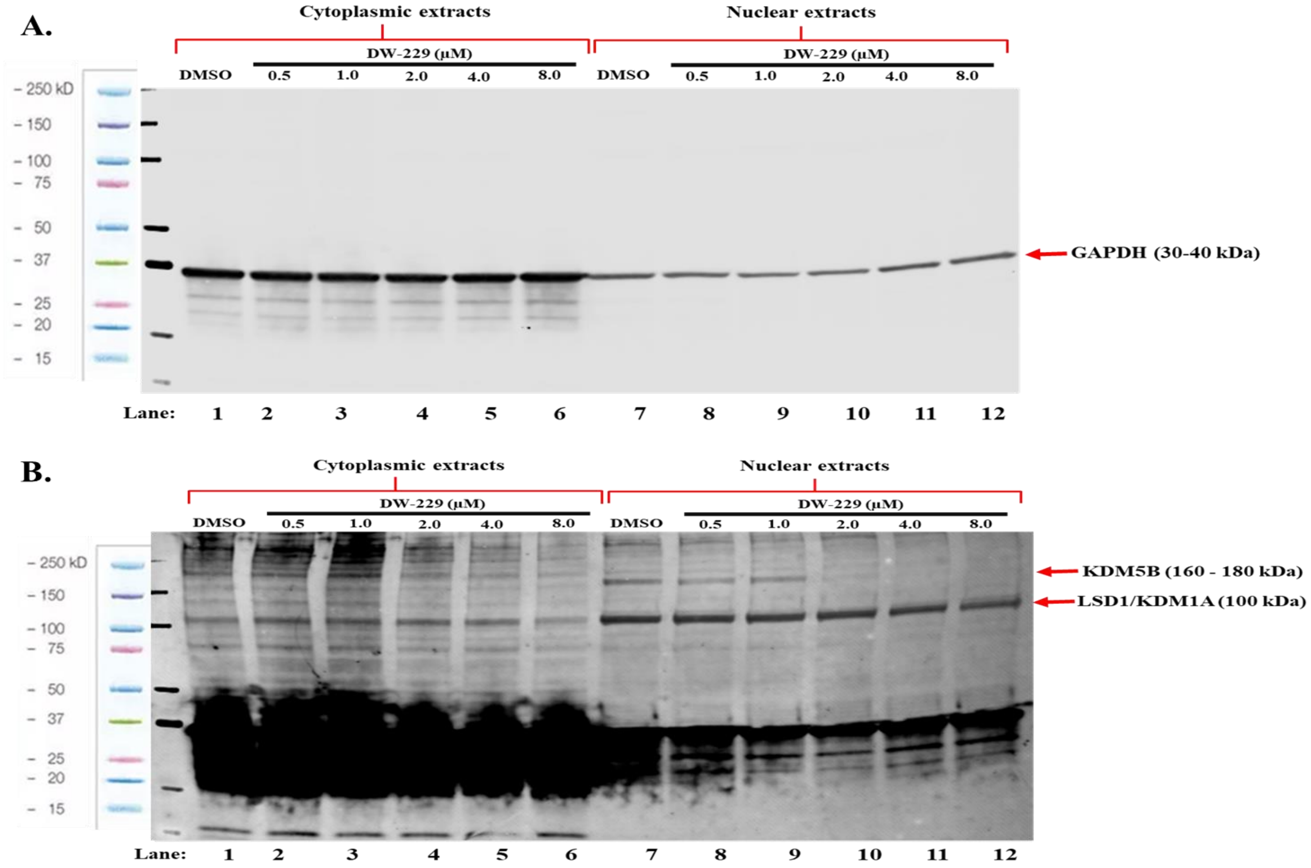


**Figure S24.** Full immunoblot gel images (A-B) of MCF-7 cells treated with **DW-229** for 24 h: *KDM5B antibody cocktail.*

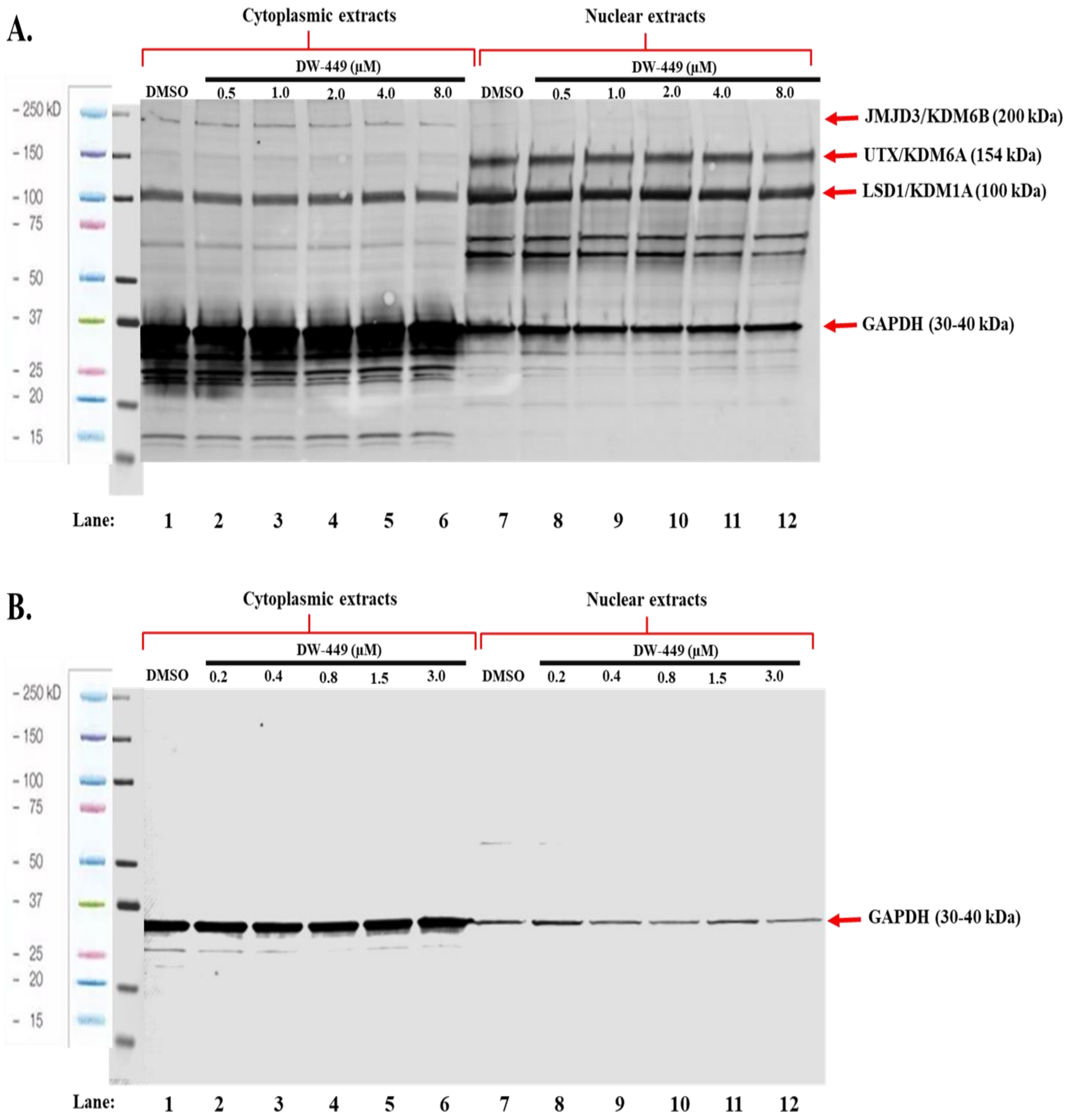


**Figure S25.** Full immunoblot gel images (A-B) of MCF-7 cells treated with **DW-449** for 24 h: *KDM1A, KDM6A, and KDM6B antibody cocktail.*

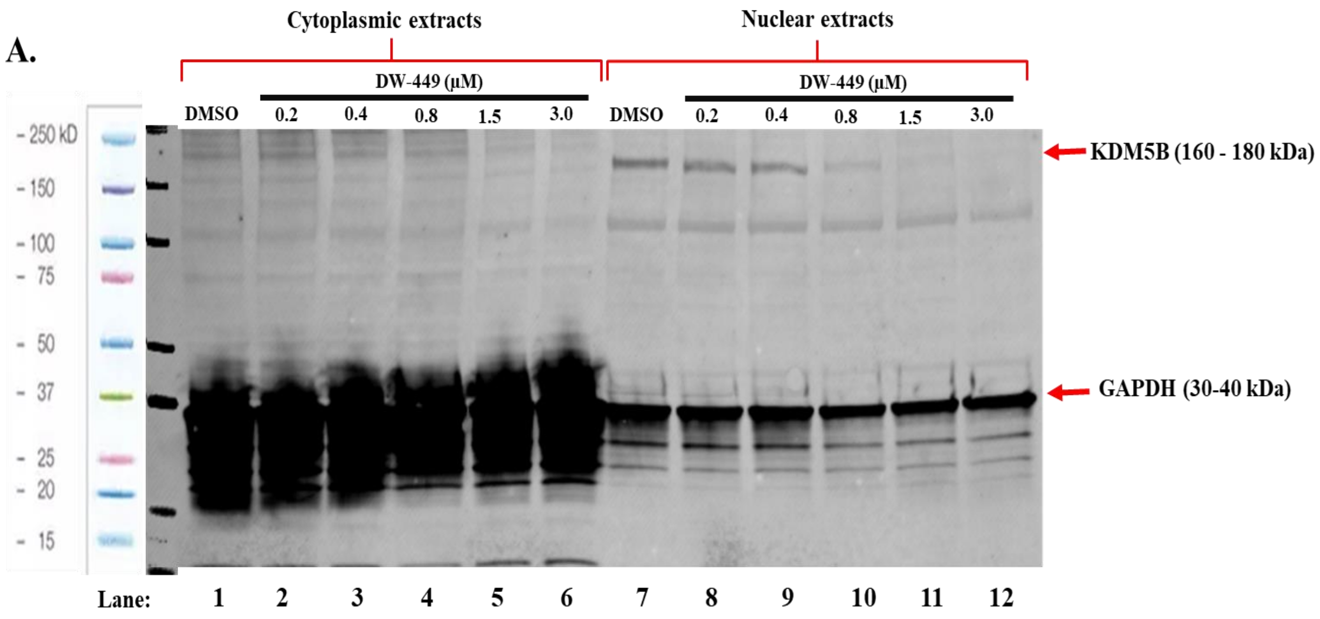


**Figure S26.** Full immunoblot gel images (A-B) of MCF-7 cells treated with **DW-449** for 24 h: *KDM5B antibody cocktail.*

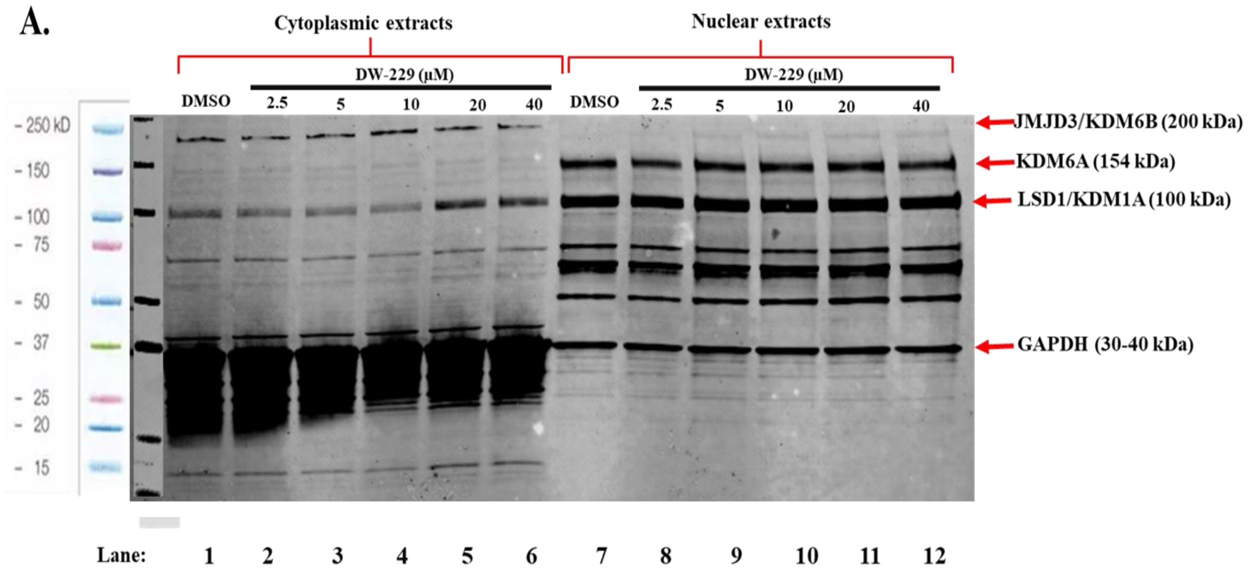


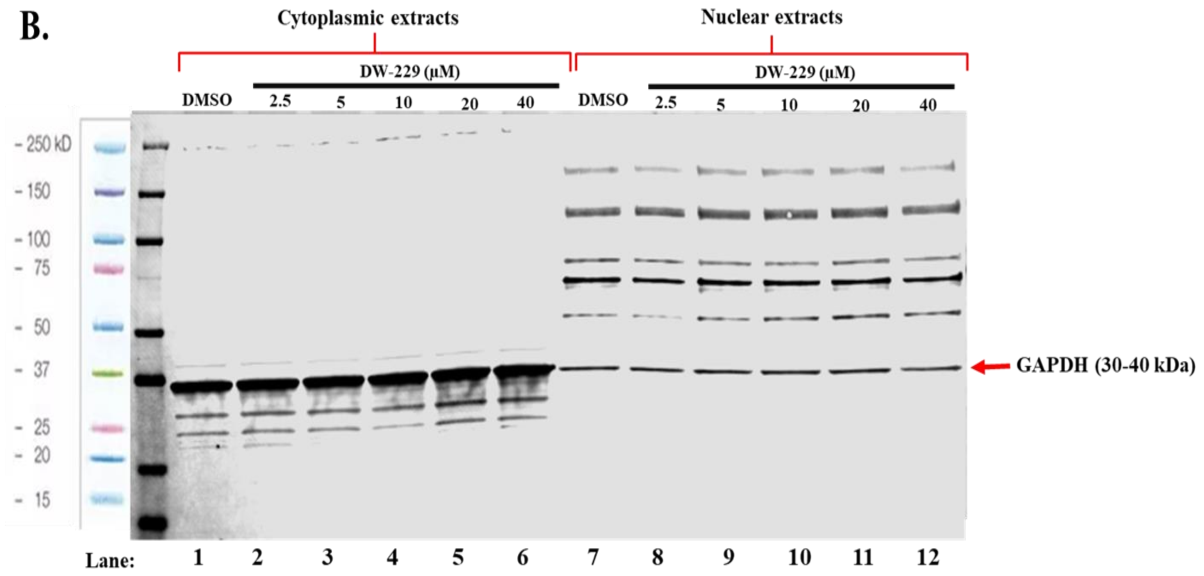


**Figure S27.** Full immunoblot gel images (A-B) of MDA-MB-231 cells treated with **DW-229** for 24 h: *KDM1A, KDM6A, and KDM6B antibody cocktail.*

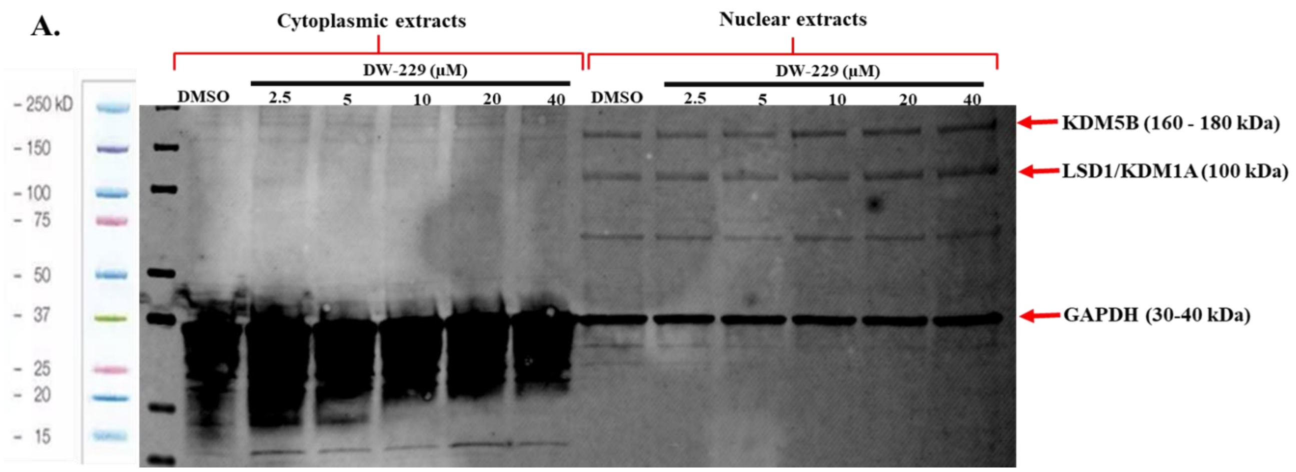


**Figure S28.** Full immunoblot gel image of MDA-MB-231 cells treated with **DW-229** for 24 h: *KDM5B antibody cocktail.*

A.

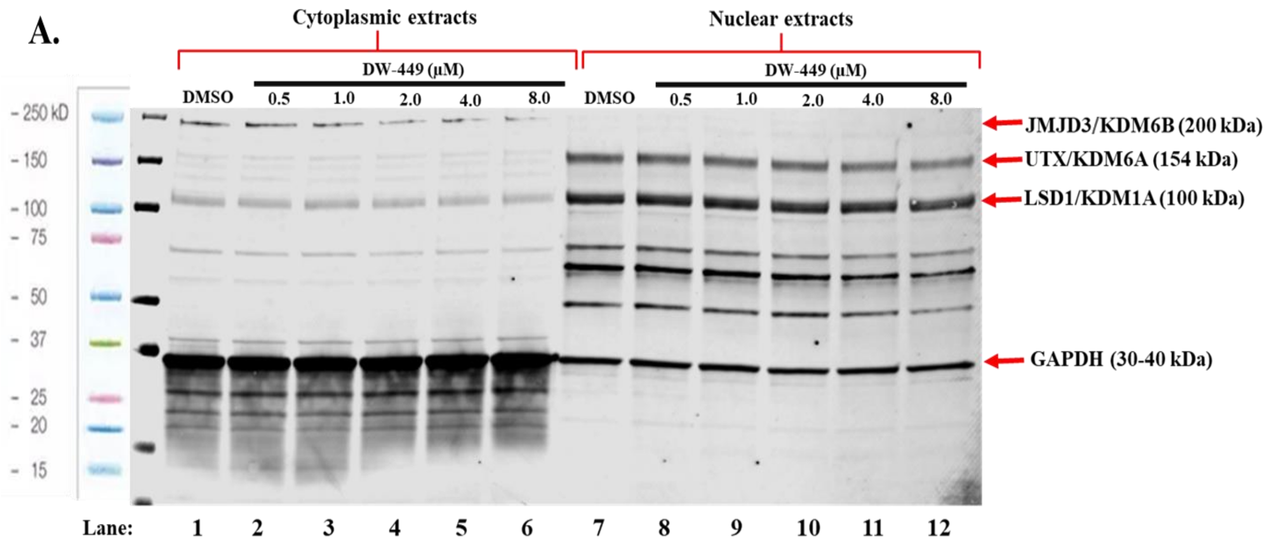


**Figure S29.** Full immunoblot gel image of MDA-MB-231 cells treated with **DW-449** for 24 h: *KDM1A, KDM6A, and KDM6B antibody cocktail.*

A.

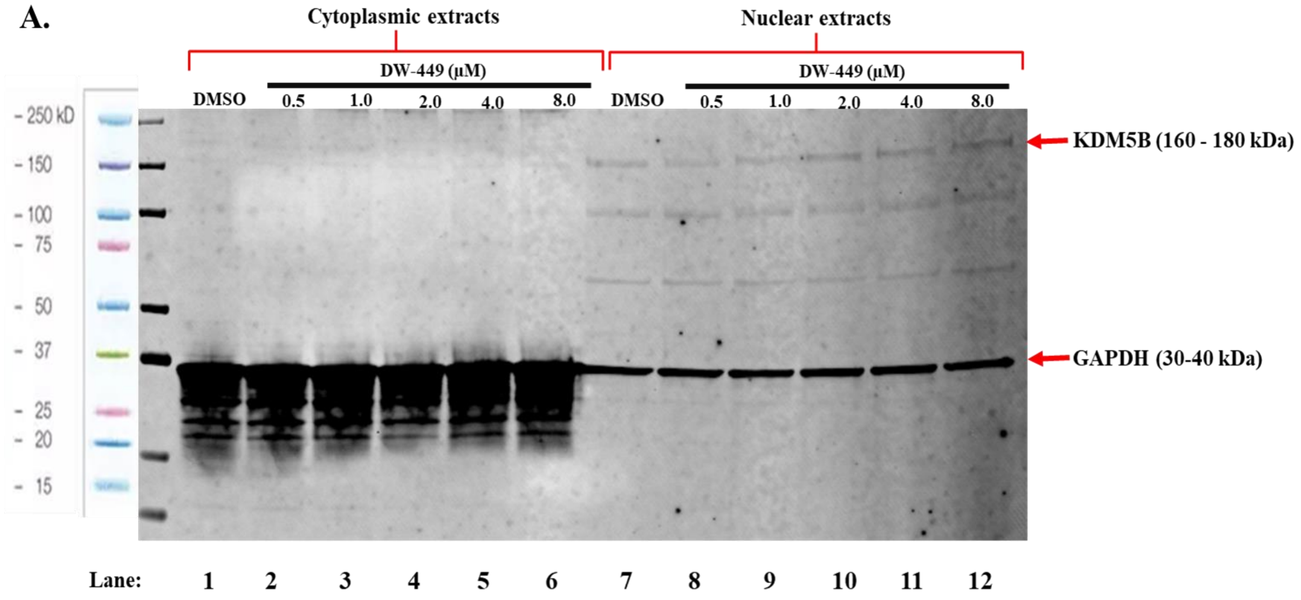


**Figure S30.** Full immunoblot gel image of MDA-MB-231 cells treated with **DW-449** for 24 h: *KDM5B antibody cocktail.*

**Table S3.** Effects of **DW-229** relative to DFP on KDM expression in MCF-7 cells. Log2 fold change and p-values highlighting the high significance of KDM2A (p = 0.011, **DW-229**/DFP 2x $IC_{50}$), KDM3A (p = 2.5e-02, **DW-229**/DFP $IC_{50}$, p = 9.4e-04, **DW-229**/DFP 2x $IC_{50}$), and KDM5B (p = 1.8e-04, **DW-229**/DFP $IC_{50}$, p = 2.9e-07, **DW-229**/DFP 2x $IC_{50}$). ns = not significant.

| | **DW229/DFP IC50** | | **DW229/DFP 2xIC50** | |
|---|---|---|---|---|
| | **log2 fold change** | **p value** | **log2 fold change** | **p value** |
| KDM1A | -0.3 | ns | -0.2 | ns |
| KDM1B | 0.2 | ns | 0.3 | ns |
| KDM2A | -0.5 | ns | -0.8 | 1.1e-02 |
| KDM2B | -0.2 | ns | -0.2 | ns |
| KDM3A | -1.0 | 2.5e-02 | -1.5 | 9.4e-04 |
| KDM3B | -0.2 | ns | -0.2 | ns |
| KDM4A | -0.3 | ns | -0.5 | ns |
| KDM4B | -0.2 | ns | -0.5 | ns |
| KDM4C | -0.1 | ns | -0.4 | ns |
| KDM4D | 0.0 | ns | 0.1 | ns |
| KDM4E | 0.0 | ns | 0.0 | ns |
| KDM5A | -0.1 | ns | -0.1 | ns |
| KDM5B | -1.2 | 1.8e-04 | -1.7 | 2.9e-07 |
| KDM5C | -0.4 | ns | -0.5 | ns |
| KDM5D | 0.1 | ns | 0.2 | ns |
| KDM6A | -0.1 | ns | -0.1 | ns |
| KDM6B | -0.2 | ns | -0.5 | ns |

| | | | | | |
|---|---|---|---|---|---|
| UTY | 0.1 | ns | | 0.1 | ns |
| KDM7A | 0.0 | ns | | 0.2 | ns |
| PHF8 | -0.1 | ns | | -0.3 | ns |
| PHF2 | -0.1 | ns | | -0.2 | ns |
| KDM8 | 0.0 | ns | | 0.0 | ns |
| TET1 | 0.0 | ns | | 0.0 | ns |
| TET2 | -0.7 | ns | | -0.4 | ns |
| TET3 | -0.3 | ns | | -0.5 | ns |

#### NMR Spectra

#### [1]HNMR Spectra compound 2

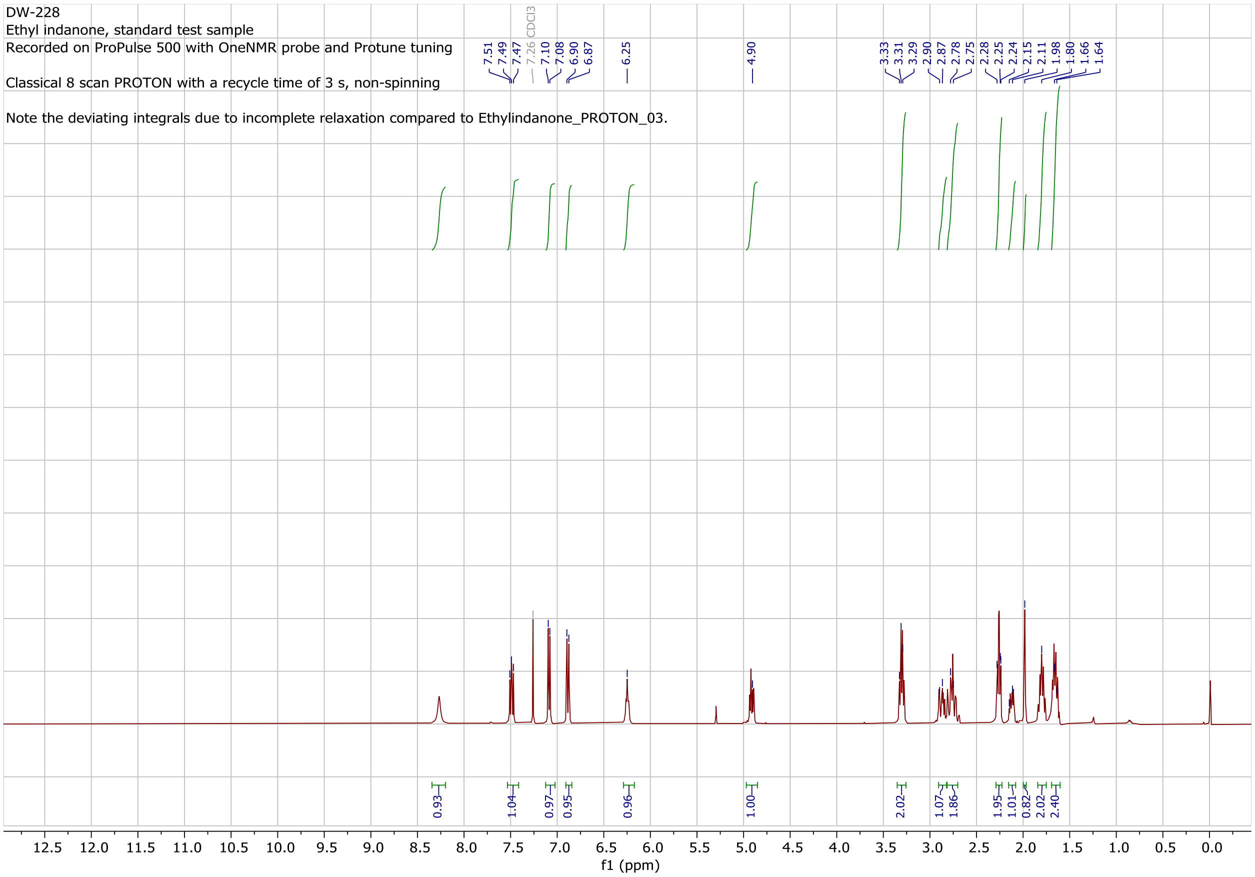

#### [1]HNMR Spectra compound 4a

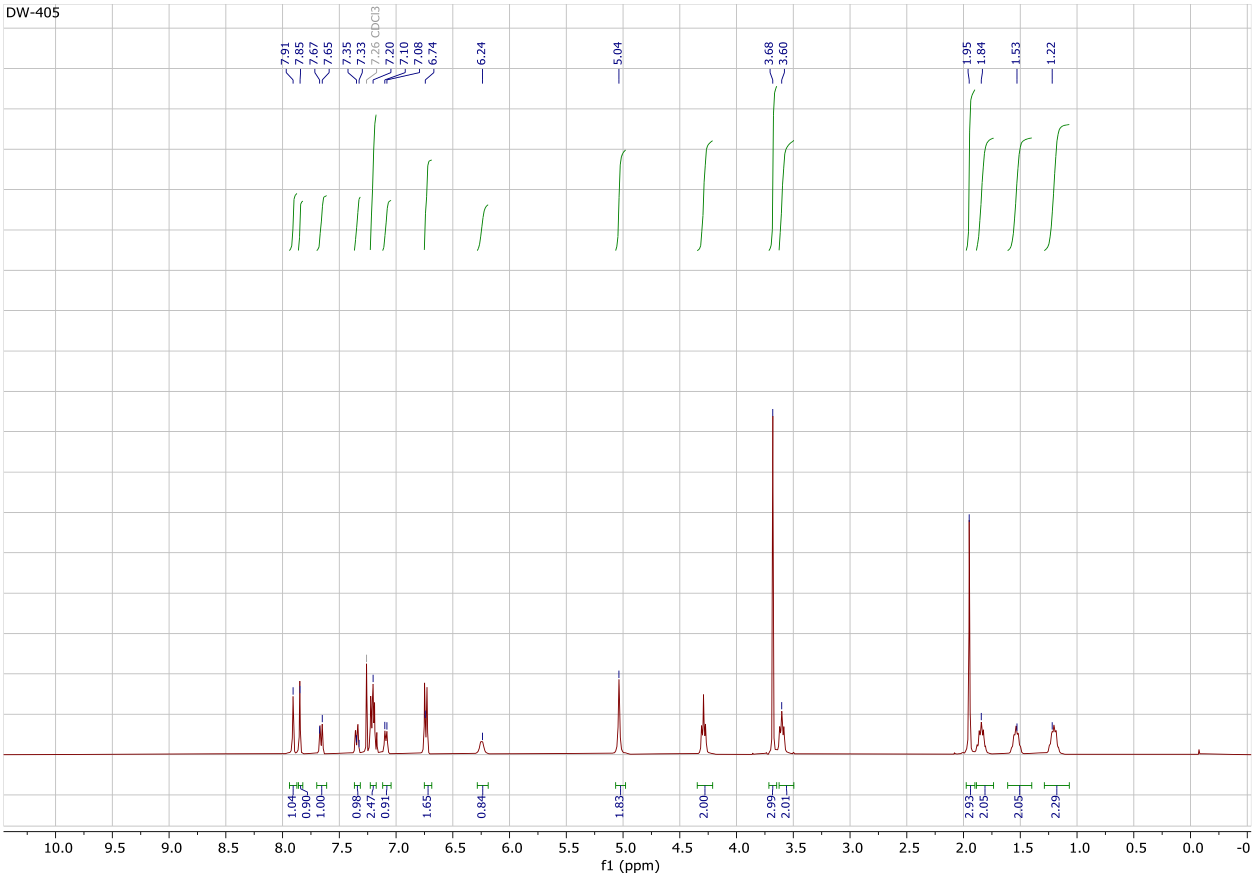

#### [1]HNMR Spectra compound 4b

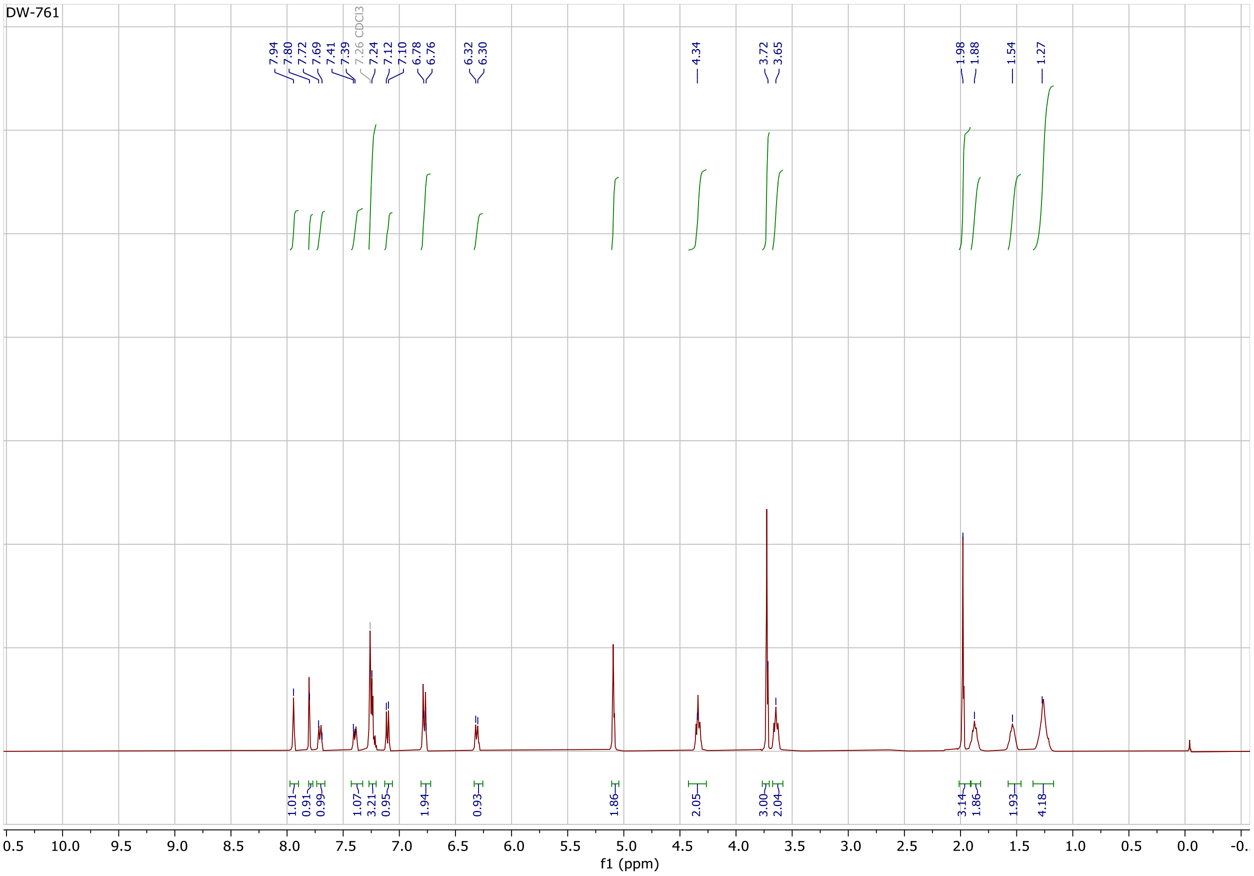

#### [1]HNMR Spectra compound 5a

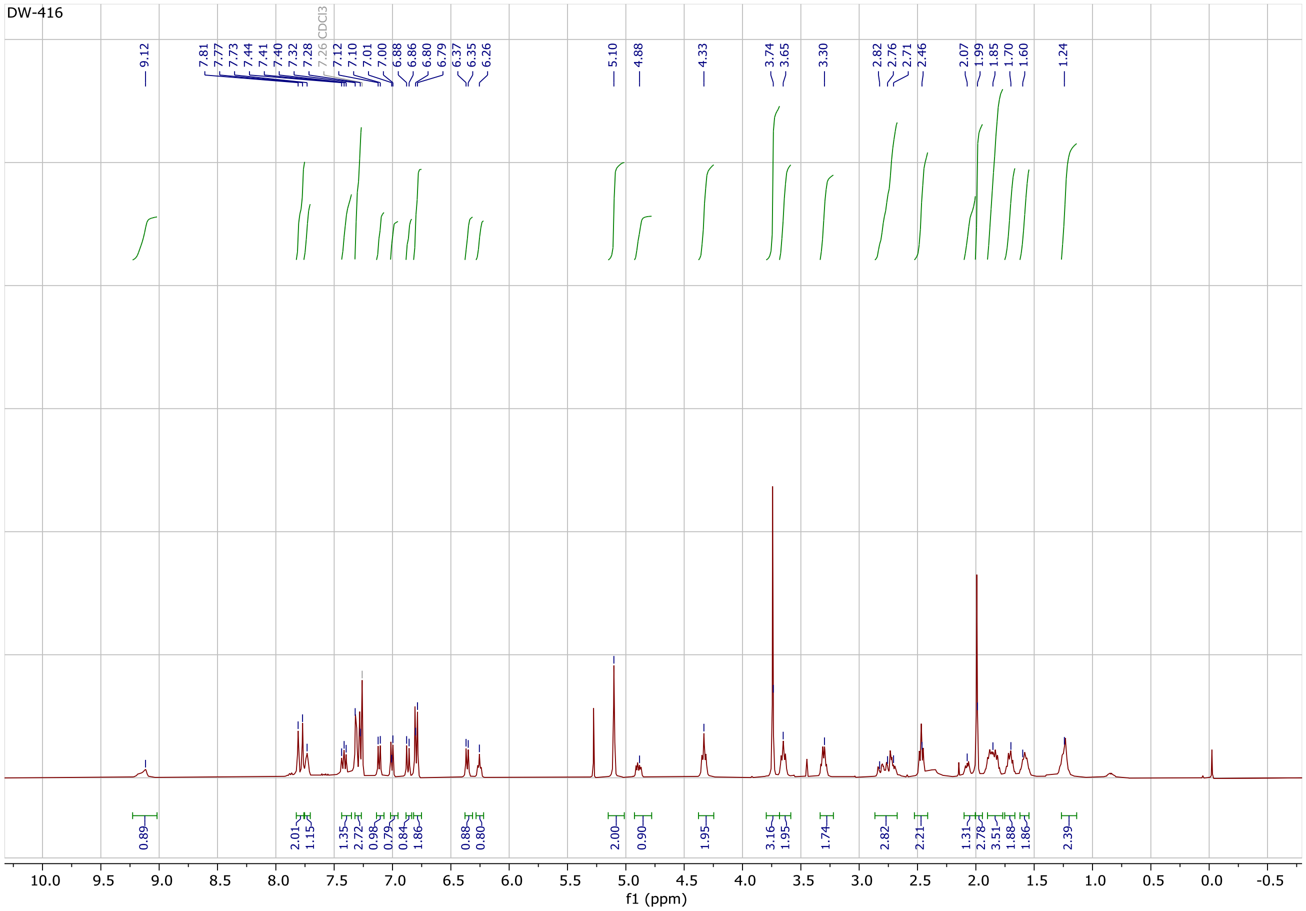

#### [1]HNMR Spectra compound 5b

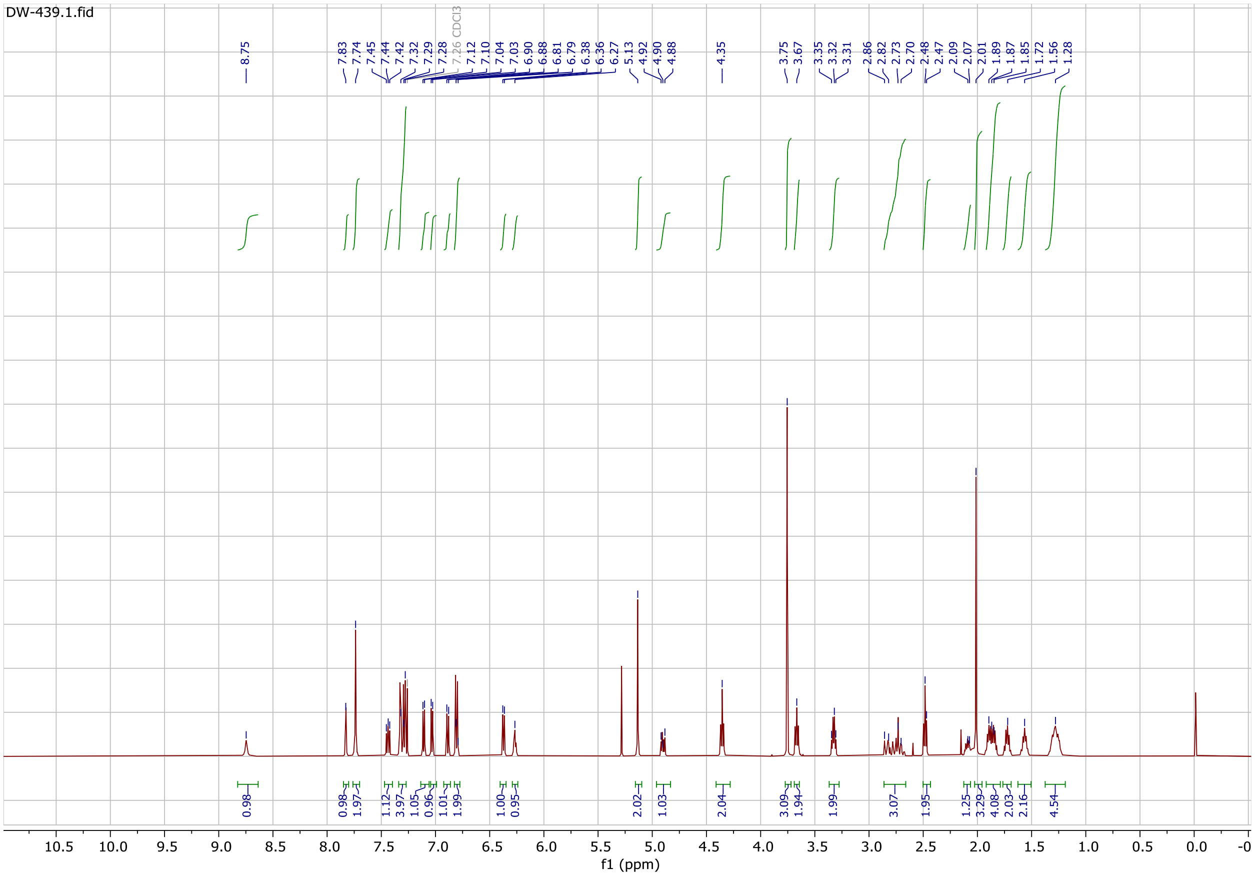

### $^{1}$H and $^{13}$ C NMR Spectra of compound DW-449

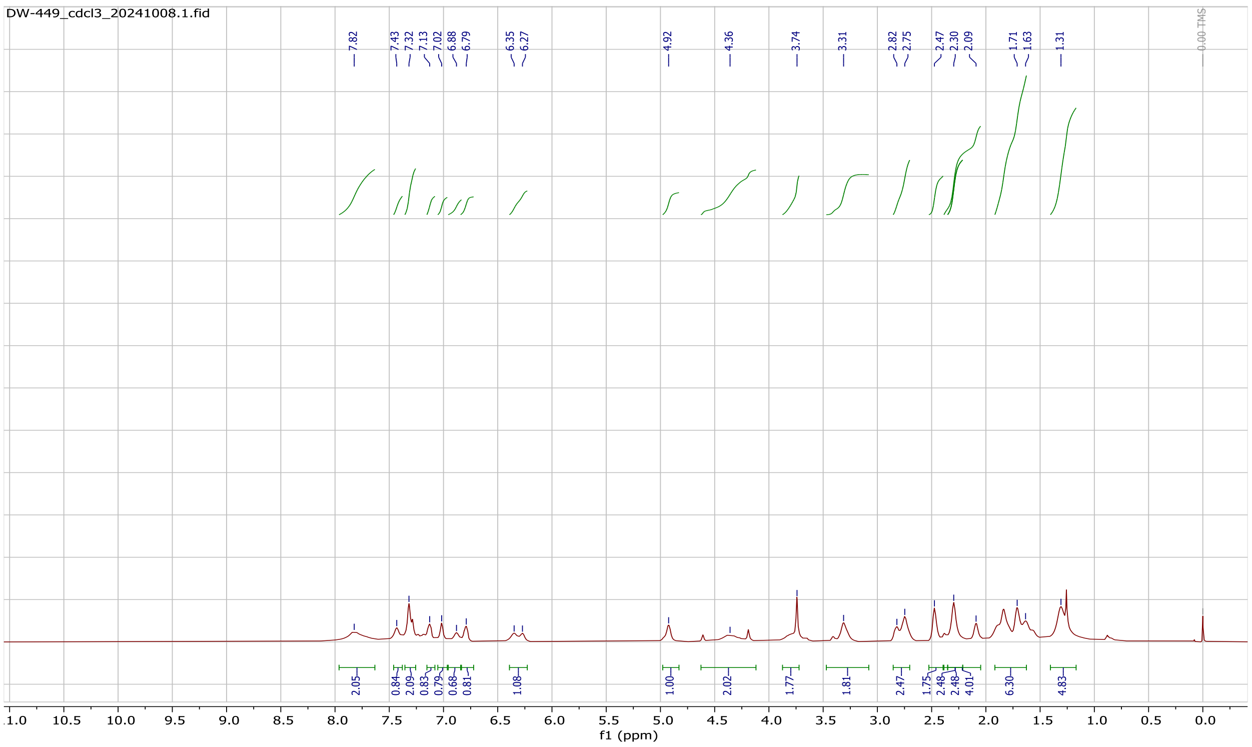


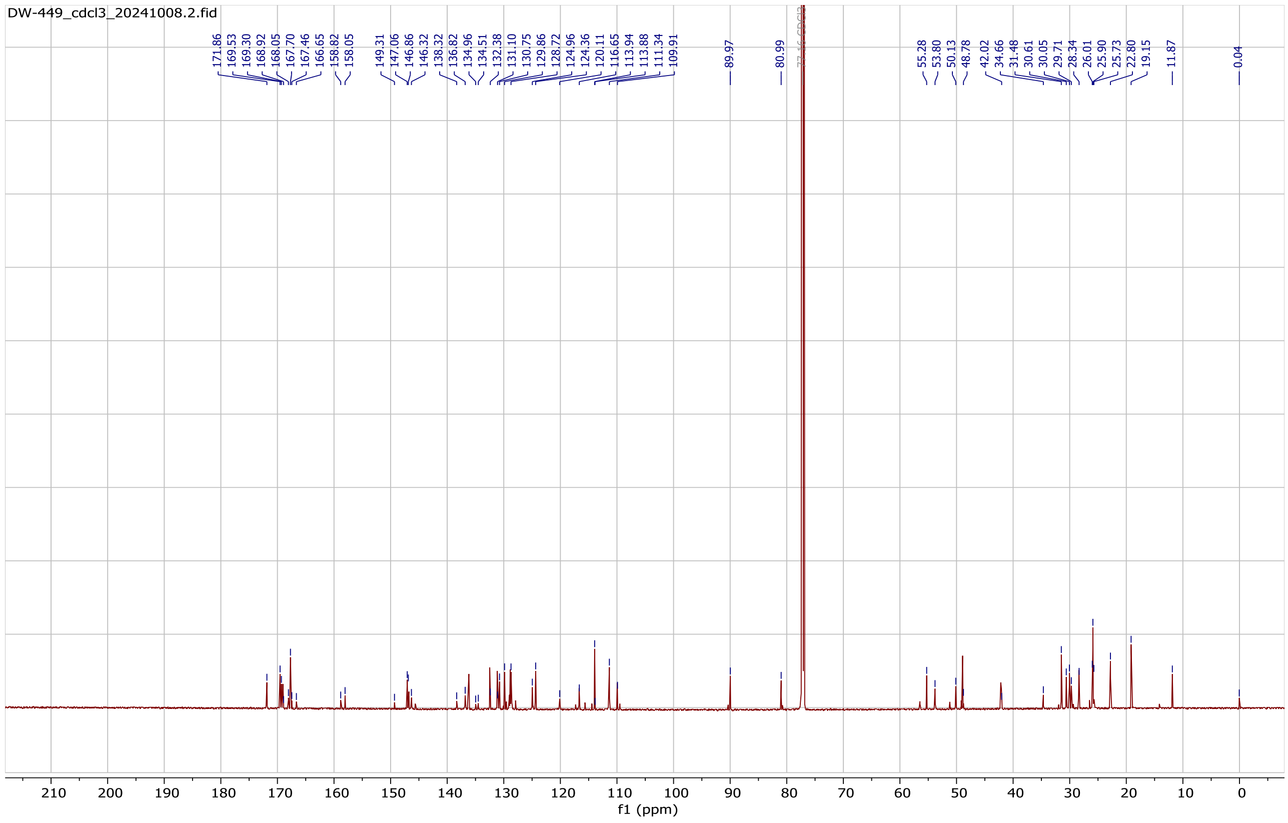

**[1]H and [13] CNMR Spectra compound 455**

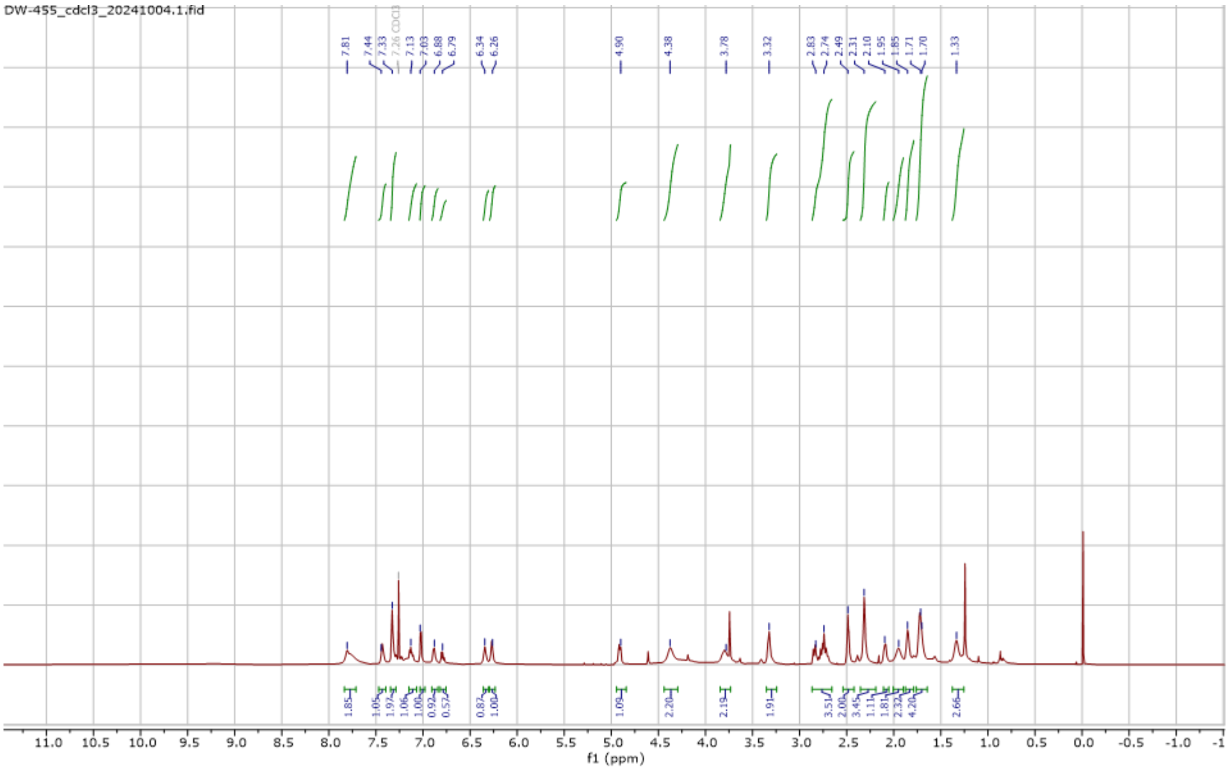

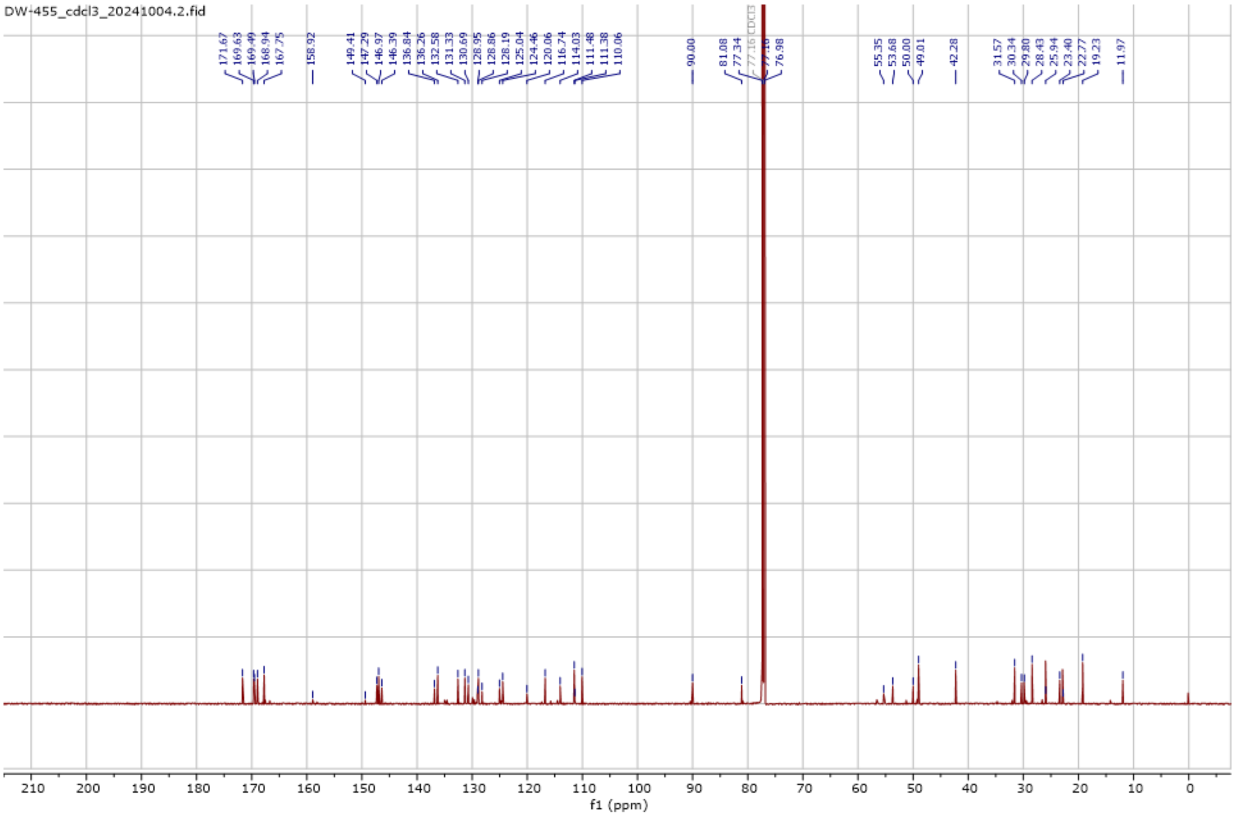

### [1]HNMR Spectra compound 6a

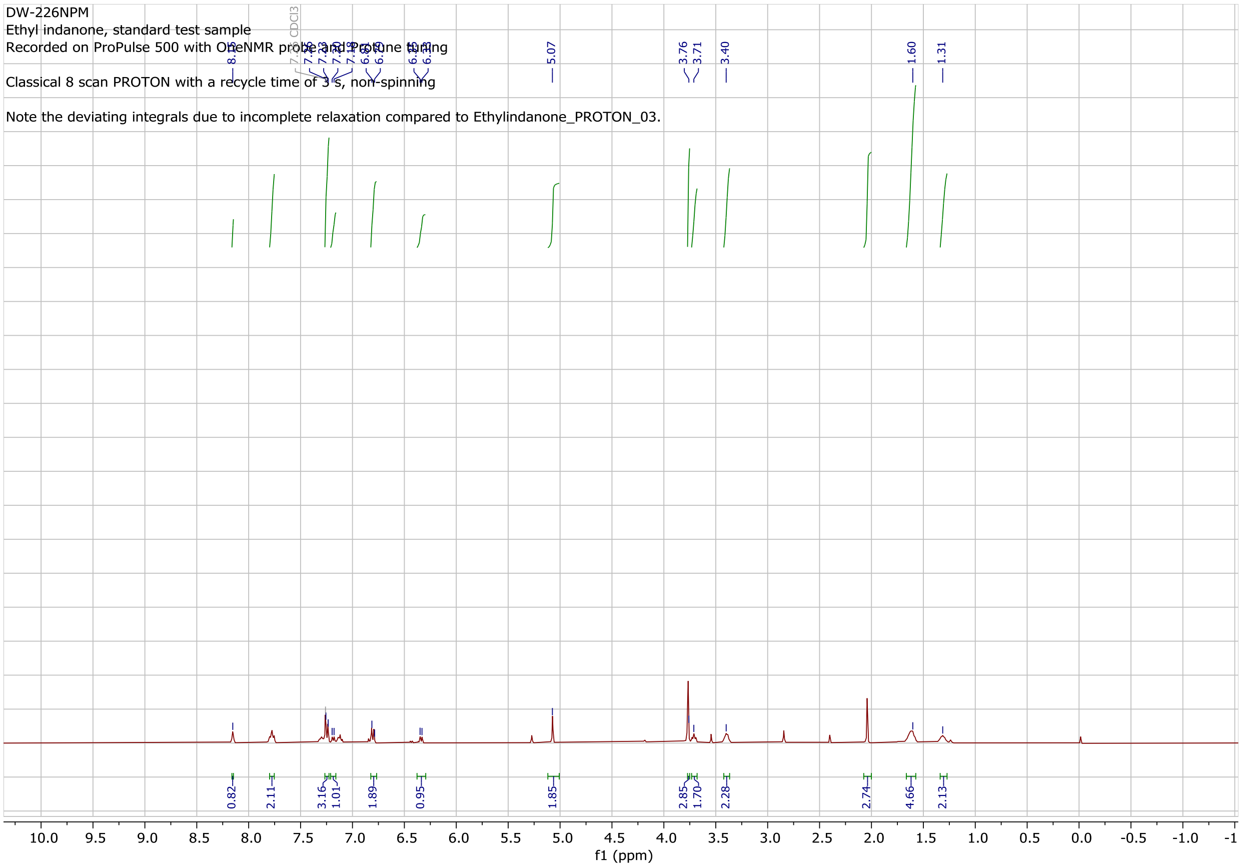

### [1]HNMR Spectra compound 6b

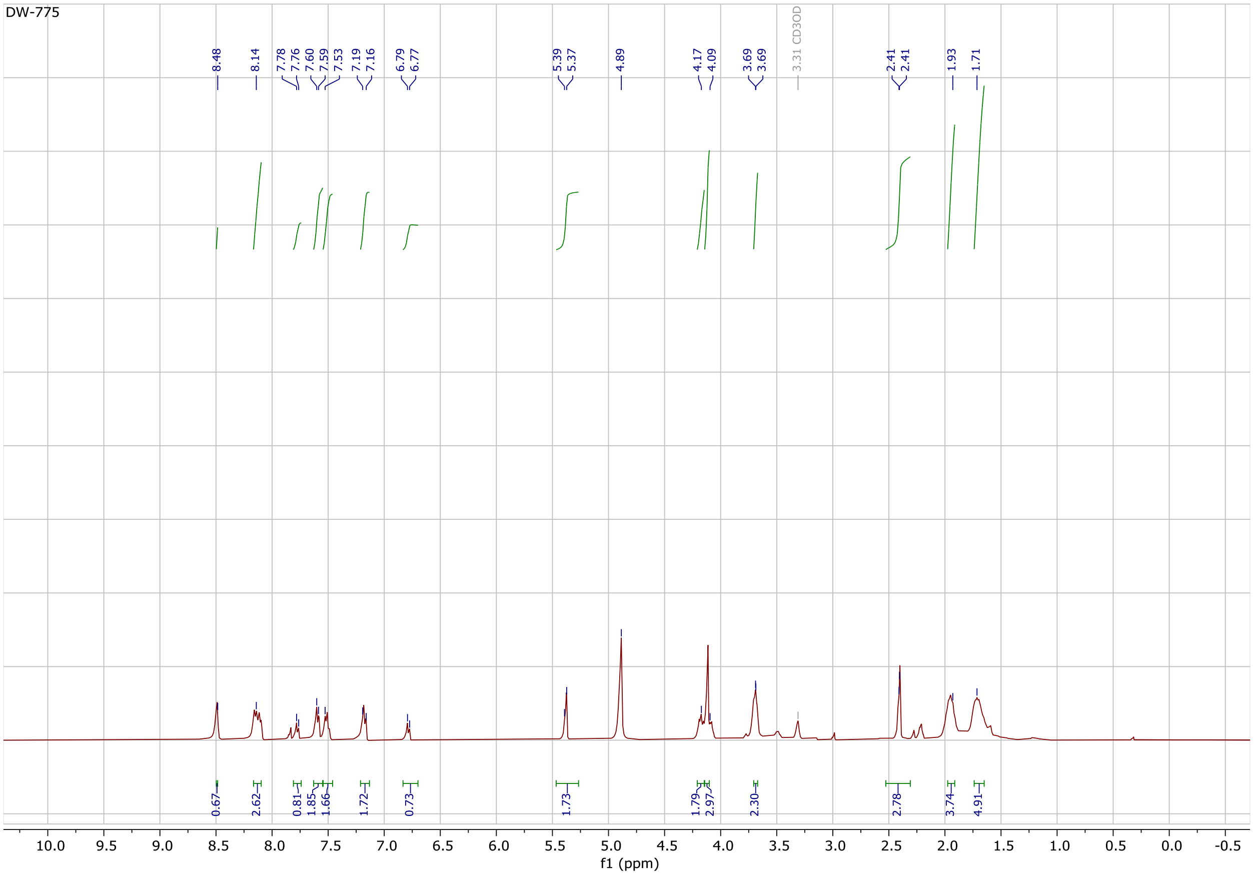

#### [1]HNMR Spectra compound 7a

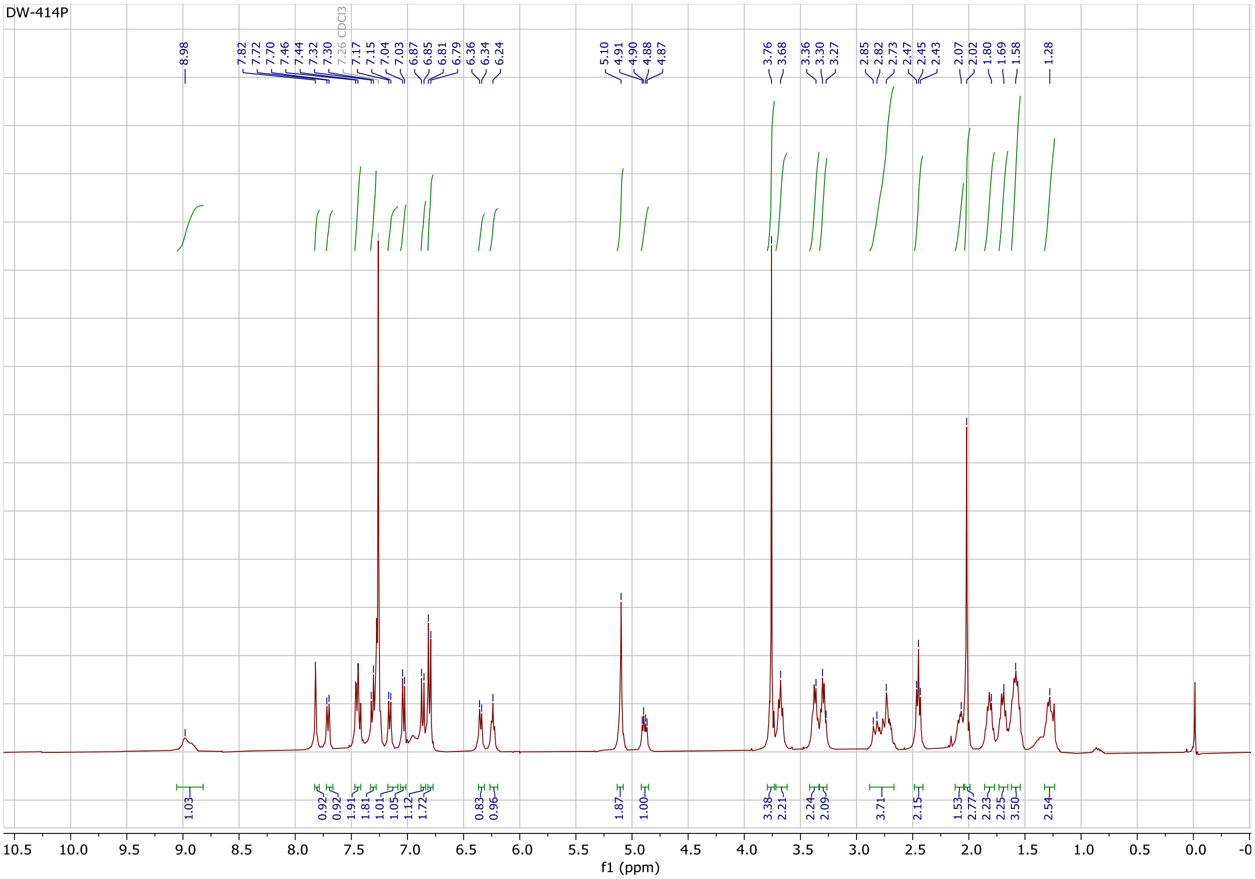

### [1]HNMR Spectra compound 7b

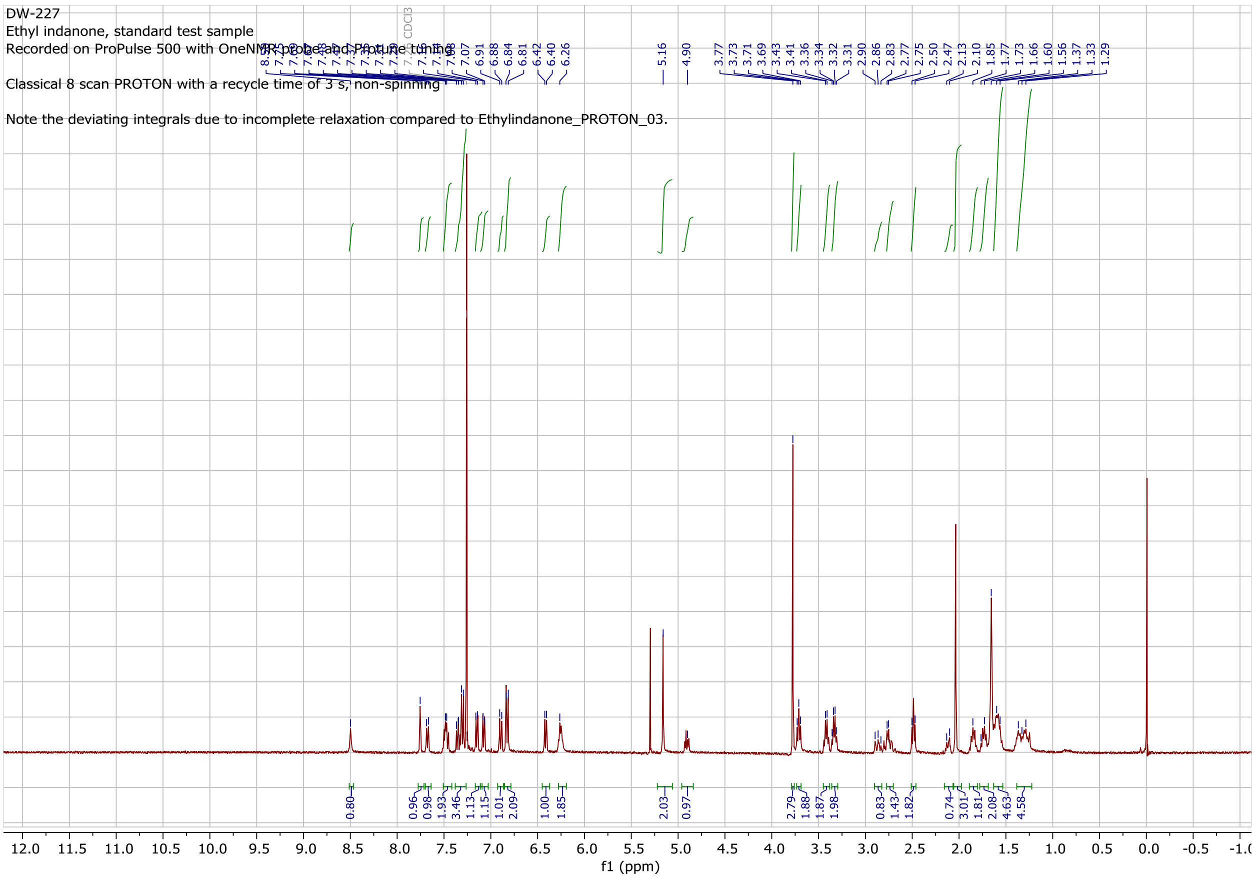

#### $^{1}H$ and $^{13}C$ NMR Spectra compound DW-451

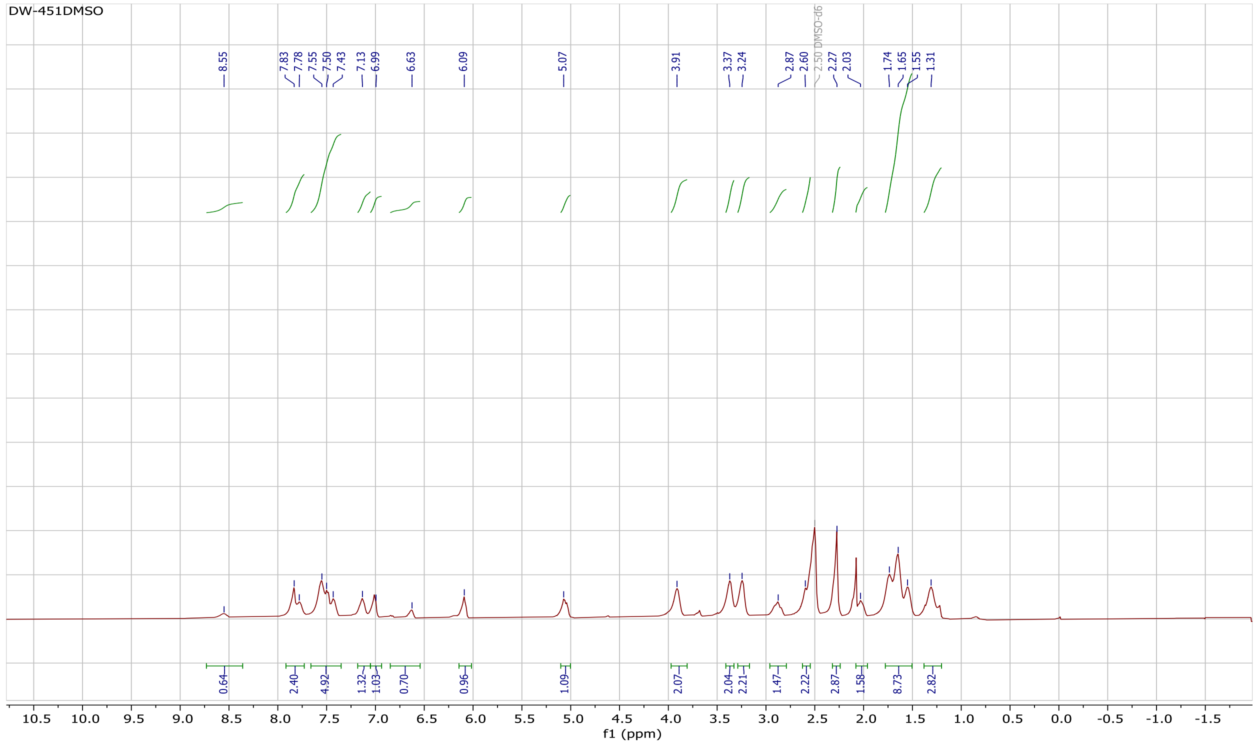

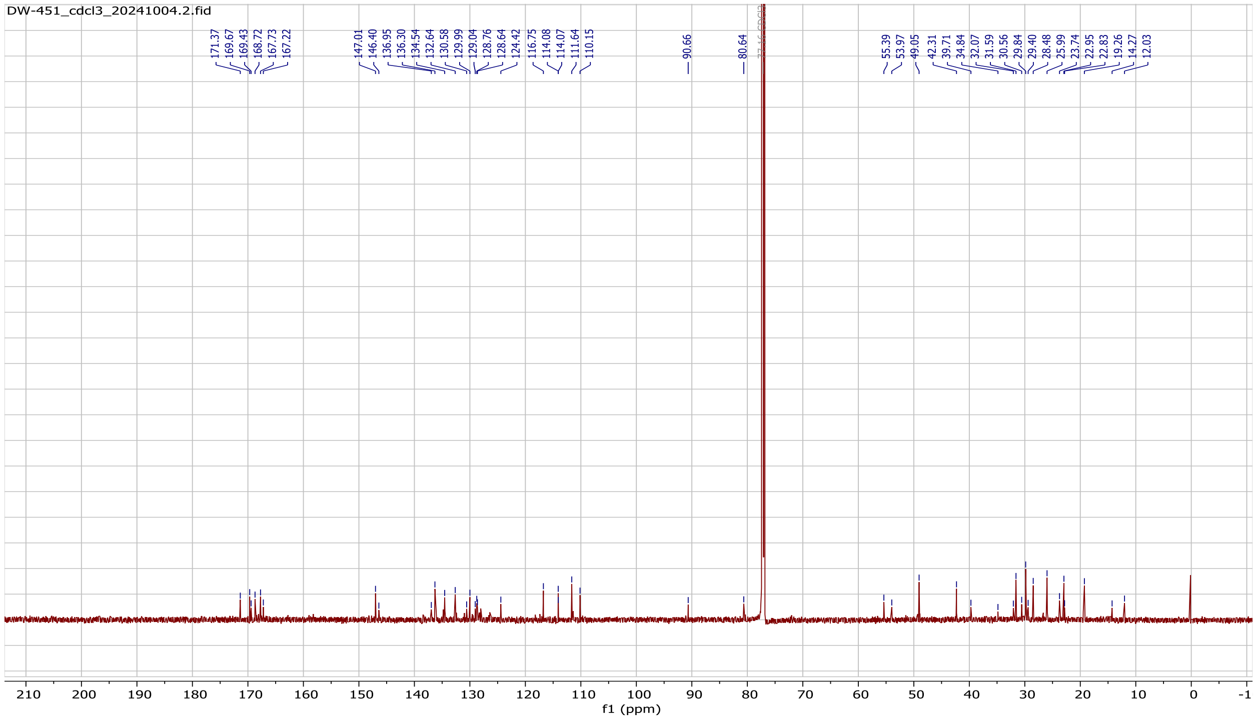

**$^{1}$H and $^{13}$ C NMR Spectra compound DW-229compound Purity and UV peak at 254 nm of**

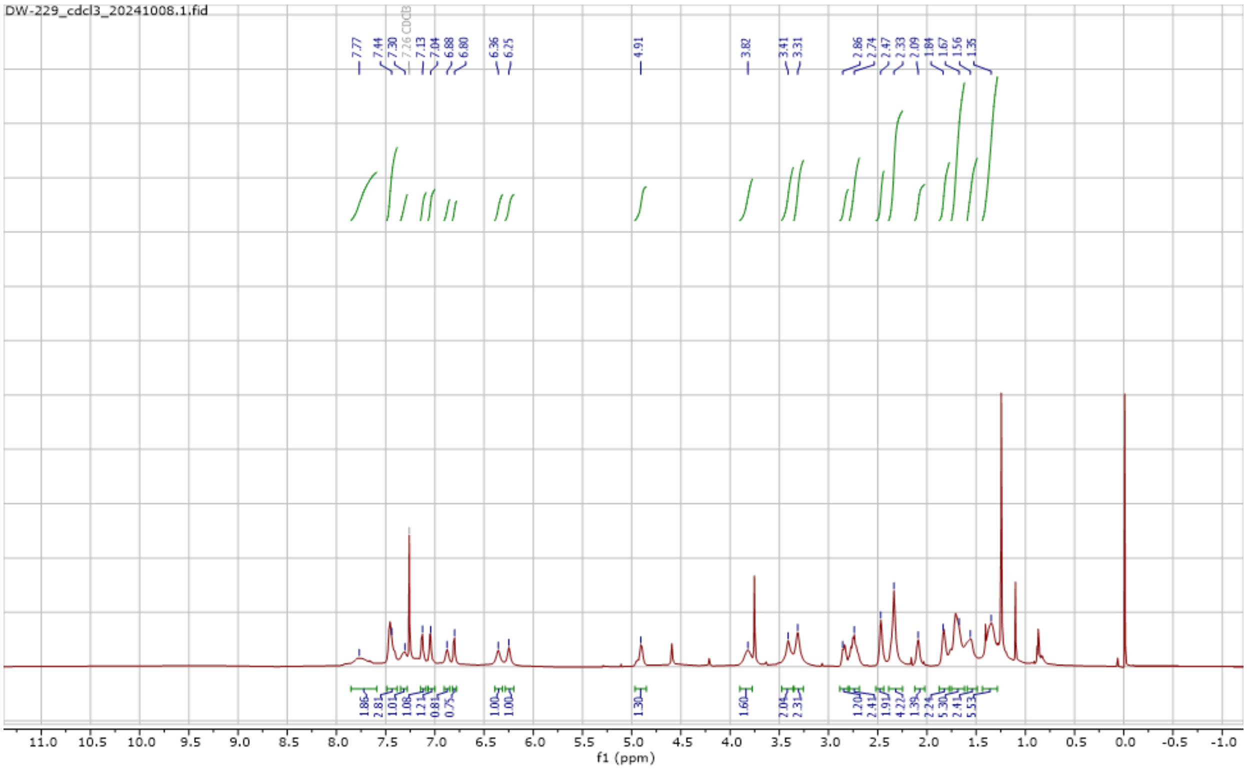

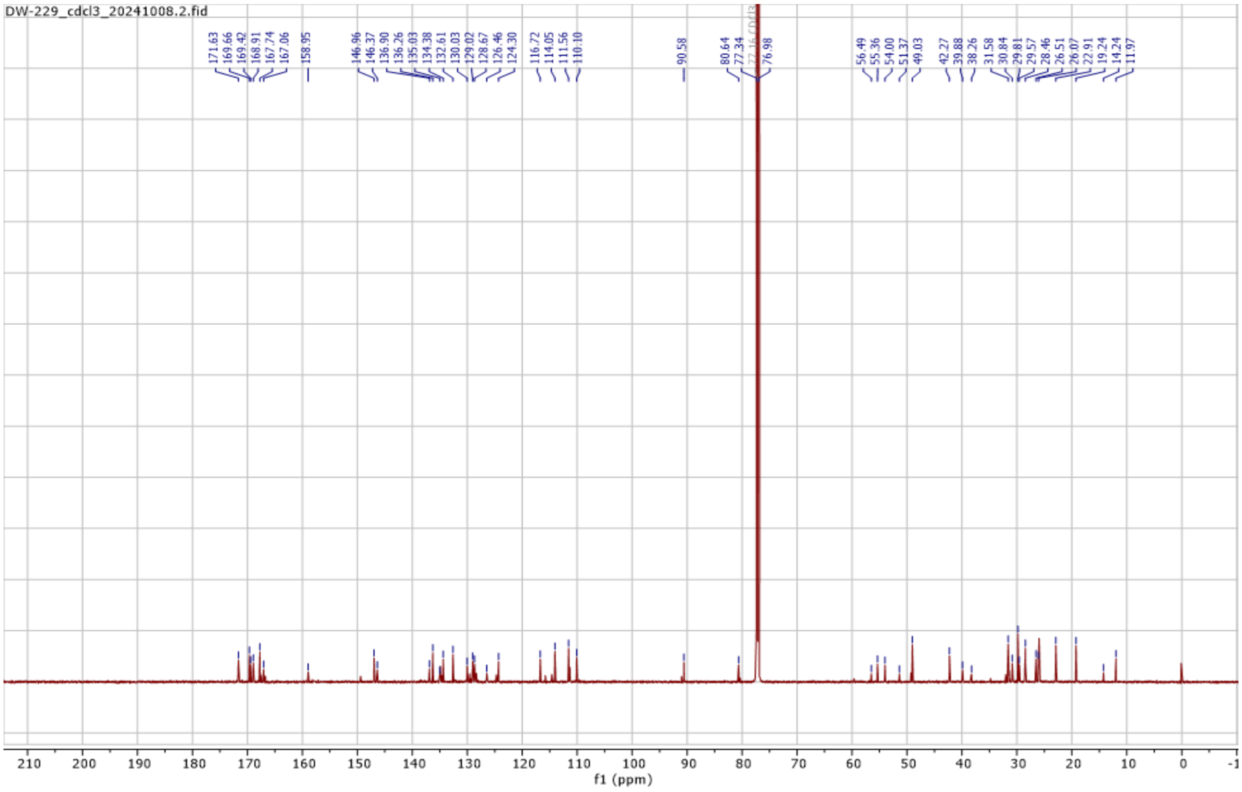

#### Compound DW-229

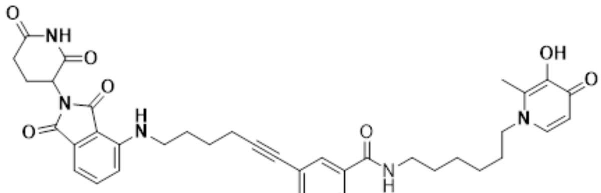


DW-229

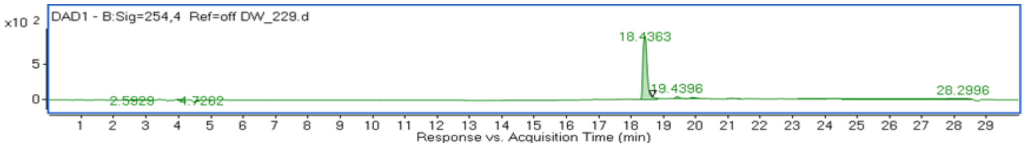


Peaks: + EIC(680.0958-684.6608) Scan

| Peak | RT | Area | Height |
|---|---|---|---|
| 1 | 18.5243 | 3379813411.16 | 168607720.36 |
| 2 | 19.5503 | 1117859122.14 | 53871845.43 |
| 4 | 21.7738 | 120032781.29 | 2271651.26 |
| 3 | 21.5001 | 49990795.16 | 2549714.22 |
| 6 | 23.9973 | 37288799.55 | 2100027.75 |
| 5 | 23.6553 | 36925788.64 | 1864098.51 |

**Purity and UV peak at 254 nm of compound DW-449**

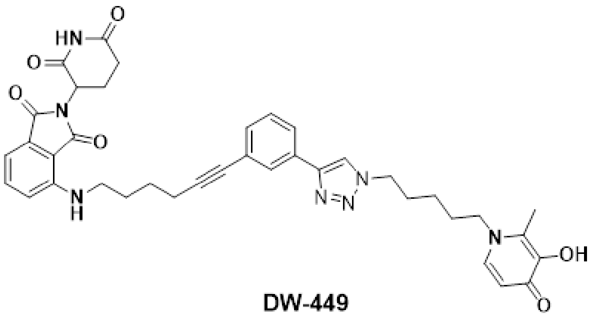

DW-449

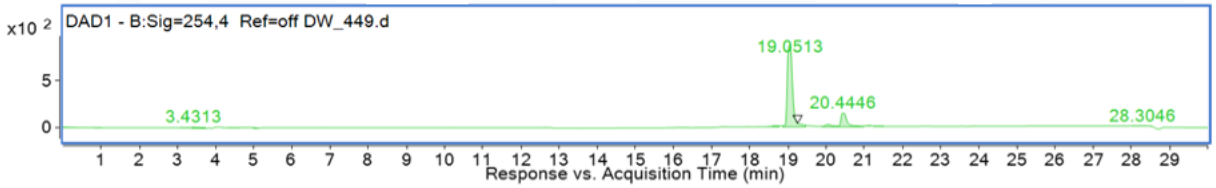


Peaks: + EIC(703.9774-708.4478) Scan

| Peak | RT | Area | Height |
|---|---|---|---|
| 1 | 19.1617 | 2411333887.96 | 139798601.42 |
| 2 | 20.1538 | 610623995.02 | 34884580.88 |
| 4 | 21.8642 | 86726487.94 | 1824763.52 |
| 6 | 23.6428 | 57988427.46 | 1617430.21 |
| 3 | 21.488 | 52909962.75 | 2178642.23 |
| 5 | 22.7536 | 34819478.69 | 1742042.86 |

#### Purity and UV peak at 254 nm of compound DW-451

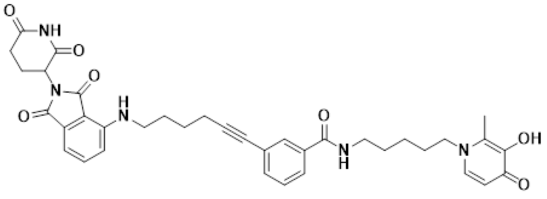

**DW-451**

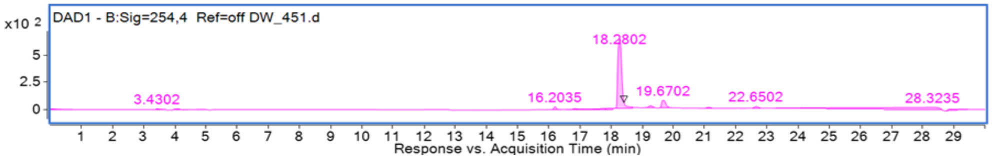


| Peak | RT | Area | Height |
|---|---|---|---|
| 2 | 18.3975 | 2612938983.33 | 149404464 |
| 3 | 19.355 | 451419919.81 | 25626396 |
| 1 | 18.0896 | 64918475.68 | 3495341 |
| 7 | 21.7836 | 48398046.27 | 1222699 |
| 5 | 20.7918 | 46923085.71 | 1720163 |
| 4 | 20.3814 | 43549586.11 | 2105039 |
| 6 | 21.4416 | 37858691.12 | 1391684 |
| 8 | 22.7414 | 32852667.56 | 1202977 |
| 9 | 23.8359 | 31038000.57 | 1135780 |

**Purity and UV peak at 254 nm of compound DW-455**

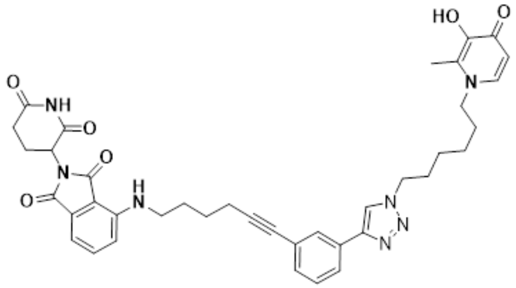

**DW-455**

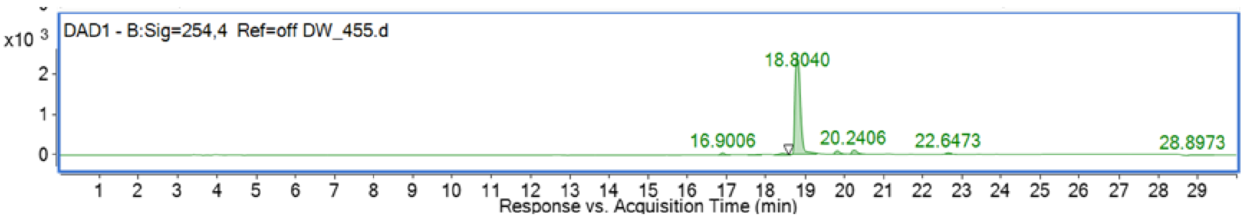


| Peak | RT | Area | Height | Typ |
|---|---|---|---|---|
| 2 | 18.9238 | 3259521811.32 | 151964463.1 | |
| 3 | 19.9158 | 1123848715.62 | 61894715.63 | |
| 4 | 21.3526 | 118809808.28 | 2676815.99 | |
| 1 | 18.5818 | 80136626.52 | 6725016.26 | |
| 6 | 22.7892 | 69001198.22 | 2427805.06 | |
| 7 | 23.6101 | 68278078.43 | 2206780.41 | |
| 5 | 22.3445 | 40611030.69 | 1968300.39 | |
